# Genomics and transcriptomics to unravel sex determination pathway and its evolution in sand flies

**DOI:** 10.1101/510586

**Authors:** Valeria Petrella, Serena Aceto, Vincenza Colonna, Giuseppe Saccone, Remo Sanges, Nikola Polanska, Petr Volf, Luigi Gradoni, Gioia Bongiorno, Marco Salvemini

## Abstract

**Background:** Phlebotomine sand flies (Diptera, Nematocera) are important vectors of several pathogens, including *Leishmania* parasites, causing serious diseases of humans and dogs. Despite their importance as disease vectors, most aspects of sand fly biology remain unknown including the molecular bases of their reproduction and sex determination, aspects also relevant for the development of novel vector control strategies.

**Results:** Using a comparative genomics/transcriptomics approach, we identified the sex determining genes in phlebotomine sand flies and proposed the first model for the sex determination cascade of these insects. For all the genes identified, we produced manually curated gene models, developmental gene expression profile and performed evolutionary molecular analysis. We identified and characterized, for the first time in a Nematocera species, the *transformer* (*tra*) homolog which exhibits both conserved and novel features. The analysis of the *tra* locus in sand flies and its expression pattern suggest that this gene is able to autoregulate its own splicing, as observed in the fruit fly *Ceratitis capitata* and several other insect species.

**Conclusions:** Our results permit to fill the gap about sex determination in sand flies, contribute to a better understanding of this developmental pathway in Nematocera and open the way for the identification of sex determining orthologs in other species of this important Diptera sub-order. Furthermore, the sex determination genes identified in our work also provide the opportunity of future biotech applications to control natural population of sand flies, reducing their impact on public health.

## Background

In animals, sex determination is the process by which early embryos of metazoan species with sexual reproduction operate a binary decision between two conditions: male or female development. This key decision results in individuals that can be identified as males, females, or in some cases hermaphrodites and, in species with a genetic sex determination system, underlies genomic differences between sexes. In most cases, the presence of heteromorphic sexual chromosomes represents the primary signal for sex determination. According to the initial decision, the primary signal is then transduced, through a genetic pathway organized in a cascade of regulatory genes, to downstream regulators responsible for sexual differentiation [1–3].

Insects are among the largest taxonomic animal groups on Earth and, not surprisingly, they exhibit a wide variety of sex determining systems, with highly variable primary signals and widely conserved genetic transduction mechanisms to downstream regulators [4–6]. *Drosophila melanogaster* (Diptera, Drosophilidae) is the model species where sex determination is known at the higher level of molecular resolution (Fig. 1). In this species, sex determination is controlled by five main genes, *Sex-lethal* (*Sxl*), *transformer* (*tra*), *transformer-2* (*tra-2*), *doublesex* (*dsx*) and *fruitless* (*fru*), hierarchically organized in a regulative cascade: *Sxl* -> *tra*+*tra-2* -> *dsx, fru*. This cascade is activated by a primary signal represented by the number of X chromosomes [7,8]. In the last 20 years, homology-based approaches in species belonging to various insects orders (Diptera, Coleoptera, Lepidoptera, Hymenoptera) led to discover only partial conservation of the *Drosophila* sex determination genetic pathway: in all species studied the *Sxl* ortholog was not involved in sex determination while the *tra* ortholog is able to control the female-splicing of its own pre-mRNA as well as to control, similarly to *Drosophila*, the female-specific splicing of the *dsx* and *fru* downstream genes [6,9,10]. In female embryos, the maternal *tra* contribution establishes the female-specific autoregulatory splicing of *tra* and leads to female development, which is epigenetically maintained during development in the absence of the initial positive signal. In male embryos, the establishment of *tra* autoregulatory feedback loop is impaired by the presence of a masculinizing factor able to interfere with the maternal and/or the zygotic *tra* function, blocking its positive autoregulation and leading to male development, as shown recently in *Musca domestica* (Sharma et al., 2017). Hence, the *tra*+*tra2->dsx*/f*ru* sex determination module with an autoregulating *tra*, firstly discovered in the Mediterranean fruit fly *Ceratitis capitata* (Pane et al., 2002; Salvemini et al., 2009a), represents the core pathway of insect sex determination [13]. The only remarkable exception is represented by the Lepidoptera order, where a different sex determination system exists with the primary signal constituted by a small RNA, the absence of the *tra* ortholog and the *dsx* splicing controlled by different splicing regulators [14].

**Fig. 1.**
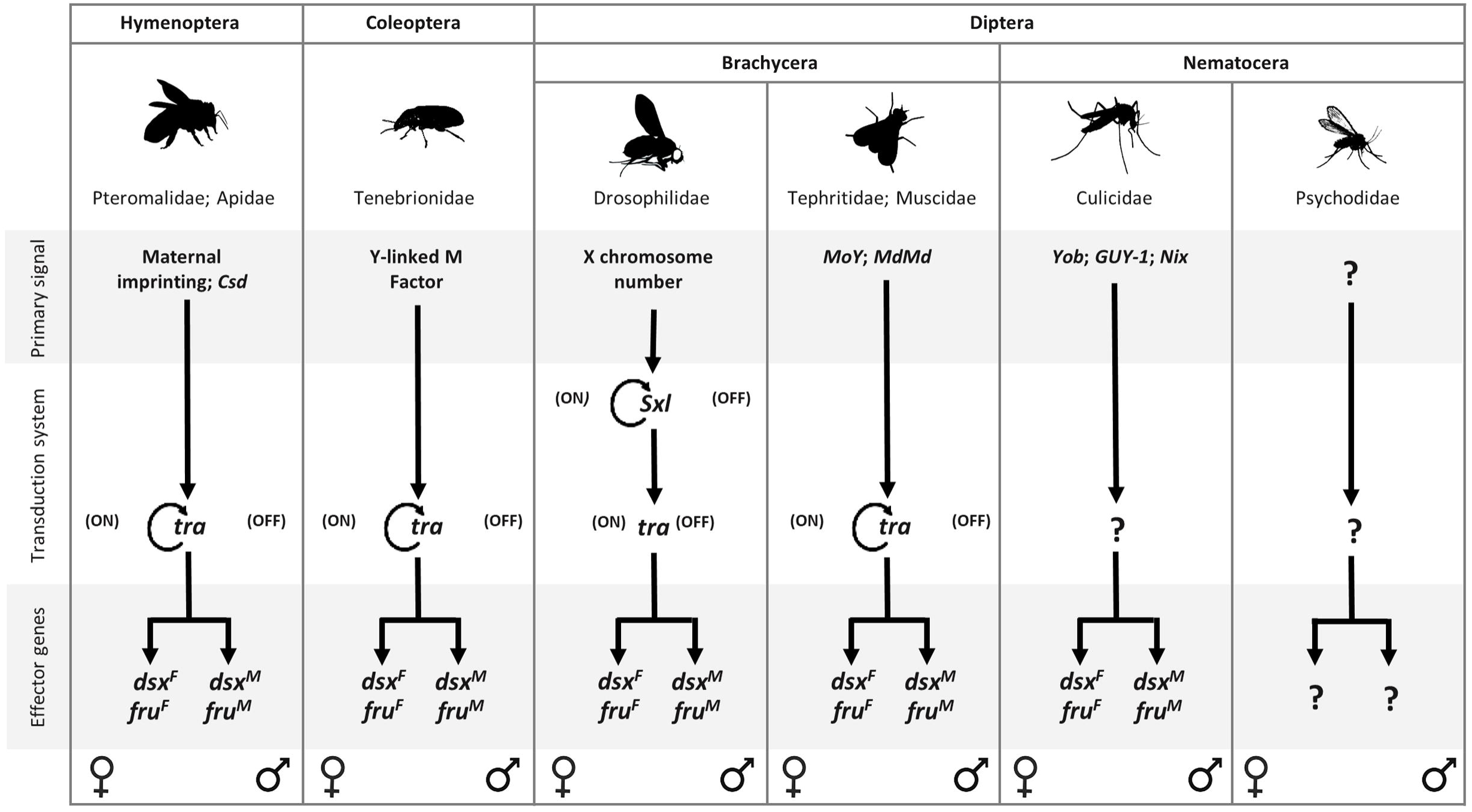
Sex determination in insect species. Orders, suborders and families of species with identified sex determining genes reported in figure are indicated. In the model system *Drosophila melanogaster,* the presence of two X chromosomes in the female embryo activates the *Sex-lethal* gene (*Sxl*) which, acting as a gene-specific splicing regulator, promotes the female-specific splicing of its own pre-mRNA and of the pre-mRNA of the gene downstream in the regulative cascade, *transformer* (*tra*). *tra* and the non-sex-specific auxiliary factor *transformer*-2 (*tra-2*) encode for splicing factors (TRA and TRA-2 proteins) able to control the splicing of at least two downstream target genes, responsible of sexual differentiation and courtshi pbehaviour: *doublesex* (*dsx*) and *fruitless* (*fru*), respectively. Both genes encode for sex-specific transcription factors that potentially binds to multiple genome loci, leading to sex-specific gene expression and subsequent sexual differentiation. In male embryo, the absence of the functional SXL protein leads to the male-specific splicing of *tra, dsx* and *fru* pre-mRNAs resulting in the activation of the male development program. In Hymenoptera, Coleoptera, and Diptera (Brachycera), different primary signals set the activity state of the *tra* homolog able to autoregulate its own splicing in the female sex and to determine female development. In mosquitoes (Diptera, Nematocera) *dsx* and *fru* genes exhibit, as for Brachycera species, a conserved alternative splicing regulation, producing sex-specific protein isoforms. Recently, genomic/transcriptomic studies of sex determination led to the discovery of novel primary signals including the Y-linked genes *Yob* and *Guy-1* in the malaria vectors *Anopheles gambiae* and *An. stephensi*, respectively, and the putative splicing factors *Nix* in the dengue vector *Aedes aegypti*. These primary signals are supposed to act upstream of *dsx* and *fru* genes in the sex determination cascade. However, their mechanism of action, direct or indirect, and the possible presence of an intermediate upstream regulator of *dsx* and *fru* splicing, is still an open question.

Within Diptera, the insect order where sex determination has been studied in the largest number of species, the *tra* ortholog has been identified only in species belonging to the Brachycera suborder [15–21]. For the basal suborder Nematocera, which includes very important hematophagous vector species such as mosquitoes, sand flies and black flies, the *tra* ortholog or its functional analog has not yet been found in any species and limited knowledge is available in general about sex determination, mainly restricted to mosquito species [22–29] (Fig. 1).

Within Nematocera, phlebotomine sand flies are second only to mosquitoes in importance as a vector of pathogens that cause diseases to humans and animals worldwide, including leishmaniases, sand fly fever, meningitis, vesicular stomatitis and Chandipura virus encephalitis [30]. Among the over 800 species of sand fly described to date, 98 are proven or suspected vectors of human leishmaniases; these include 42 *Phlebotomus* species in the Old World and 56 *Lutzomyia* species (*sensu*) in the New World [31].

Leishmaniasis are diseases of great public health concern, being endemic in over 98 countries, with more than 350 million people at risk and 2,357,000 disability-adjusted life years lost [32]. It is estimated that about 1.3 million new cases of leishmaniasis (0.2-0.4 million visceral and 0.7-1.2 million cutaneous leishmaniasis) occur every year, with 20,000-40,000 deaths caused by the visceral form. With expanding endemicity, leishmaniasis is becoming a worldwide re-emerging public health problem [33].

Despite their importance as disease vectors, most aspects of sand fly biology remain unknown, including sex determination and sexual differentiation. To fill this gap and contribute to a better understanding of the evolution of sex determination mechanisms in insects, in the present study we applied a comparative genomic/transcriptomic approach to identify and characterize sex determining genes in sand fly species. For the first time we present in a unique study the analysis of the key components of the sex determining cascade, also identifying, the first *transformer* homolog in a Nematocera species.

## Results and Discussion

### Identification of *PpeSxl, Ppetra, Ppetra-2, Ppedsx, and Ppefru* sex determining genes in the sand fly *Phlebotomus perniciosus*

In the Old World, the sand fly *Phlebotomus perniciosus* (Diptera, Nematocera) is the main vector of *Leishmania infantum* (Kinetoplastida: Trypanosomatidae), the parasitic protozoan that causes visceral and cutaneous leishmaniasis in humans and canine reservoir host, as well as of various known and emerging arboviruses considered relevant from an European public health perspective (Toscana Virus, Naples Virus, Sicilian Virus) [34]. Proteins encoded by insect sex determining genes are characterized by domains very well conserved across insect orders and distinctive of each gene family: the DNA-binding DM (<u>D</u>oublesex <u>M</u>ab3) domain for the DSX proteins [35], the protein-protein BTB (<u>B</u>road-Complex, <u>T</u>ramtrack and <u>B</u>ric a brac) binding domain for the FRU proteins [36] and the RNA-binding RRM (<u>R</u>NA <u>R</u>ecognition <u>M</u>otif) domain for SXL and TRA-2 proteins [37]. Conversely, the female-specific serine-arginine rich TRA protein exhibits a general low conservation of its primary sequence and the absence of functional characterized domains. In most of the insect species analyzed to date, the only conserved parts of the TRA protein are the TRACAM (*<u>C</u>eratitis*-*<u>A</u>pis*-*<u>M</u>usca*) domain, putatively involved in the autoregulation of the *tra* gene, the DIPTERA domain, and the HYMENOPTERA domain, the last two with unknown function [13,18,38].

We performed a TBLASTN search against the available *P. perniciosus* adult transcriptome database (http://pernibase.evosexdevo.eu) [39] to identify transcripts encoding for sex determining proteins, using other insects sex determining protein sequences as query terms (Additional file 1: Table S1). In *P. perniciosus* we identified the complete open reading frames (ORF) of the transcripts encoding for the putative SXL, TRA-2 and male- and female-specific isoforms of DSX (Additional file 2: Figures S1-S5). We named the corresponding genes as *PpeSxl, Ppetra-2*, and *Ppedsx*. In addition, we identified partial ORFs encoding for FRU, and we named the gene as *Ppefru*. The incomplete transcripts encoding for FRU proteins lack their 3’ ends and therefore complete ORFs were obtained by 3’ RACE (Additional file 2: Figures S6-S7), as described in supplementary methods. Using the TBLASTN approach no *tra* ortholog was found in the *P. perniciosus* transcriptome. This result was expected due to the low level of nucleotide and protein sequence conservation of the *tra* gene, also in closely related insect species and considering that the cloning of *tra* in *Ceratitis* was performed by synteny rather than by direct homology [15,16,40,41].

We validated the transcription and the splicing pattern of *PpeSxl, Ppetra-2, Ppedsx* and *Ppefru* by RT-PCR on mRNAs extracted from adult *P. perniciosus* males and females, using the *Ppesod* gene as endogenous positive control and to exclude genomic DNA contamination of the cDNAs (Fig. 2A). The RT-PCR primer pairs for the *PpeSxl* transcript amplified in both sexes multiple non-sex-specific transcripts probably produced by alternative splicing (Additional file 2: Figure S8), as observed in other insect species [42,43]. Functional analyses of *Sxl* in several dipteran species [44,45] show that *Sxl* is a master switch gene of sex determination only in Drosophilidae [9,46]. Therefore, we supposed that *Sxl* is probably not essential for the sex determination in *P. perniciosus* and decided to exclude it from further analyses.

**Fig. 2.**
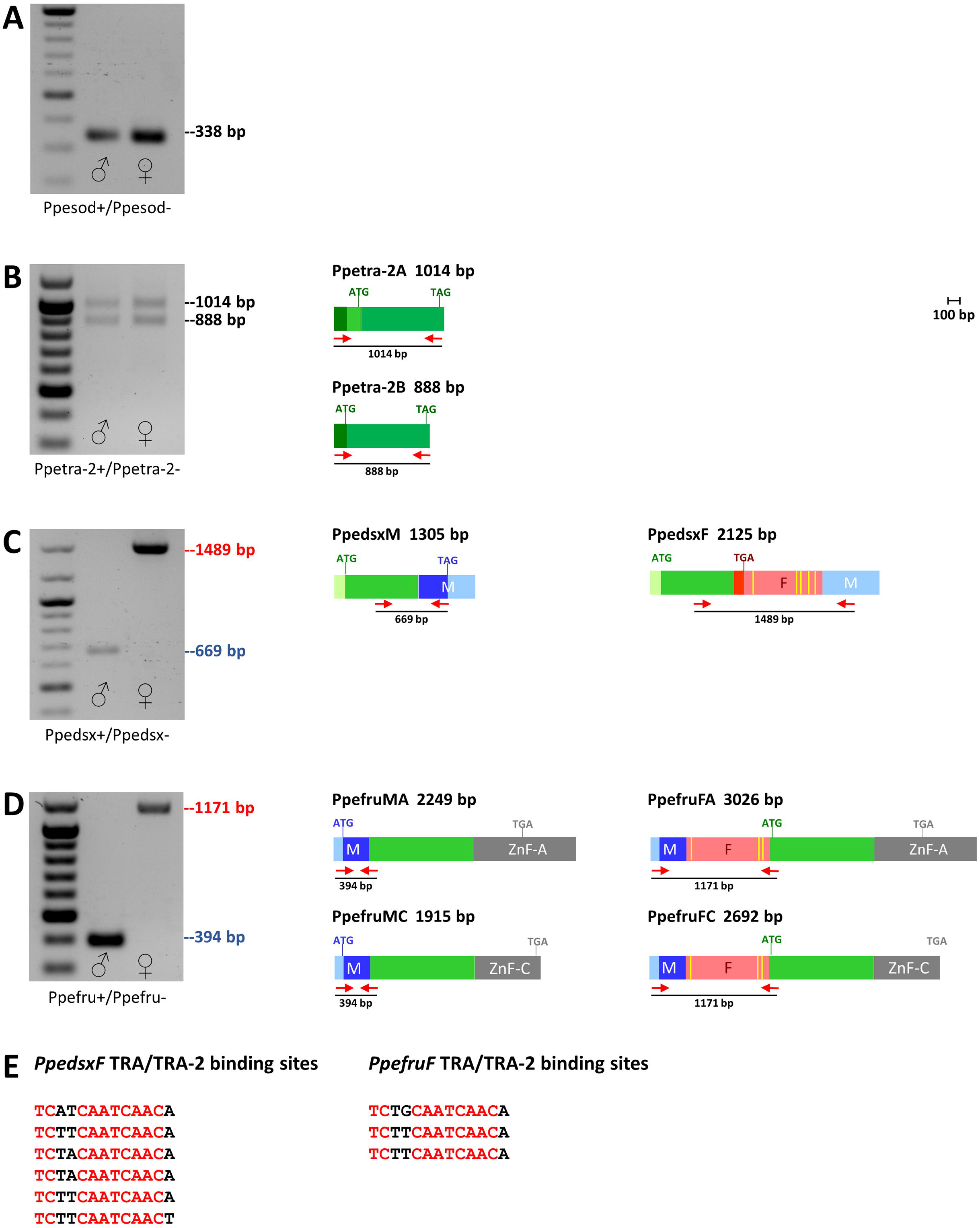
Sex determining genes expression at adult stage in *P. perniciosus*. A) Positive RT-PCR control with Ppesod+/Ppesod-primer pairs. These PCR primers span a 112-bp long intron of *Ppesod* gene and permit to exclude genomic DNA contamination of cDNAs. B) *Ppetra-2* RT-PCR amplification. C) *Ppedsx* RT-PCR amplification. D) *Ppefru* RT-PCR amplification. Light green boxes represent untranslated regions. Dark green boxes represent non-sex-specific coding regions. Azure boxes represent male-specific untranslated regions. Pink boxes represent female-specific untranslated regions. Blue and red boxes represent male-specific and female-specific coding regions, respectively. The position of primers utilized for each gene are indicated by short red arrows. Yellow vertical bars indicate the position of the putative TRA/TRA-2 binding sites. E) Putative TRA/TRA-2 binding sites identified in *Ppedsx* and *Ppefru* female-specific transcripts.

The RT-PCR analysis of the *Ppetra-2* transcript showed a non-sex-specific expression at adult stage and revealed the existence of a second isoform (*Ppetra-2B*) expressed in both sexes (Fig. 2B). Cloning and sequencing of *Ppetra-2B* showed that it encodes for a putative TRA-2 protein with slight amino acid (aa) differences in the N-terminus respect to PpeTRA-2A. A similar *tra-2* non-sex-specific splicing pattern was reported in the whitefly *Bemisia tabaci*, where the two encoded TRA-2 isoforms differ at their N-terminus for a wider region of 123 aa [47].

The RT-PCR analysis of the *Ppedsx* and *Ppefru* transcripts revealed that both genes are regulated by sex-specific alternative splicing as in other insect species (Fig. 2C-2D). Notably, in both *Ppedsx* and *Ppefru* female-specific transcripts we identified a cluster of putative TRA/TRA-2 binding sites (Fig. 2E).

In Diptera, the presence of a conserved TRA/TRA-2 binding site cluster in *dsx* and *fru* genes is always associated to the presence of the TRA active protein [15]. Encouraged by finding conserved TRA/TRA-2 binding sites in *Ppedsx* and *Ppefru* and by the presence of a PpeTRA-2 with a highly conserved RRM domain (Additional file 2: Figure S2), we pursued a strategy to identify the ortholog of *tra* in *P. perniciosus.* This approach was based on the hypothesis that also in sand flies the *tra* gene could regulate its own sex-specific alternative splicing binding a cluster of TRA/TRA-2 binding sites. Therefore, we analyzed the *P. perniciosus* adult transcriptome with the DREG tool of the Emboss Suite (http://emboss.sourceforge.net/) to detect transcripts containing putative TRA/TRA-2 binding sites. We identified an assembled transcript (c23543.g1.i2, 3858 bp-long) containing the highest number of TRA/TRA-2 binding sites, with six elements clustered in a 324bp-long sequence (Fig. 3A) and located between two putative exons encoding for a serine-arginine rich sequence. Using RT-PCR primer pairs spanning the region containing the TRA/TRA-2 binding sites we were able to amplify two male-specific (M1 and M2) and three female-specific (F1, F2 and F3) cDNA fragments (Fig. 3B), demonstrating that the c23543.g1.i2 transcript undergoes sex-specific alternative splicing regulation, as expected for a *tra* ortholog. The five full-length cDNAs were cloned and sequenced after 5’ and 3’ RACE experiments, performed as described in Methods. The virtual translation of the five cDNAs revealed that M1, M2, F2 and F3 encode for very short polypeptides due to premature stop codons. Only the female-specific F1 cDNA has a full ORF and encodes for a SR rich sequence (282 aa) containing a short region similar to the TRA Diptera domain (Additional file 2: Figure S9). We named this putative protein as PpeTRA and the corresponding gene as *Ppetra*. PpeTRA is missing the putative autoregulation TRACAM domain, which is present in all the known autoregulative TRA proteins of insects. To date, PpeTRA represents the shortest insect TRA protein, excluding the non-autoregulating TRA of *D. melanogaster* (197 aa) (Additional file 2: Figure S9).

**Fig. 3.**
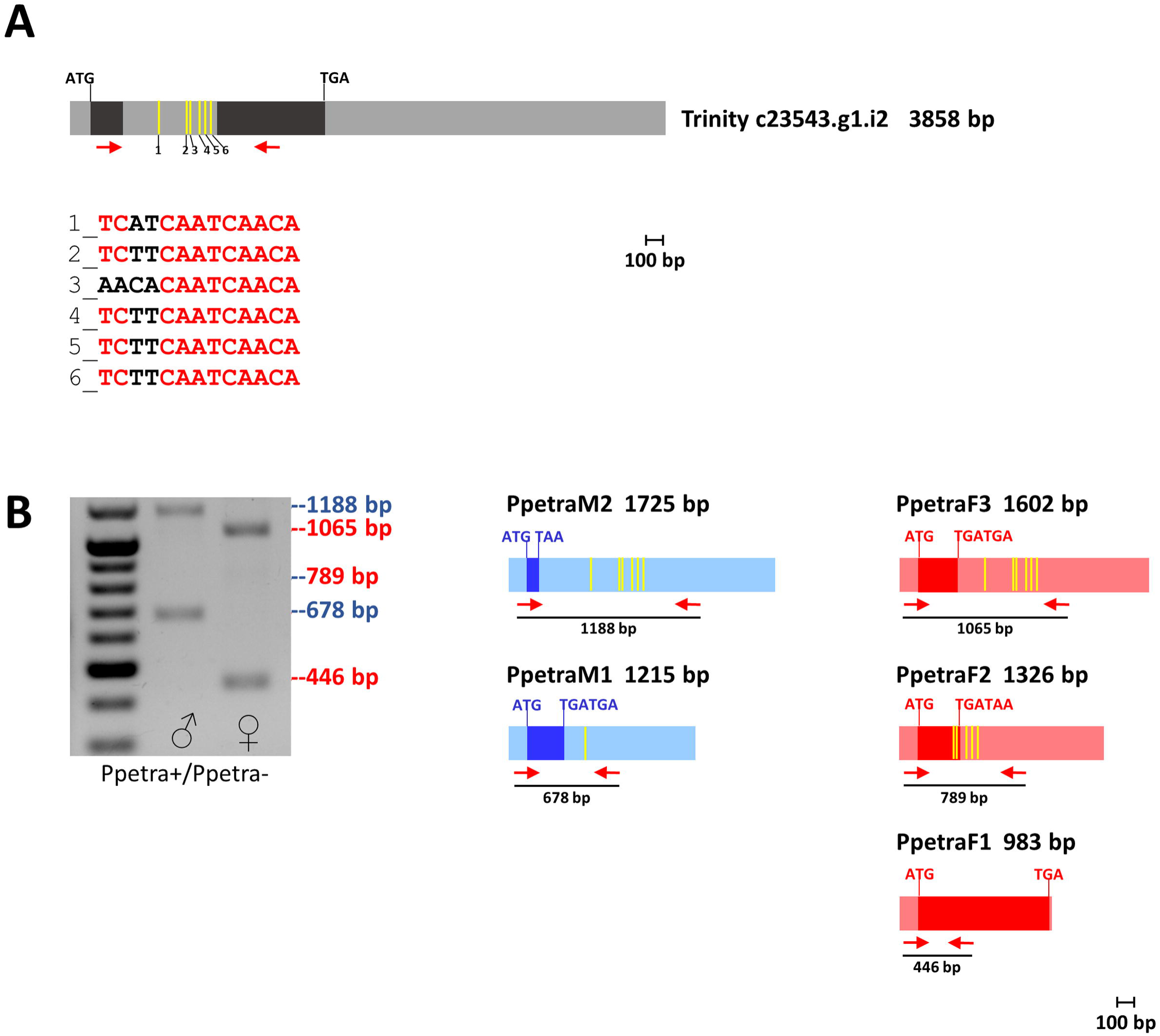
*Ppetra* transcripts and expression at adult stage. A) *In-silico* identified *Ppetra* transcript, containing six putative TRA/TRA-2 binding sites, indicated by yellow vertical bars. B) *Ppetra* RT-PCR amplification. Azure boxes represent male-specific untranslated regions. Pink boxes represent female-specific untranslated regions. Blue and red boxes represent male-specific and female-specific coding regions, respectively. The positions of primers utilized are indicated by short red arrows.

### Developmental expression analysis of sex determining genes in *P. perniciosus*

We performed an RT-PCR analysis on total RNA extracted from samples of mixed sexes from different developmental stages (embryos, larvae of 1^st^, 2^nd^ and 4^th^ instar and pupae) to analyze the developmental expression pattern of the sex determining genes newly identified in *P. perniciosus* (Fig. 4). We used the *Ppesod* gene, constitutively expressed in *P. perniciosus* (Petrella et al., 2015), as endogenous positive control, and the same primer pairs of the RT-PCR analyses performed on adult samples, spanning the alternatively spliced regions of *tra, tra-2, dsx* and *fru* genes, as reported in figure 2 and 3. We found that *Ppetra* is expressed since embryonic stage as observed for other dipteran species [16,18,48], producing sex-specific transcripts. We amplified, in all developmental stages, fragments of 446bp, 678 bp and 1065 bp corresponding to *Ppetra* F1, M1 and F3 transcripts, respectively (Fig. 4). *Ppetra-2* is expressed from the first instar larval stage until adulthood, differently from other dipteran species, such as *Ceratitis capitata* and *Musca domestica*, where it is expressed also at embryonic stage [49,50]. Both the *Ppetra-2A* and *B* transcripts were detected in all stages but embryos (Fig. 4). *Ppedsx* and *Ppefru* are expressed from first-instar larval stage and second-instar larval stages, respectively, until adulthood, both producing sex-specific transcripts by alternative-splicing during development (Fig. 4). *Ppedsx* developmental expression pattern seems to be different respect to other dipteran species, including *Drosophila, C. capitata* and the tiger mosquito *Aedes aegypti*, where *dsx* is expressed also at the embryonic stage [23,51,52]. Conversely, the *Ppefru* developmental expression pattern is conserved respect to *fru*-P1 promoter expression pattern observed in *Drosophila* and in *A. aegypti* (Salvemini et al., 2010; 2013) with expression starting at late larval stage until adulthood.

**Fig. 4.**
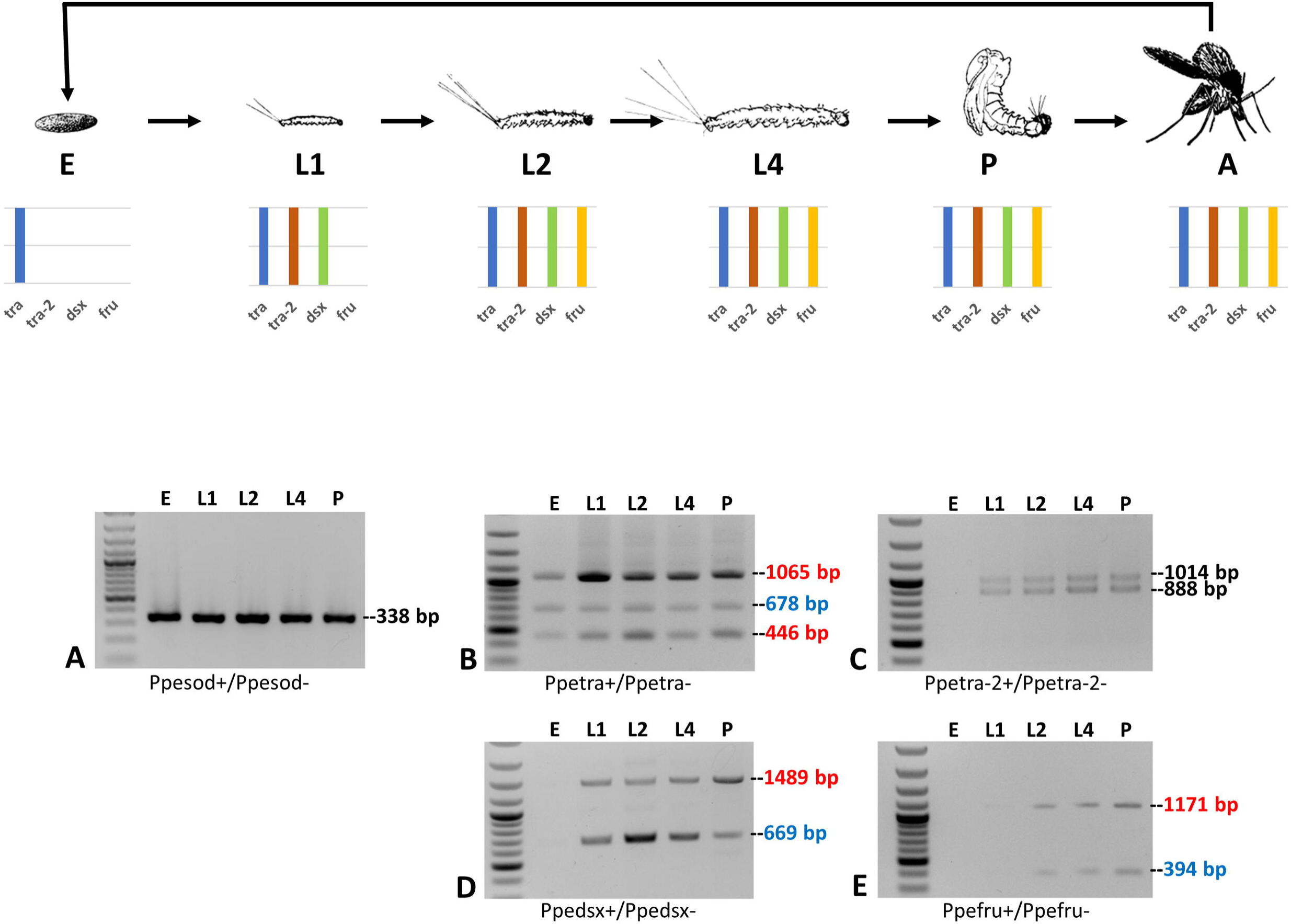
Sand fly life cycle and developmental expression analyses of sex determining genes in *P. perniciosus*. RT-PCR amplifications of *Ppesod* (A), *Ppetra* (B), *Ppetra-2* (C), *Ppedsx* (D) and *Ppefru* (E) were performed on the following samples: E = 0-36h embryos; L1 = first instar larvae; L2 = second instar larvae; L4 = fourth instar larvae; P = pupae; all samples are composed of mixed sexes. The *P. perniciosus sod* gene, utilized as positive control, is constitutively expressed throughout development. The coloured bars indicate the presence/absence of expression at each developmental stage of *Ppetra* (blue), *Ppetra-2* (green), *Ppedsx* (green) and *Ppefru* (yellow) transcripts.

### Evolution of *tra* genomic organization and of alternative splicing regulation in Phlebotominae

The *Ppetra* gene is the first *tra* ortholog isolated in a Nematocera species and the shortest *tra* gene (1.7Kb) isolated to date in insects. To study its genomic organization we amplified, cloned and sequenced the 1725 bp fragment corresponding to the *Ppetra* locus, using a primer pair located in the 5’ and 3’ UTR of *Ppetra* transcripts and adult genomic DNA. Aligning genomic *Ppetra* against the five *Ppetra* cDNA sequences, we reconstruct the exon-intron organization of the *Ppetra* gene and identified the alternative splicing events producing the *Ppetra* transcript isoforms (Fig. 5A). The *Ppetra* gene has four exons and three introns, all with conserved GT-AG boundaries (Additional file 3: Figure S10). In females, *Ppetra* produces three transcripts. Exon 1, 2, 3 and 4 are used to produce a mature mRNA corresponding to the F1 transcripts, with an ORF encoding for the 282 aa-long PpeTRA protein. In addition to this, distinct parts of intron 2 are retained in two other transcripts, one by an alternative 3’ acceptor splicing site (transcript F2) and the other by an intron retention mechanism (transcript F3). In both the F2 and F3 transcripts the presence of in-frame stop codons causes short truncated PpeTRA isoforms. In males, *Ppetra* produces two transcripts: the M2 transcript is an unspliced transcript because it retains all the introns, while the M1 transcript is produced through an alternative 5’ donor splicing site choice. In the two *Ppetra* males-specific transcripts, the introduction of premature stop codons leads to short truncated PpeTRA isoforms.

**Fig. 5.**
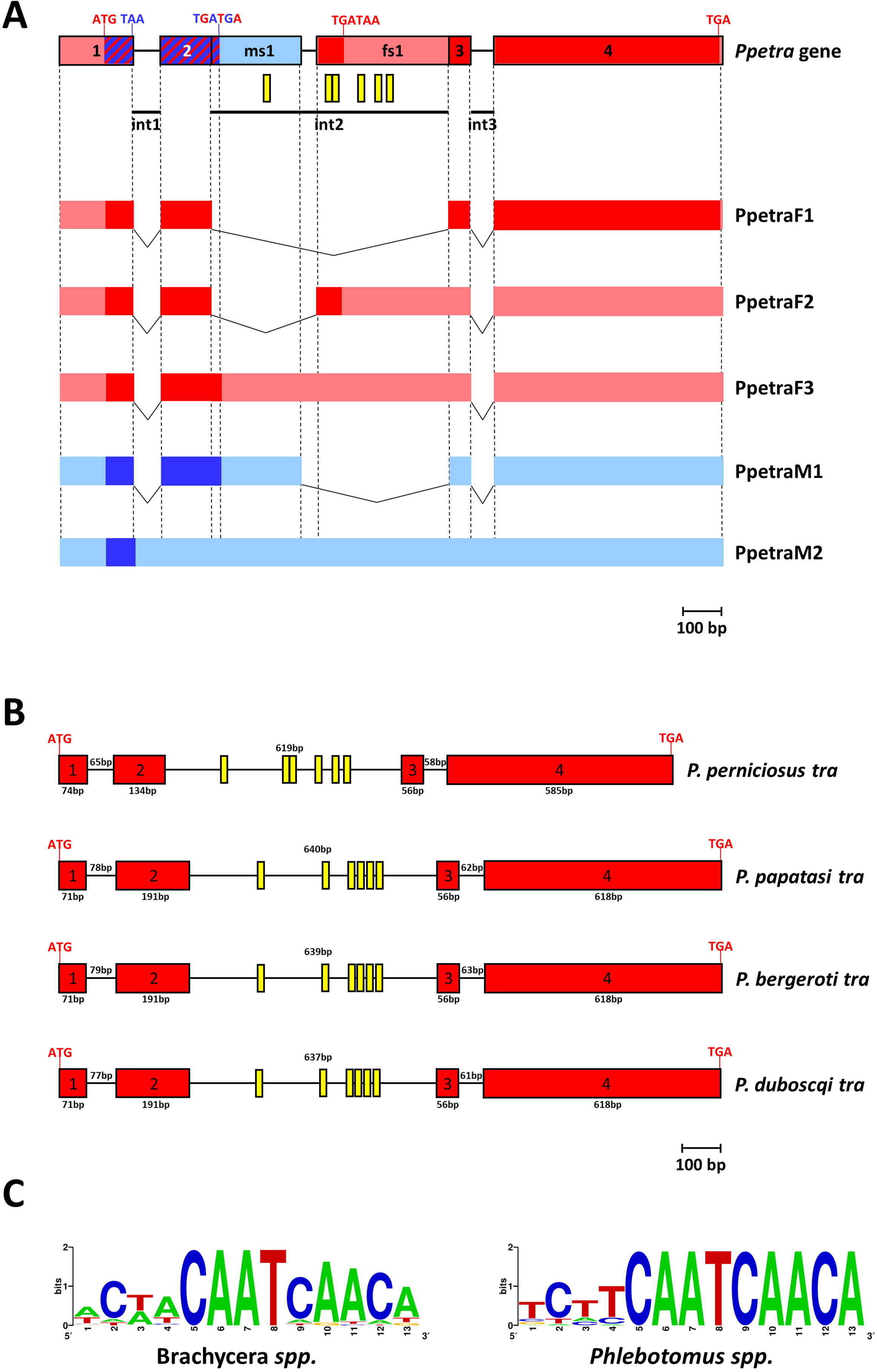
*P. perniciosus tra* genomic organization and evolution. A) *Ppetra* gene locus and sex-specific transcripts. Azure boxes represent male-specific untranslated regions. Pink boxes represent female-specific untranslated regions. Blue and red boxes represent male-specific and female-specific coding regions, respectively. Striped blue-red boxes represent coding regions utilized both in the male and female sex. Yellow vertical bars indicate the position of the putative TRA/TRA-2 binding sites. B) Comparison of *tra* gene structures in *Phlebotomus* species. C) WebLogo consensus sequence of the putative TRA/TRA-2 binding sites identified in *Phlebotomus tra* species and of TRA/TRA-2 binding sites of Brachycera *tra* genes.

To study the evolution of the genomic organization and alternative splicing regulation in sand flies, we searched the *tra* orthologs in seven other Phlebotominae species by TBLASTN using PpeTRA as query. For two species, *P. papatasi* and *L. longipalpis*, genome and transcriptome assemblies were available (PpapI1, PpapI1.4, LlonJ1 and LlonJ1.4; https://www.vectorbase.org/). For the other two Old World sand fly species, *P. bergeroti* and *P. duboscqi*, we assembled a draft genome using available sequencing data and the MINIA genome assembler [53]. In addition, we produced *de novo* transcriptome assemblies by using all the available sequencing data (up to March 2018) for *L. longipalpis* and for two New World species, *L.* (*Nyssomyia*) *umbratilis* and *L.* (*Nyssomyia*) *neivai* using the Trinity *de novo* assembler [54,55] (see Additional file 15: supplementary methods).

By querying PpeTRA against the genomes and transcriptomes of the Phlebotominae species, we identified the *tra* ortholog in *P. papatasi* (*Ppatra*), *P. bergeroti* (*Pbetra*) and *P. duboscqi* (*Pdutra*) but this approach could not identify any ortholog in the genomes/transcriptomes of the three New World sand fly species. Furthermore, neither the TRA/TRA-2 binding sites *in silico* approach, that led to the identification of *Ppetra*, nor a molecular approach in *L. longipalpis* by touch down RT-PCR with degenerated primers designed on the alignment of *Ppetra* and *Ppatra* sequences (data not shown) could identify *tra* in New World sand flies. Similarly, Geuverink and Beukeboom [38] *in silico* identified a putative *tra* gene in the Old World sand fly species *P. papatasi* and reported its apparent absence in the New World sand fly *L. longipalpis.* As for *Ppetra*, the putative *P. papatasi* TRA missed the most consistent recognition motif of a TRA protein, i.e. the putative autoregulation TRACAM domain, leading the authors to be cautious about the true nature of their identified sand fly *tra* ortholog [38].

We reconstructed gene models for the *P. papatasi, P. bergeroti* and *P. duboscqi tra* orthologs (Additional file 3: Figures S11-S13), which encode for a 311 aa-long SR-protein with 61% identity respect to the PpeTRA and missing, as in *P. perniciosus*, a conserved TRACAM domain (Additional file 4: Figure S14). The four *Phlebotomus tra* genes revealed a conserved genomic organization with four exons and three introns, with small differences in exons/introns lengths (Fig. 5B). In the intron 2 of *P. papatasi, P. bergeroti* and *P. duboscqi*, we identified, as in *P. perniciosus*, six conserved TRA/TRA-2 binding sites (Fig. 5C). To study the alternative splicing regulation of the *tra* gene in *Phlebotomus* species, we compared the intronic sequences of the four species (Additional file 5: Figure S15). As in *P. perniciosus*, all *tra* introns exhibit conserved GT-AG terminal dinucleotides. Intron 2, which is regulated by sex-specific alternative splicing in *P. perniciosus*, has a putative conserved alternative splicing sites (SS) also in *P. papatasi, P. bergeroti* and *P. duboscqi*. In the four species, the 5’ donor SS of intron 2 seems to be weak and suboptimal, while the 3’ acceptor SS is a canonical strong splicing site. Finally, all the four species have a strong canonical male-specific alternative 5’ donor SS at about 230 bp downstream of exon 2 (Additional file 5: Figure S15). These findings led us to suppose that in *P. perniciosus*, as well as in the other three *Phlebotomus* species, the male-specific splicing of the *tra* pre-mRNA represents the default splicing mode. In contrast, in females, the repression of the male-specific 5’ donor SS of intron 2 is most probably due to the binding of TRA and TRA-2 proteins on the TRA/TRA-2 binding site cluster, leading to the usage of the upstream 5’ donor SS to form to the female-specific *tra* transcript, thus producing a functional TRA only in females. This hypothesis on the conserved splicing regulation was confirmed in *P. papatasi* by RT-PCR on adult RNA from males and females (Additional file 6: Figure S16).

Figure S17 shows a comparison of the *tra* genomic locus among insect species (Additional file 6: Figure S17). Despite differences in exon number and intron length, the sex-specific splicing regulation of the *tra* gene exhibits a striking conservation. In all the considered species, including *P. perniciosus*, an alternative 5’ donor SS choice leads to a full TRA protein only in the female sex. To study the protein organization, we compared PpeTRA with other arthropod TRA proteins (Fig. 6A). PpeTRA exhibits similar domain organization respect to insect TRAs, with a DIPTERA domain located within the RS domain, as observed also in TRA of *Lucilia cuprina* (LcTRA), *Cochliomyia hominivorax* (ChTRA) and *Glossina morsitans* (GmTRA). At the same time, PpeTRA misses the TRACAM domain; the *Ppetra* regions corresponding to the last 31 nucleotides of exon 2 and to the first 45 nucleotides of exon 3 (upstream and downstream of the *Ppetra* sex-specifically regulated intron, respectively) encode for a PpeTRA protein portion that exhibits only 8 out of 25 conserved amino acids respect to the insect TRACAM domain (Fig. 6B).

**Fig. 6.**
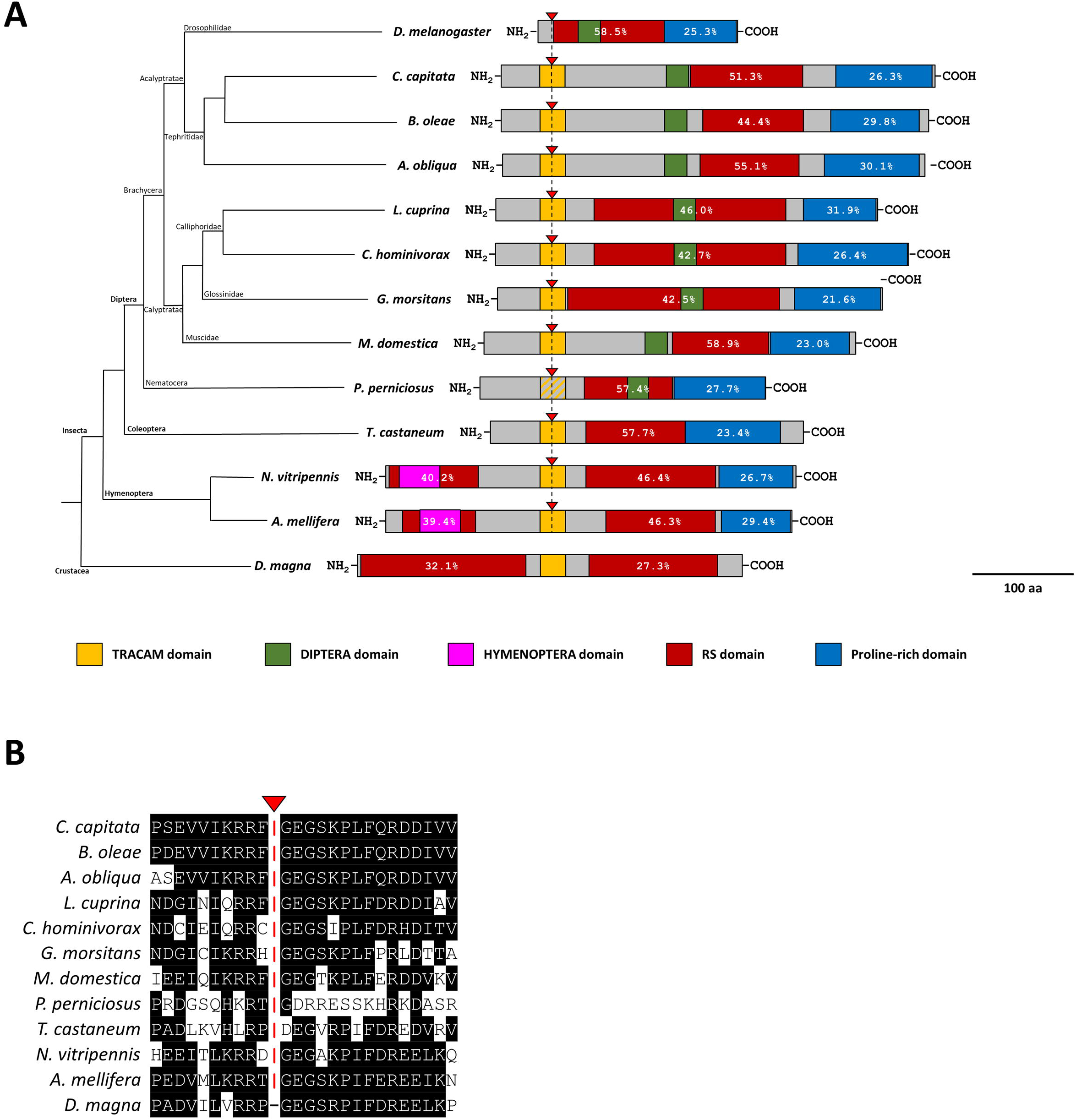
Phylogenetic relationship and protein sequence comparison of TRA/FEM proteins. A) TRA/FEM protein schemes were aligned using the conserved sex-specific splicing site located within the TRACAM domain encoding region as reference point (indicated by the red triangle). This sex-specific splicing site is conserved in all the autoregulating *tra* genes. In sand flies TRA, the TRACAM domain is absent. Striped yellow-grey box represents the position of the homologous sex-specific splicing site in *P. perniciosus* TRA. *D. melanogaster* TRA protein was aligned using the position of the non-conserved sex-specific splicing site. The crustacean *Daphnia magna* TRA was aligned using the position of the conserved TRACAM domain. Percentages within red and blue boxes indicates the percentage of R and S residues and of P residues in the RS and Proline-rich domains, respectively. To define the boundaries of the RS domain, we considered the position of the first RS or SR couple of residues till the final RS or SR couple of residues and we considered the selected region an RS domain only if its percentage of R and S is > of 25%. B) Multiple alignment of insect TRACAM domains and the *P. perniciosus* TRA homologous region. Amino acids conserved in at least two species are highlighted in black. The conserved sex-specific splicing site is indicated by red triangle.

In conclusion, the conserved *tra* structure, the conserved sex-specific alternative splicing regulation and the presence of a conserved TRA/TRA-2 binding site cluster in the sex-specifically regulated *tra* intron strongly support the hypothesis of the autoregulation of the *tra* gene in sand flies, as observed for other dipteran and non-dipteran species. At the same time, the absence in sand fly TRAs of a putative TRACAM domain (supposedly involved in the *tra* autoregulation) [18], led us to hypothesize a still unknown molecular mechanism of the TRA autoregulation specific to sand flies.

### Evolution of *tra-2* genomic organization in Phlebotominae

*tra-2* is a single-copy gene that has been characterized in *D. melanogaster* [56,57] and in several other dipteran species such as *D. virilis* [58], the house fly *M. domestica* [50], the tephritids *C. capitata* [49,59] and twelve *Anastrepha* species [60], the calliphorid *Lucilia cuprina* [19], and the Nematocera sciarid species *Sciara ocellaris* and *Bradysia coprophila* [61]. In these species, *tra-2* is transcribed during development in both sexes, producing an RNA-binding protein with two RS domains flanking an RRM domain. TRA-2 RRM is followed by a 19 aa-long linker region, which is a distinctive and unique feature of the TRA-2 proteins [62]. Within Brachycera suborder, TRA-2 is required for the sex-specific splicing regulation of the *dsx* and *fru* genes and, outside Drosophilidae, it is also involved in the autoregulation of female-specific alternative splicing of the *tra* gene [18,49,50,60,63].

Using the available genomic resources of *P. papatasi*, the assembled draft genomes of *P. bergeroti* and *P. duboscqi* and the identified putative TRA-2 proteins of *P. perniciosus*, we reconstructed the partial putative exon-intron structure of *tra-2* of the Old World sand flies consisting of 4 exons and 3 introns (Additional file 7: Figures S18-20). In addition, we identified in *P. papatasi, P. bergeroti* and *P. duboscqi* a putative alternative 5’ donor splicing site located downstream of the 5’ donor splicing site of the exon 1 of *tra-2*, which is conserved in *P. perniciosus* where it leads to the production of the PpeTRA-2B isoform. As these species belong to different subgenera (*Phlebotomus* and *Larroussius*) this suggests that a similar non-sex-specific alternative splicing event could be conserved also in other Old World sand flies (Additional file 7: Figures S18-20). More in general, among dipteran, *tra-2* shows an overall conservation of exons encoding for functional domains and both RRM and RS1 domains are coded by several exons. In Nematocera Old World sand flies, the RS1 domain is encoded by a unique exon, while the RRM domain and the linker region are organized in two exons (Additional file 8: Fig. S21).

As observed for the *tra* ortholog, *tra-2* seems to be absent in transcriptome and genome assembly of the New World sand fly *L. longipalpis*. However, we found well conserved TRA-2 encoding transcripts missing the N-terminus coding region in the *L. umbratilis* and *L. neivai*. This finding suggests that the *tra-2* ortholog could be present also in *Lutzomyia* but not correctly assembled in the *L. longipalpis* released transcriptome/genome assemblies. In figure S22 the multiple alignment of sand fly putative TRA-2 protein is reported. A very well conserved RRM+linker region and RS1 region are present in all the species analyzed. A RS2 region was detected only in *P. perniciosus, L. umbratilis* and *L. neivai*. The high percentage of conserved residues of the RS2 region (22/51) suggests its conservation also in other *Phlebotomus* species (Additional file 8: Fig. S22).

In summary, with our work we identified for the first time the *tra-2* gene in sand flies. Previously, the *tra-2* ortholog of Nematocera was characterized only in the sciarid species *S. ocellaris* and *B. coprophila* and in the mosquito *An. gambiae* and *Ae. aegypti*, where two and four orthologs were found, respectively. In *S. ocellaris* and *B. coprophila* TRA-2 is highly conserved and shows conserved sex-determination function when expressed in *Drosophila* [61]. Conversely, putative TRA-2 identified in mosquitoes seem to be divergent respect to other dipteran TRA-2 and possibly not involved in the control of sex-specific splicing of *dsx* and *fru* targets [23,64]. Recent functional tests by transgene-mediated RNAi against *Ae. aegypti tra-2* orthologs have shown no female-to-male sex reversion, as obtained in *tra-2* RNAi functional studies in Brachycera species, but a novel female-specific zygotic lethality. This finding supports the hypothesis that *tra-2* does not play a conserved role in *Ae. aegypti* sex determination while it controls a novel female-specific vital functions which need to be clarified [65]. Here we show that, as for *tra-2* of sciarid species, in sand flies *tra-2* encodes for a protein conserved in its structure and domains, suggesting a conserved role in the sex determination through sex-specific alternative splicing regulation of both *dsx* and *fru* downstream target genes. In addition, we propose that *tra-2* could be involved in the autoregulation of the *tra* gene also in Old World sand flies. The absence of a *tra* ortholog in New World sand flies poses a very interesting problem about the function of *tra-2* in these species and about the evolution of the alternative splicing regulation of *dsx* and *fru* genes.

### Evolution of *dsx* and *fru* genomic organization and alternative splicing regulation in Phlebotominae

To study the evolution of the genomic organization and of the alternative splicing regulation of *dsx* and *fru* genes in the sand flies, we aligned DSXs and FRUs of *P. perniciosus, P. papatasi* and *L. longipalpis* against the genome sequences of *P. papatasi* and *L. longipalpis* using TBLASTN (Additional file 9: Figures S23-S26, Additional file 15: Supplementary Methods). By manually refining the exon-intron junctions, we obtained the structure of the genes. Compared with the orthologs in *D. melanogaster, An. gambiae* and *Ae. aegypti,* we observed an overall conservation of the exon/intron organization and of the alternative splicing regulation in sand flies (Additional file 9: Figures S27-S28).

In particular, as observed in other dipteran species [23,25,51,52,66,67], in sand flies *dsx* is organized in 4 exons spread over a large genomic region (146 Kb in *P. papatasi* and at least 191 Kb in *L. longipalpis*). Exon one, which contains the ATG signal, encodes for the DSX OD1 domain and is followed by the second exon encoding for the non-sex-specific part of the DSX OD2 domain. Exon three is female-specific and encodes for the female-specific DSX C-terminus. Exon four is present in transcripts of both sexes as 3’untranslated region in females and encoding for male-specific DSX C-terminus in males (Fig. S27). Interestingly, the nucleotide sequence of the region surrounding the 3’ acceptor female-specific splicing site of the *dsx* gene is strictly conserved among *Phlebotomus* species (Additional file 9: Figures S29). A similar observation was recently reported by Kyrou and colleagues [68] for *Anopheles* mosquito. This region was utilized to develop a gene drive-based population suppression strategy resulted very effective in small scale caged experiments [68]. This finding suggests that also the *dsx* gene of *Phlebotomus* species could be an ideal target to develop future similar strategies for sand fly control in field.

The *fru* gene in sand flies is organized in eight exons distributed over a very large genomic region (at least 125 Kb in *P. papatasi* and 213 Kb in *L. longipalpis*). Exons one and two (named S1 and S2, respectively) are common and female-specific respectively, with exon S1 encoding the male-specific N-terminus of FRU and exon S2 utilized only in females as 5’ untranslated region. Exons three and four (named C1 and C2) encode for the BTB domain, while exons five to seven (named C3, C4 and C5) encode the poorly conserved Connector region. The terminal exon eight encodes for a zinc-finger domain of type C (Additional file 9: Figure S28). Using the *P. perniciosus fru* ZnF-A and the *D. melanogaster* protein sequence of ZnF-B as queries, TBLASTN analysis of the genomic scaffold 549 of *P. papatasi* PpapI1 assembly, containing the *fru* exon eight, revealed the presence of putative exons encoding very well conserved ZnF domains. This finding suggests that also in sand flies the *fru* gene could encodes for multiple FRU isoforms by alternative splicing at the 3’end of the primary transcripts (data not shown).

Figure 7A shows a schematic representation of the sex-specifically regulated regions of both *dsx* and *fru* genes in *D. melanogaster, An. gambiae, Ae. aegypti, P. papatasi* and *L. longipalpis*. As for most of the Brachycera species, in *Drosophila dsx* and *fru* sex-specific alternative splicing is achieved through two different mechanisms. For *dsx*, a 3’ alternative acceptor splicing site choice coupled with alternative polyadenylation leads to sex-specific transcripts with different 3’ ends encoding for sex-specific DSX C-termini (Fig. 7A) [69]. For *fru*, a 5’ alternative donor splicing site choice leads to sex-specific transcripts with different 5’ ends. In males, a male-specific FRU, with a unique N-terminus is obtained through the usage of an ATG signal present in the *fru* male-specific exon (Fig. 7A) [70]. In females, a stop codon in the female-specific exon produces a transcript with a very short open reading frame, probably not translated (Fig. 7A). For both the genes, the male-specific splicing represents the default mode. In female, the presence of TRA and the consequent formation of the TRA/TRA-2 complex which binds the TRA/TRA-2 binding sites in *dsx* and *fru* female-specific exons, promotes female specific splicing [69–71].

**Fig. 7.**
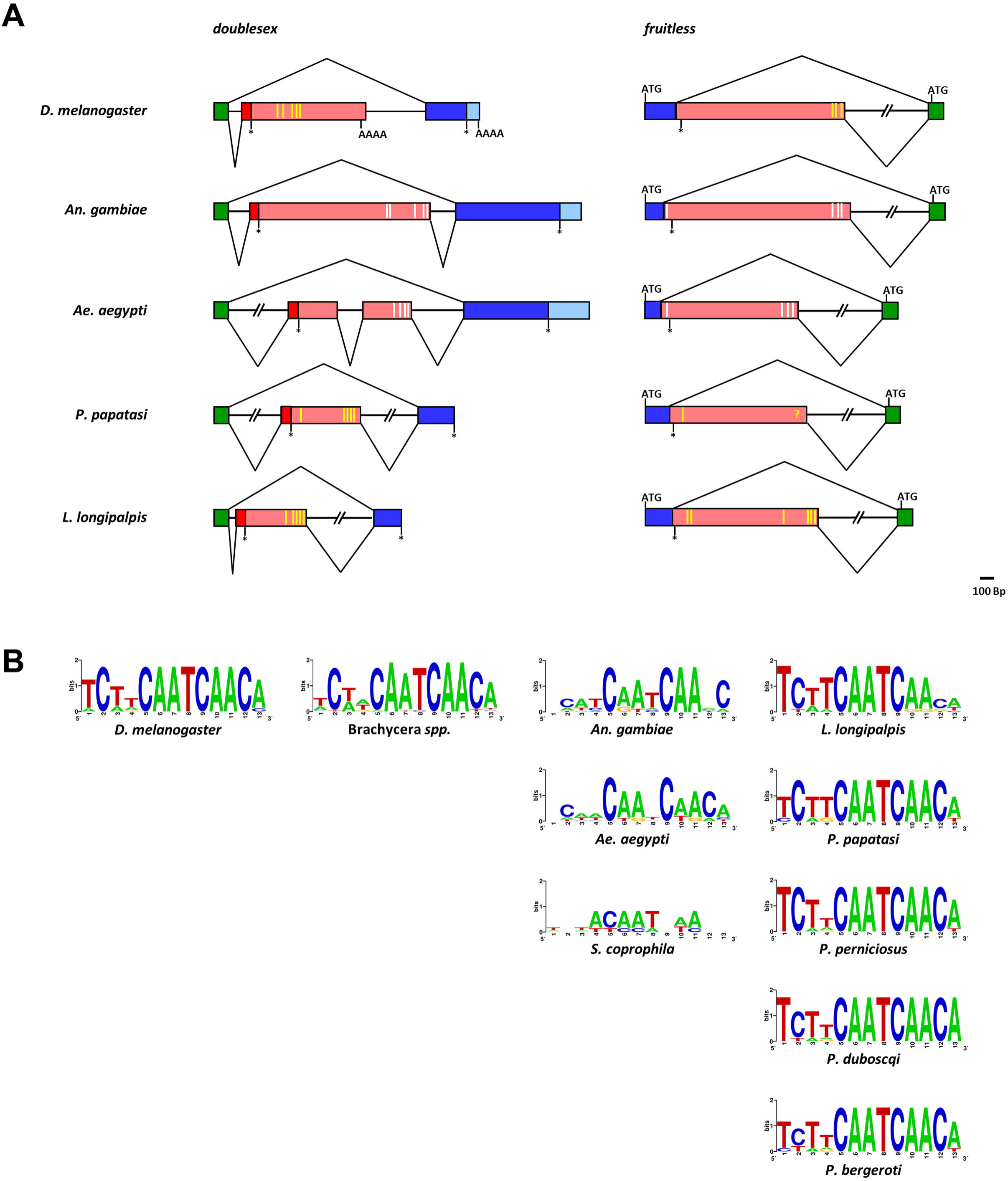
Evolution of sex-specific alternative splicing regulation of *dsx* and *fru* gene. A) Comparative schematic representation of sex-specifically regulated regions of dsx and fru genes in *D. melanogaster*, mosquito and sand fly species. Green boxes represent non-sex-specific coding regions. Azure boxes represent male-specific untranslated regions. Pink boxes represent female-specific untranslated regions. Blue and red boxes represent male-specific and female-specific coding regions, respectively. Yellow vertical bars indicate the position of the putative TRA/TRA-2 binding sites. White vertical bars indicate the degenerated mosquito putative TRA/TRA-2 binding sites. B) WebLogo consensus sequence of the putative TRA/TRA-2 binding sites of Brachycera and Nematocera species. Within sand flies, *L. longipalpis* exhibits the less conserved TRA/TRA-2 binding sites, as expected for a species with upstream regulator/s of the alternative splicing of *dsx* and *fru* genes, different from *tra*.

In Nematocera, *dsx* and *fru* orthologs have been characterized in very few species including the mosquito *An. gambiae* and *Ae. aegypti* [22–25,72–74]. While sex-specific splicing regulation of the *fru* orthologs in both mosquito species is very well conserved respect to *Drosophila* (Fig. 7A) [22,24], for *dsx* a different mechanism was described in each species. In *An. gambiae*, male-specific DSX is obtained by skipping the female-specific *dsx* exon; instead the male-specific exon sequence is used in females as 3’ untranslated region due to the absence of an alternative polyadenylation signal [25]. In *Ae. aegypti, dsx* presents two female-specific exons, like in Sciaridae [73], that are escaped in males. In females, inclusion of both or only the second female-specific exon results in two isoforms. In both *Ae. aegypti* and *An. gambiae*, due to the absence of an alternative polyadenylation signal in the female-specific *dsx* exons, male-specific exons are used as 3’ untranslated region [75].

In sand flies, *fru* has a very well conserved alternative splicing regulation, identical to *D. melanogaster* and mosquitoes, based on a 3’ alternative acceptor splicing site choice mechanism. The *dsx* gene alternative splicing regulation is instead similar to *An. gambiae* regulation, with an exon-skipping of a female-specific cassette exon only in males and with the males-specific exonic sequence, present also in female-specific transcripts, used as untranslated region (Fig. 7A).

The analysis of *dsx* and *fru* female-specific exons in *P. perniciosus, P. papatasi* and *L. longipalpis* revealed the presence of clusters of the *cis*-acting regulatory element named TRA/TRA-2 binding sites. In particular, we identified nine elements in *P. perniciosus* (six located in the *PpedsxF* and three in the *PpefruF* transcripts), six elements in *P. papatasi* (five located in the *PpadsxF* and one in the *PpafruF* transcripts) and eleven elements in *L. longipalpis* (five located in the *LlodsxF* and six in the *LloefruF* transcripts) (Additional file 12: Table S2). The identified TRA/TRA-2 binding sites are organized in clusters of at least three elements except for the single element identified in the *PpafruF* female-specific exon (Additional file 9: Figure S25A).

As for *P. perniciosus* (subgenus *Lariossus*), in both *P. bergeroti* and *P. duboscqi* the *fru* S1 exon, encoding for the putative male-specific FRUM N-terminus, is followed by a putative female-specific S2 exon containing three conserved TRA/TRA-2 binding sites (Additional file 11: Figures S30-S33). Similarly, the *dsx* female-specific exon in *P. bergeroti* and *P. duboscqi* shows six clustered TRA/TRA-2 binding sites, as observed in the other sand fly species (Additional file 11: Figures S32-S33). The absence of a cluster of TRA/TRA-2 binding sites in *P. papatasi fru* could be also due to an incorrect assembly of the corresponding *fru* genomic region.

Intra-species alignment of the TRA/TRA-2 binding sites in sand flies revealed high sequence conservation. In figure 6B, the WebLogo (http://weblogo.berkeley.edu/) consensus sequences for TRA/TRA-2 binding sites of various dipteran species are reported. Differently from other Nematocera species, such as the mosquitoes *An. gambiae* and *Ae. aegypti* and the sciarid fly *S. ocellaris*, within each 13-bp long TRA/TRA-2 binding sites of sand flies we observed an invariable “core” of 8 bp (CAATCAAC) and a low variability, as observed in *Drosophila*, in the first four bases and in the terminal base of the element. In a previous work we proposed that in mosquitoes, the degeneration of the putative TRA/TRA-2 binding sites is related with the absence of the *tra* ortholog and with the low level of TRA-2 conservation, suggesting that different upstream regulators are involved in the control of *dsx* and *fru* genes in this Nematocera species [22]. Conversely, the high conservation of the TRA/TRA-2 binding sites in *Phlebotomus*, which resembles the sequence conservation level of the TRA/TRA-2 binding sites observed in *dsx* and *fru* genes of Brachycera, indicates that these elements, located in untranslated regions of both genes, are under strong selective pressure. Overall, our findings suggest that also in sand flies TRA and TRA-2 are involved in the regulation of the sex-specific alternative splicing of *dsx* and *fru* genes, as observed in Brachycera.

### Phylogeny and selection at sex determination genes in sand flies

Figure 8 shows the Neighbor-Joining trees obtained from amino acid alignments of selected domains of the TRA (Fig. 8A), TRA2 (Fig. 8B), DSX (Fig. 8C) and FRU (Fig. 8D) of *P. perniciosus* and other species (see Methods and Additional file 13: Table S3). For all proteins, phylogenies segregates sequences in general agreement with the species phylogeny. We investigated natural selection at molecular level as the ratio between the mean nonsynonymous and synonymous substitution rates (ω) of the examined coding regions. To check if the ω ratios differed significantly among the tree branches, we compared one-, two- and three-ratio models [76] for each gene. The one-ratio model assumes an equal ω for all the branches, whereas the two- and three-ratio models consider two and three different ω values, respectively. In addition, we tested the branch-site model that assumes positive selection at specific sites within specific the tree branches [77,78]. The results obtained, and the statistical significance of each comparison, are shown in supplementary Table S4 (Additional file 14: Table S4). Overall ω is always lower than 1, showing that purifying selection acts on these genes (Additional file 14: Table S4).

**Fig. 8.**
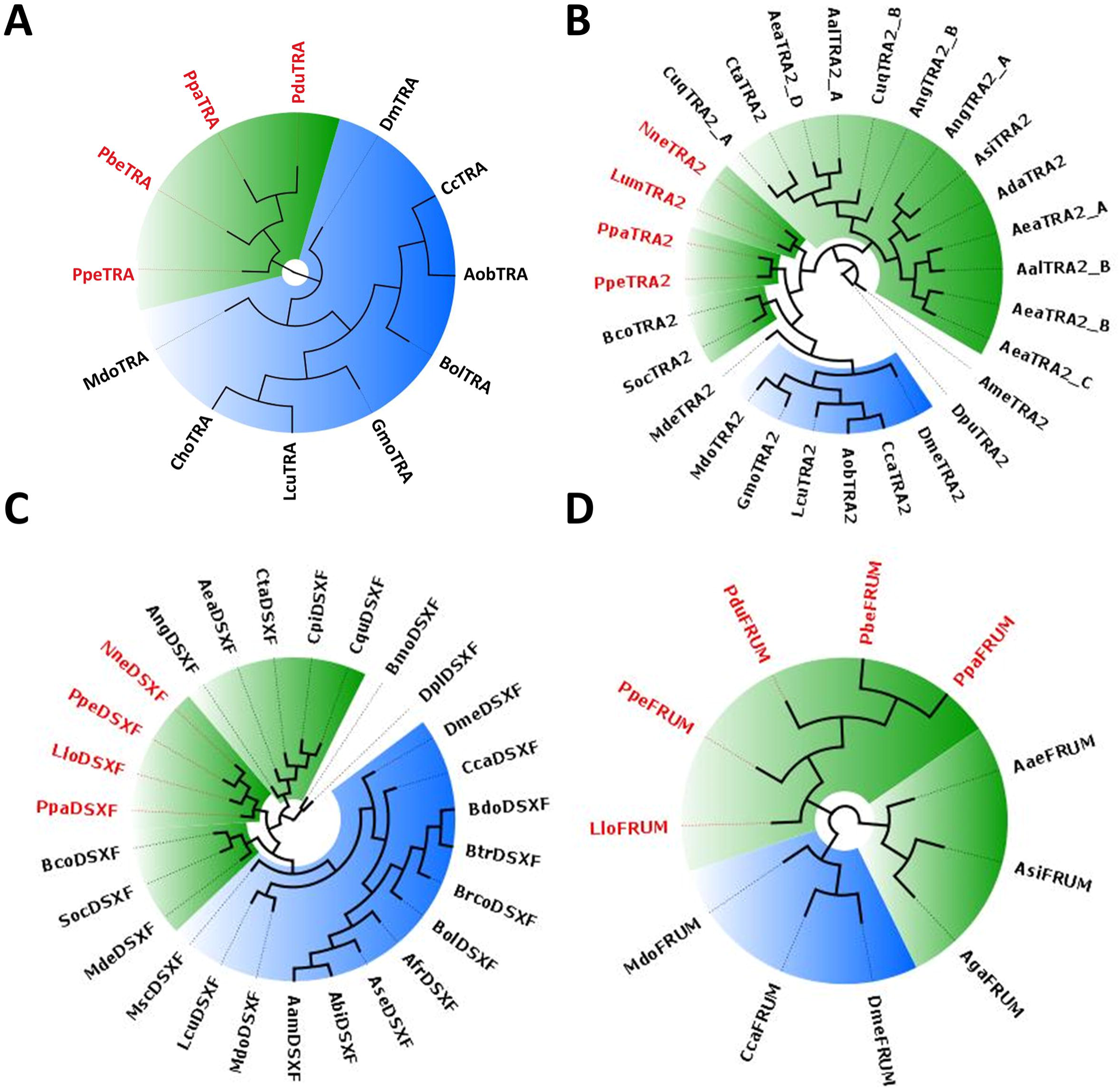
Neighbor-Joining trees obtained from the amino acid alignment of selected domains of the TRA, TRA-2, FRU and DSX proteins. For the TRA alignment (A) TRACAM and DIPTERA domains were utilized. For the TRA-2 alignment (B) we utilized the RRM domain and the linker region. For the DSX alignment (C) OD1 and OD2 domains of the DSXF isoform were utilized. For FRU alignment (D) we utilized the male-specific N-terminal region and the BTB domain. Brachycera species are highlighted in blue and Nematocera species are highlighted in green. The protein IDs of the species belonging to Phlebotominae are in red.

The evolutionary analysis of the TRACAM and DIPTERA *tra* domains shows that the one-ratio model best fits the data (ω = 0.0864) and the absence of positive selection.

Within the *tra-2* RMM and linker domains, the two-ratio model is better supported than the one- and three-ratio models, with the mosquito branch showing the lowest ω value (0.0407) when compared to the other branches (ω = 0.0738). The branch-site model identifies two positively selected sites within the branch that does not include mosquitoes; however, the comparison with its null model is not statistically supported.

Within the *dsx* OD1 and OD2 domains, the one-ratio model can be excluded in favor of the two- and three-ratio models. The two-ratio model fits the data better than the three-ratio model, showing more relaxed selective constraints of the Phlebotominae branch (ω = 0.0732) when compared to the other branches of the tree (ω = 0.0367). The branch-site model that assumes positive selection at specific sites within the Phlebotominae branch identifies three sites with ω significantly higher than 1 (Additional file 14: Table S4); however, the comparison with the null model that assumes absence of positive selection is not statistically significant.

Finally, within the *fru* male-specific domain, the three-ratio model is supported better than the one- and two-ratio models, showing a relaxation of the selective constraints within the Phlebotominae branch (ω = 0.192) when compared to the mosquito branch (ω = 0.1033) and to the branch including *Drosophila, Ceratitis* and *Musca* (ω = 0.0288). Site and branch-site models do not show evidence of positive selection.

In conclusion, the analysis of the evolutionary pressure acting on the examined sex-determination genes shows evidence of strong purifying selection. However, different selective constraints act on specific branches of the *dsx* and *fru* and *tra-2* genes, whereas the evolutionary rates of the *tra* genes appear more uniform.

### Conclusions

Our results permit to hypothesize a model for the sex determination cascade of Phlebotominae sand flies as shown in figure 9, which represents the first complete and conserved sex determination cascade observed in Nematocera species. In particular, we identified all the key sex determining genes, that in figure 1 are represented by question marks and, for the first time in a Nematocera species, we identified the homolog of the *transformer* gene. In addition, our data strongly suggest the conservation of the autoregulation of the sand fly *tra* gene as observed in Brachycera and in other insect orders. The availability of the sequence of this *tra* gene will help to identify its homologs in other Nematocera species, many of them representing important vectors of human diseases. Our model needs to be confirmed by functional analyses and verified also in New World sand fly species, where the *tra* gene seems to be absent.

**Fig. 9.**
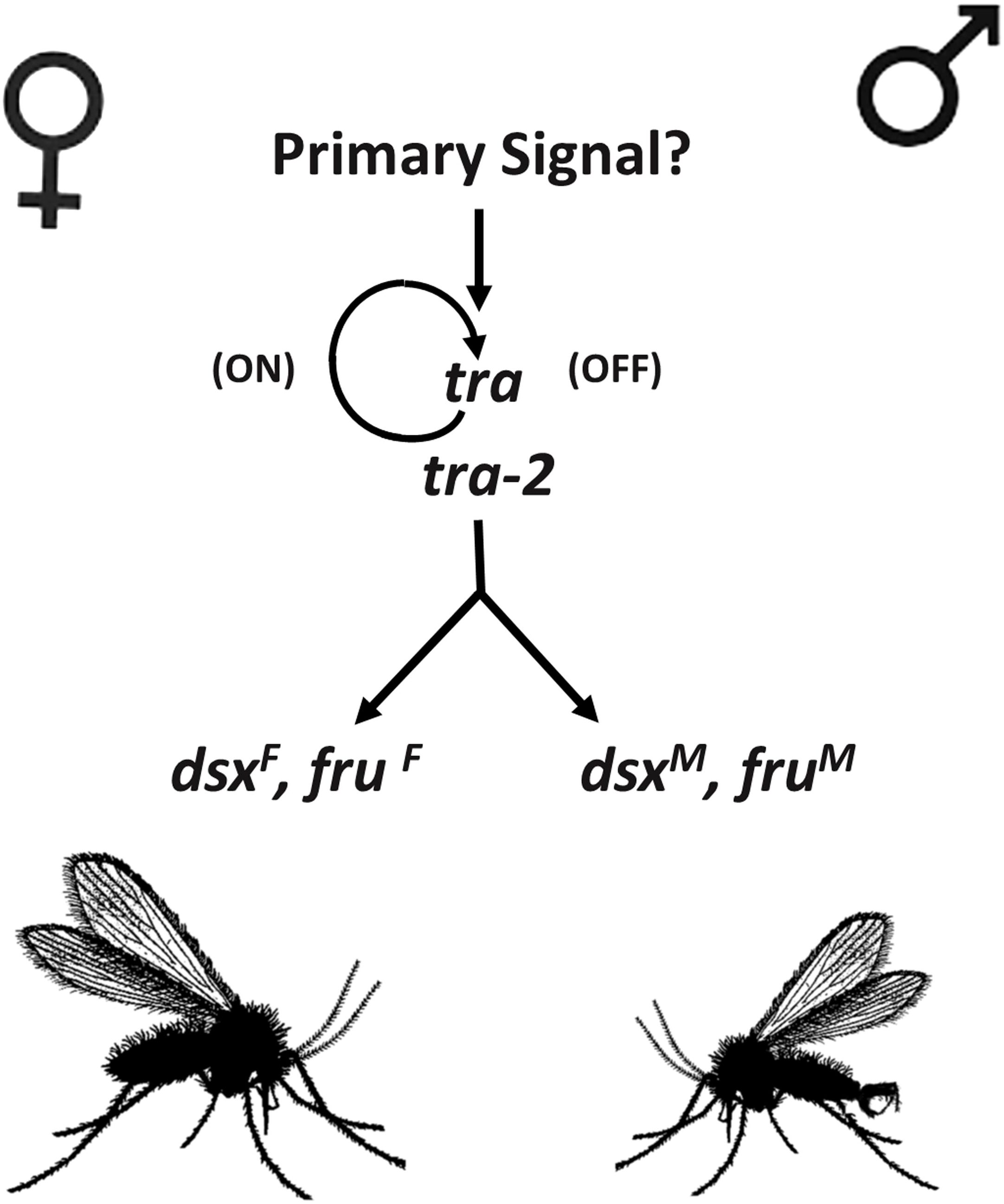
Model for sex determination in sand flies. In female embryos, a maternal *tra* mRNA or TRA protein and a maternal auxiliary TRA-2 protein led to the activation of a positive feedback autoregulative loop. The early TRA and TRA-2 proteins drive the female-specific splicing of the zygotically transcribed *tra* pre-mRNA so that new TRA protein can be produced. The newly synthesized protein controls the maintenance of *tra* autoregulation and the female-specific splicing of *dsx* and *fru* pre-mRNAs leading to female development. In male embryos, *tra* autoregulation is impaired by a male-specific factor, resulting in absence of the TRA protein, determining the male-specific splicing of the *dsx* and *fru* genes and thus inducing male development.

A further interesting question to be addressed in future is relative to the molecular nature of the primary signals of sex determination in sand flies, to date completely unknown. To this aim the *P. perniciosus* species could be an optimal starting point considering that it is the only Old World sand fly species with described heteromorphic sexual chromosomes [79]. The identification of male determining factors and sex-specific genomic loci in sand fly species could not only help to complete the understanding of sex determination mechanisms in Nematocera but also to shed light on chromosome evolution in insects [80–83].

Finally, our results open the possibility of future biotech applications to control natural populations of sand flies to reduce their impact on public health by using technologies available for other insect pests [84–86]. In particular, the *tra* gene could be utilized to produce sexing strains to be implemented for SIT-based control program [87], still missing for sand flies, while the *dsx* gene could be used to develop gene drive systems for population suppression, as recently proposed for *Anopheles* mosquitoes [68].

## Methods

### Sand flies sex determination genes cloning

The samples of *P. perniciosus* used in this study were from laboratory colonies held at the PV laboratory (Charles University, Department of Parasitology, Prague – Czech Republic) and at the LG laboratory (Istituto Superiore di Sanità, Rome – Italy). The samples of *P.papatasi* and *L. longipalpis* used in this study were from laboratory colonies held at the PV laboratory (Charles University, Department of Parasitology, Prague – Czech Republic). The sand fly colonies were reared under standard conditions as previously described [89]. Total RNA was extracted from pools of virgin males and sugar-fed females (7–10 days old) of adult *P. perniciosus, P. papatasi* and *L. longipalpis* using the PureLink® RNA Mini kit (Life Technologies) according to manufacturer’s instruction, followed by on-column PureLink® DNase (Ambion) treatment. Total RNA was resuspended in 100□μl of ddH_2_O and quantified using the NanoDrop 2000c spectrophotometer. The protein coding sequences of insect sex determining genes were used as query to perform TBLASTN search against the PERNI data set (Additional file 1: Table S1) [39]. The transcripts corresponding to the putative *P. perniciosus* orthologues (Additional file 1: Table S1) were utilized to design PCR primer pairs (see Additional file 15: supplementary methods). First-strand cDNA was synthesized from 200 ng of male and female total RNA using the EuroScript Reverse Transcriptase kit (Euroclone) with oligo-dT, in a final volume of 20 μl. To amplify the orthologue of the *fruitless* gene, cDNA was synthesized with the gene-specific primers. PCR amplifications were performed on 1 μl of 1:20 dilution of the cDNA template from adult males and females, in a final volume of 50 μl, using the Dreamtaq DNA polymerase (Thermo Fisher Scientific) or the PfuUltra HF DNA polymerase (Agilent Technologies). Appropriate annealing temperatures were adjusted to individual primer pairs (Additional file 15: Supplementary Methods). The 3’ end of the *Ppetra* cDNAs were determined with the 3 RACE System for Rapid Amplification of cDNA Ends (Invitrogen); the 5’ end of the *Ppetra* cDNA was determined with the 5’/3’ RACE kit 2nd generation (Roche). Reverse transcription was performed as recommended by the suppliers. The obtained cDNA fragments were cloned using the Strataclone PCR cloning Kit (Agilent Technologies), and the positive clones were sequenced on an ABI 310 Automated Sequencer (Applied Biosystems). cDNA sequences were deposited at the GenBank database with the following accession numbers: PpedsxM MK286442; PpedsxF MK286443; PpetraM1 MK286444; PpetraM2 MK286445; PpetraF1 MK286446; PpetraF2 MK286447; PpetraF3 MK286448; Ppetra-2A MK286449; Ppetra-2B MK286450; PpefruMA MK286451; PpefruMC MK286452; PpefruFA MK286453; PpefruFC MK286454; PpatraF MK286455.

### Developmental expression analysis

Total RNA was extracted from the different developmental stages of *P. perniciosus* (embryos, 1st, 2nd and 4th instar, and pupae) using the High Pure RNA Tissue Kit (Roche) according to manufacturer’s instruction, followed by on-column DNase treatment. First-strand cDNA was synthesized from 0.5□μg of total RNA using the First Strand cDNA Synthesis Kit for RT-PCR with both oligo-dT primers and random examers, or with the fruC-nested gene-specific primer. PCR amplifications were performed on 1 μl of 1:20 dilution of the cDNA template in a final volume of 50 μl using the EmeraldAmp PCR Master Mix (Clontech). Appropriate annealing temperatures and cycle conditions were adjusted to individual primer pairs (see supplemental methods).

### *Ppetra* genomic organization

To identify the intronic region sequence of the *P. perniciosus transformer* gene, genomic DNA was extracted from a single adult female using the NucleoSpin Tissue XS (Macherey-Nagel) according to manufacturer. PCR amplification was conducted on 10 ng of genomic DNA in a final volume of 50 μl using the primers Ppetra5utr/Ppetrastop3utr and the following thermal cycle: 95 °C for 3 min, 35 cycles of 94 °C for 30 sec, 56 °C for 30 sec, 72 °C for 2.30 min, final extension of 10 min at 72 °C. The amplification product was cloned and sequenced as described above. The *Ppetra* genomic locus sequence was deposited at the GenBank database with the following accession number: MK286466.

### Phylogeny and evolutionary analysis

Nucleotide and encoded amino acid sequence of homologs of the *Ppedsx, Ppefru, Ppetra* and *Ppetra-2* genes were downloaded from GenBank and the relative accession numbers are listed in Table S3. Amino acid sequences were aligned using MUSCLE [90]. Due to high sequence divergence, for each gene the alignments were restricted to the encoded protein regions whose alignment is not ambiguous, as follow: TRA (TRACAM domain and DIPTERA domain), TRA-2 (RRM domain and linker region), DSXF (OD1 and OD2 domains), FRUM (Male-specific N-terminal region and BTB domain). Based on their amino acid alignments, nucleotide sequences were aligned using PAL2NAL [91]. Neighbor-Joining trees were constructed on the amino acid alignments using MEGA7 [92], with 1,000 bootstrap replicates. The coding sequences of the *dsx, fru, tra* and *tra-2* homolog genes were analyzed with the CODEML program from PAML v.4.8 [93] to evaluate their evolutionary rates. Different evolutionary models were compared (branch, sites and branch-sites) to test for variation of the ratio between non-synonymous and synonymous substitution rate (ω) at specific codons in the sequences and among the branches of the trees. For each comparison, a likelihood ratio test was applied to establish which model best fits the data.

## List of abbreviations

DM: Doublesex Mab3
BTB: Broad-Complex, Tramtrack and Bric a brac
RRM: RNA Recognition Motif
TRACAM: TRA *Ceratitis*-*Apis*-*Musca*
ORF: Open Reading Frame
RACE: Rapid Amplification of cDNA END

## Declarations

### Ethics approval and consent to participate

Not applicable.

### Consent for publication

Not applicable.

### Availability of data and material

All the sequencing data produced in this work are deposited in the GenBank public database or present in the supplementary methods. *P. perniciosus* transcriptome assembly utilized in this work is freely available at http://pernibase.evosexdevo.eu while the corresponding RNA-seq raw data are available at the SRA NCBI database under the accession number PRJNA287743. Genome or transcriptome assemblies produced in this study are available upon request or reproducible using instructions present in the supplementary methods.

### Authors’ contributions

MS conceived the study. MS and VP planned the experiments. VP performed all the molecular analyses. NP helped with DNA and RNA extractions and RT-PCR analyses. MS performed all bioinformatic analysis with additional contribution of SA, VC and RS. GS suggested the search for *tra* ortholog by using TRA/TRA-2 binding site sequences. MS performed the manual curation of sex determination genes and comparative genomics analyses. SA performed the phylogeny and evolutionary analyses. PV contributed with reagents and biological samples. GB and LG maintained the *P. perniciosus* colony and collected samples. MS, SA and VC wrote the manuscript with inputs by GS, RS, PV and LG. All authors read and approved the final manuscript.

## Acknowledgements

The authors are deeply grateful to Riccardo Bianchi and Marta Marchili, Istituto Superiore di Sanità, Roma, Italy, for technical support in rearing sand flies. We deeply acknowledge the Sand Fly Genome Sequencing Consortium and the Baylor College of Medicine Human Genome Sequencing Center (BCM-HGSC) for releasing their unpublished data prior to project completion.

## Additional files

**Additional file 1: Table S1**. TBLASTN search of sex determination orthologs in the perniBASE dataset.

(XLSX 11.6 kb)

**Additional file 2: Figure S1**. Multiple sequence alignment of SXL proteins. **Figure S2**. Multiple sequence alignment of TRA-2 proteins. **Figure S3**. Multiple sequence alignment of DSX amino-terminal regions. **Figure S4**. Multiple sequence alignment of DSXF carboxy-terminal regions. **Figure S5**. Multiple sequence alignment of DSXM carboxy-terminal regions. **Figure S6.** Multiple sequence alignment of FRUM amino-terminal regions. **Figure S7**. Multiple sequence alignment of FRU proteins. **Figure S8.** Sex-lethal gene expression at adult stage in *P. perniciosus*. **Figure S9**. Multiple sequence alignment of TRA proteins. (PDF 896 kb)

**Additional file 3: Figure S10**. Manually-curated *P. perniciosus transformer* gene model. **Figure S11**. Manually-curated *P. papatasi transformer* gene model. **Figure S12**. Manually-curated *P. bergeroti transformer* gene model. **Figure S13**. Manually-curated *P. duboscqi transformer* gene model. (PDF 350 kb)

**Additional file 4: Figure S14**. Multiple sequence alignment of TRA proteins in *Phlebotomus* spp. (PDF 252 kb)

**Additional file 5: Figure S15**. Multiple alignment of tra introns in *Phlebotomus* spp. (PDF 573 kb)

**Additional file 6: Figure S16**. *tra* gene expression at adult stage in *P. papatasi*. **Figure S17**. Phylogenetic relationship, genomic structure and sex-specific splicing regulation of *transformer* orthologues in insects. (PDF 877 kb)

**Additional file 7: Figure S18**. Manually-curated P. *papatasi tra-2* partial gene model. **Figure S19**. Manually-curated *P. bergeroti tra-2* partial gene model. **Figure S20**. Manually-curated *P. duboscqi tra-2p* artial gene model. (PDF 142 kb)

**Additional file 8: Figure S21**. Comparison of genomic structures of dipteran *tra-2* genes. **Figure S22**. Multiple sequence alignment of sand fly TRA-2 proteins. (PDF 188 kb)

**Additional file 9: Figure S23**. Manually-curated *P. papatasi doublesex* gene model. **Figure S24**. Manually-curated *L. longipalpis doublesex* gene model. **Figure S25**. Manually-curated *P. papatasi fruitless* gene model. **Figure S26**. Manually-curated *L. longipalpis fruitless* partial gene model. **Figure S27**. Comparison of genomic structures of dipteran *dsx* genes. **Figure S28**. Comparison of genomic structures of dipteran *fru* genes. (PDF 284 kb)

**Additional file 10: Figure S29**. Crispr/Cas9 target sites in sand fly *dsx* genes (PDF 156 kb)

**Additional file 11: Figure S30**. Manually-curated *P. bergeroti fruitless* partial gene model. **Figure S31**. Manually-curated *P. duboscqi fruitless* partial gene model. **Figure S32**. Manually-curated *P. bergeroti doublesex* partial gene model. **Figure S33**. Manually-curated *P. duboscqi doublesex* partial gene model (PDF 143 kb)

**Additional file 12**: TRA/TRA-2 binding sites of Brachycera and Nematocera species. (XLSX 17.0 kb)

**Additional file 13**: Accession numbers and ID of the sequences used in phylogenetic and evolutionary analyses. (XLSX 17.0 kb)

**Additional file 14**: Statistics of the evolutionary analyses and comparison of different evolutionary models (XLSX 17.0 kb)

**Additional file 15**: Supplementary Methods (PDF 284 kb)

## References

1. Haag ES, Doty AV. Sex determination across evolution: Connecting the dots. PLoS Biol. 2005.

2. Zarkower D. Establishing sexual dimorphism: Conservation amidst diversity? Nat. Rev. Genet. 2001. p. 175–85.

3. Marshall Graves JA. Weird animal genomes and the evolution of vertebrate sex and sex chromosomes. Annu Rev Genet [Internet]. 2008;42:565–86. Available from: http://www.ncbi.nlm.nih.gov/pubmed/18983263

4. Verhulst EC, Van De Zande L. Insect sex determination: A cascade of mechanisms. Sex Dev. 2014;8:5–6.

5. nSáchez L. Sex-determining mechanisms in insects. Int J Dev Biol [Internet]. 2008;52:837–56. Available from: http://www.ncbi.nlm.nih.gov/pubmed/18956315

6. Gempe T, Beye M. Function and evolution of sex determination mechanisms, genes and pathways in insects. BioEssays. 2011. p. 52–60.

7. SchuDtt C, Nöthiger R. Structure, function and evolution of sex-determining systems in Dipteran insects. Development. 2000;127:667–77.

8. Erickson JW, Quintero JJ. Indirect effects of ploidy suggest X chromosome dose, not the X:A ratio, signals sex in Drosophila. PLoS Biol. 2007;5:2821–30.

9. Saccone G, Pane A, Polito LC. Sex determination in flies, fruitflies and butterflies. Genetica. 2002;116:15–23.

10. Bopp D, Saccone G, Beye M. Sex determination in insects: Variations on a common theme. Sex Dev. 2014;8:20–8.

11. Pane A, Salvemini M, Delli Bovi P, Polito C, Saccone G. The transformer gene in Ceratitis capitata provides a genetic basis for selecting and remembering the sexual fate. Development. 2002;129:3715–25.

12. Salvemini M, Robertson M, Aronson B, Atkinson P, Polito LC, Saccone G. Ceratitis capitata transformer-2 gene is required to establish and maintain the autoregulation of Cctra, the master gene for female sex determination. Int J Dev Biol [Internet]. 2009 [cited 2015 Nov 2];53:109–20. Available from: http://www.ncbi.nlm.nih.gov/pubmed/19123132

13. Verhulst EC, van de Zande L, Beukeboom LW. Insect sex determination: It all evolves around transformer. Curr. Opin. Genet. Dev. 2010. p. 376–83.

14. Katsuma S, Kiuchi T, Kawamoto M, Fujimoto T, Sahara K. Unique sex determination system in the silkworm, Bombyx mori: current status and beyond. Proc Jpn Acad, Ser B [Internet]. 2018;94:205. Available from: https://www.jstage.jst.go.jp/article/pjab/94/5/94_PJA9405B-01/_pdf/-char/ja

15. Saccone G, Salvemini M, Polito LC. The transformer gene of Ceratitis capitata: A paradigm for a conserved epigenetic master regulator of sex determination in insects. Genetica. 2011;139:99–111.

16. Ruiz MF, Milano A, Salvemini M, Eirín-López JM, Perondini ALP, Selivon D, et al. The gene transformer of anastrepha fruit flies (Diptera, tephritidae) and its evolution in insects. PLoS One [Internet]. 2007 [cited 2015 Nov 2];2:e1239. Available from: http://www.pubmedcentral.nih.gov/articlerender.fcgi?artid=2080774&tool=pmcentrez&rendertype=abstract

17. Li F, Vensko SP, Belikoff EJ, Scott MJ. Conservation and Sex-Specific Splicing of the transformer Gene in the Calliphorids Cochliomyia hominivorax, Cochliomyia macellaria and Lucilia sericata. PLoS One. 2013;8:1–14.

18. Hediger M, Henggeler C, Meier N, Perez R, Saccone G, Bopp D. Molecular characterization of the key switch F provides a basis for understanding the rapid divergence of the sex-determining pathway in the housefly. Genetics. 2010;184:155–70.

19. Concha C, Scott MJ. Sexual development in Lucilia cuprina (Diptera, Calliphoridae) is controlled by the transformer gene. Genetics. 2009;182:785–98.

20. Luo Y, Zhao S, Li J, Li P, Yan R. Isolation and Molecular Characterization of the Transformer Gene From Bactrocera cucurbitae (Diptera: Tephritidae). J Insect Sci. 2017;17.

21. Laohakieat K, Aketarawong N, Isasawin S, Thitamadee S, Thanaphum S. The study of the transformer gene from Bactrocera dorsalis and B. correcta with putative core promoter regions. BMC Genet. 2016;17.

22. Salvemini M, D’Amato R, Petrella V, Aceto S, Nimmo D, Neira M, et al. The Orthologue of the Fruitfly Sex Behaviour Gene Fruitless in the Mosquito Aedes aegypti: Evolution of Genomic Organisation and Alternative Splicing. PLoS One. 2013;8.

23. Salvemini M, Mauro U, Lombardo F, Milano A, Zazzaro V, Arcà B, et al. Genomic organization and splicing evolution of the doublesex gene, a Drosophila regulator of sexual differentiation, in the dengue and yellow fever mosquito Aedes aegypti. BMC Evol Biol. 2011;11:41.

24. Gailey DA, Billeter JC, Liu JH, Bauzon F, Allendorfer JB, Goodwin SF. Functional conservation of the fruitless male sex-determination gene across 250 Myr of insect evolution. Mol Biol Evol. 2006;23:633–43.

25. Scali C. Identification of sex-specific transcripts of the Anopheles gambiae doublesex gene. J Exp Biol [Internet]. 2005;208:3701–9. Available from: http://jeb.biologists.org/cgi/doi/10.1242/jeb.01819

26. Criscione F, Qi Y, Tu Z. GUY1 confers complete female lethality and is a strong candidate for a male-determining factor in anopheles Stephensi. Elife. 2016;5.

27. Krzywinska E, Dennison NJ, Lycett GJ, Krzywinski J. A maleness gene in the malaria mosquito Anopheles gambiae. Science (80-). 2016;353:67–9.

28. Hall AB, Basu S, Jiang X, Qi Y, Timoshevskiy VA, Biedler JK, et al. A male-determining factor in the mosquito Aedes aegypti. Science (80-). 2015;348:1268–70.

29. Biedler JK, Tu Z. Sex Determination in Mosquitoes. Adv In Insect Phys. 2016. p. 37–66.

30. Maroli M, Feliciangeli MD, Bichaud L, Charrel RN, Gradoni L. Phlebotomine sandflies and the spreading of leishmaniases and other diseases of public health concern. Med. Vet. Entomol. 2013. p. 123–47.

31. Young DG, Duncan MA. Guide of the identification and geografic distribution of Lutzomyia Sand Flies in Mexico, West Indies, Central and South America (Diptera: Psychodidae). Mem Am Entomol Inst. 1994;54:1–881.

32. WHO WHO. Control of the Leishmaniases. World Health Organization, Geneva. Tech Rep Ser. 2010.

33. Alvar J, Vélez ID, Bern C, Herrero M, Desjeux P, Cano J, et al. Leishmaniasis worldwide and global estimates of its incidence. PLoS One. 2012.

34. Depaquit J, Grandadam M, Fouque F, Andry PE, Peyrefitte C. Arthropod-borne viruses transmitted by Phlebotomine sandflies in Europe: a review. Euro Surveill. 2010. p. 19507.

35. Volff JN, Zarkower D, Bardwell VJ, Schartl M. Evolutionary Dynamics of the DM Domain Gene Family in Metazoans. J Mol Evol. 2003.

36. Perez-Torrado R, Yamada D, Defossez PA. Born to bind: The BTB protein-protein interaction domain. BioEssays. 2006. p. 1194–202.

37. Cléry A, Blatter M, Allain FHT. RNA recognition motifs: boring? Not quite. Curr. Opin. Struct. Biol. 2008. p. 290–8.

38. Geuverink E, Beukeboom LW. Phylogenetic distribution and evolutionary dynamics of the sex determination genes doublesex and transformer in insects. Sex Dev. 2014;8:38–49.

39. Petrella V, Aceto S, Musacchia F, Colonna V, Robinson M, Benes V, et al. De novo assembly and sex-specific transcriptome profiling in the sand fly Phlebotomus perniciosus (Diptera, Phlebotominae), a major Old World vector of Leishmania infantum. BMC Genomics [Internet]. 2015 [cited 2015 Nov 2];16:847. Available from: http://www.pubmedcentral.nih.gov/articlerender.fcgi?artid=4619268&tool=pmcentrez&rendertype=abstract

40. McAllister BF, McVean GAT. Neutral evolution of the sex-determining gene transformer in Drosophila. Genetics. 2000;154:1711–20.

41. Kulathinal RJ, Skwarek L, Morton RA, Singh RS. Rapid evolution of the sex-determining gene, transformer: Structural diversity and rate heterogeneity among sibling species of drosophila. Mol Biol Evol. 2003;20:441–52.

42. Serna E, Gorab E, Ruiz MF, Goday C, Eirín-López JM, Sánchez L. The gene Sex-lethal of the sciaridae family (order diptera, suborder nematocera) and its phylogeny in dipteran insects. Genetics. 2004;168:907–21.

43. Traut W, Niimi T, Ikeo K, Sahara K. Phylogeny of the sex-determining gene *Sex-lethal* in insects. Genome [Internet]. 2006;49:254–62. Available from: http://article.pubs.nrc-cnrc.gc.ca/ppv/RPViewDoc?issn=1480-3321&volume=49&issue=3&startPage=254&ab=y

44. Meise M, Hilfiker-Kleiner D, Dübendorfer a, Brunner C, Nöthiger R, Bopp D. Sex-lethal, the master sex-determining gene in Drosophila, is not sex-specifically regulated in Musca domestica. Development. 1998;125:1487–94.

45. Saccone G, Peluso I, Artiaco D, Giordano E, Bopp D, Polito LC. The Ceratitis capitata homologue of the Drosophila sex-determining gene sex-lethal is structurally conserved, but not sex-specifically regulated. Development. 1998;125:1495–500.

46. Graham P, Penn JKM, Schedl P. Masters change, slaves remain. BioEssays. 2003. p. 1–4.

47. Xie W, Guo L, Jiao X, Yang N, Yang X, Wu Q, et al. Transcriptomic dissection of sexual differences in Bemisia tabaci, an invasive agricultural pest worldwide. Sci Rep [Internet]. 2014;4:4088. Available from: http://www.pubmedcentral.nih.gov/articlerender.fcgi?artid=3924218&tool=pmcentrez&rendertype=abstract

48. Pane A, Salvemini M, Delli Bovi P, Polito C, Saccone G. The transformer gene in Ceratitis capitata provides a genetic basis for selecting and remembering the sexual fate. Development. 2002;129.

49. Salvemini M, Robertson M, Aronson B, Atkinson P, Polito LC, Saccone G. Ceratitis capitata transformer-2 gene is required to establish and maintain the autoregulation of Cctra, the master gene for female sex determination. Int J Dev Biol. 2009;53:109–20.

50. Burghardt G, Hediger M, Siegenthaler C, Moser M, Dübendorfer A, Bopp D. The transformer2 gene in Musca domestica is required for selecting and maintaining the female pathway of development. Dev Genes Evol. 2005;215:165–76.

51. Burtis KC, Baker BS. Drosophila doublesex gene controls somatic sexual differentiation by producing alternatively spliced mRNAs encoding related sex-specific polypeptides. Cell. 1989;56:997–1010.

52. Saccone G, Salvemini M, Pane A, Polito LC. Masculinization of XX Drosophila transgenic flies expressing the Ceratitis capitata DoublesexM isoform. Int J Dev Biol [Internet]. 2008 [cited 2015 Nov 2];52:1051–7. Available from: http://www.ncbi.nlm.nih.gov/pubmed/18956338

53. Salikhov K, Sacomoto G, Kucherov G. Using cascading bloom filters to improve the memory usage for de Brujin graphs. Lect Notes Comput Sci (including Subser Lect Notes Artif Intell Lect Notes Bioinformatics). 2013. p. 364–76.

54. Grabherr MG, Haas BJ, Yassour M, Levin JZ, Thompson DA, Amit I, et al. Full-length transcriptome assembly from RNA-Seq data without a reference genome. Nat Biotechnol. 2011;29:644–52.

55. Haas BJ, Papanicolaou A, Yassour M, Grabherr M, Philip D, Bowden J, et al. reference generation and analysis with Trinity. Nat Protoc [Internet]. 2014;8:1–43. Available from: http://www.pubmedcentral.nih.gov/articlerender.fcgi?artid=3875132&tool=pmcentrez&rendertype=abstract

56. Belote JM, Baker BS. Sex determination in Drosophila melanogaster: analysis of transformer-2, a sex-transforming locus. Proc Natl Acad Sci U S A [Internet]. 1982;79:1568–72. Available from:http://www.pubmedcentral.nih.gov/articlerender.fcgi?artid=346016&tool=pmcentrez&rendertype=abstract

57. Goralski TJ, Edström JE, Baker BS. The sex determination locus transformer-2 of Drosophila encodes a polypeptide with similarity to RNA binding proteins. Cell. 1989;56:1011–8.

58. Chandler D, McGuffin ME, Piskur J, Yao J, Baker BS, Mattox W. Evolutionary conservation of regulatory strategies for the sex determination factor transformer-2. Mol Cell Biol [Internet]. 1997;17:2908–19. Available from: http://www.pubmedcentral.nih.gov/articlerender.fcgi?artid=232143&tool=pmcentrez&rendertype=abstract

59. Gomulski LM, Dimopoulos G, Xi Z, Soares MB, Bonaldo MF, Malacrida AR, et al. Gene discovery in an invasive tephritid model pest species, the Mediterranean fruit fly, Ceratitis capitata. BMC Genomics. 2008;9:243.

60. Sarno F, Ruiz MF, Eirín-López JM, Perondini AL, Selivon D, Sánchez L. The gene transformer-2 of Anastrepha fruit flies (Diptera, Tephritidae) and its evolution in insects. BMC Evol Biol [Internet]. 2010;10:140. Available from: http://bmcevolbiol.biomedcentral.com/articles/10.1186/1471-2148-10-140

61. Martín I, Ruiz MF, Sánchez L. The gene transformer-2 of Sciara (Diptera, Nematocera) and its effect on Drosophila sexual development. BMC Dev Biol [Internet]. 2011;11:19. Available from: http://bmcdevbiol.biomedcentral.com/articles/10.1186/1471-213X-11-19

62. Dauwalder B, Amaya-Manzanares F, Mattox W. A human homologue of the Drosophila sex determination factor transformer-2 has conserved splicing regulatory functions. Proc Natl Acad Sci [Internet]. 1996;93:9004–9. Available from: http://www.pnas.org/cgi/doi/10.1073/pnas.93.17.9004

63. Liu G, Wu Q, Li J, Zhang G, Wan F. RNAi-mediated knock-down of transformer and transformer 2 to generate male-only progeny in the oriental fruit fly, Bactrocera dorsalis (Hendel). PLoS One. 2015;10.

64. Salvemini M, D’Amato R, Petrella V, Aceto S, Nimmo D, Neira M, et al. The Orthologue of the Fruitfly Sex Behaviour Gene Fruitless in the Mosquito Aedes aegypti: Evolution of Genomic Organisation and Alternative Splicing. PLoS One. 2013;8.

65. Hoang KP, Teo TM, Ho TX, Le VS. Mechanisms of sex determination and transmission ratio distortion in Aedes aegypti. Parasit Vectors [Internet]. 2016;9:49. Available from: http://www.parasitesandvectors.com/content/9/1/49

66. Hediger M, Burghardt G, Siegenthaler C, Buser N, Hilfiker-Kleiner D, Dübendorfer A, et al. Sex determination in *Drosophila melanogaster* and *Musca domestica* converges at the level of the terminal regulator *doublesex*. Dev Genes Evol. 2004;214:29–42.

67. Kuhn S, Sievert V, Traut W. The sex-determining gene *doublesex* in the fly *Megaselia scalaris*: Conserved structure and sex-specific splicing. Genome [Internet]. 2000;43:1011–20. Available from: http://www.nrc.ca/cgi-bin/cisti/journals/rp/rp2_abst_e?gen_g00-078_43_ns_nf_gen43-00

68. Kyrou K, Hammond AM, Galizi R, Kranjc N, Burt A, Beaghton AK, et al. A CRISPR-Cas9 gene drive targeting doublesex causes complete population suppression in caged Anopheles gambiae mosquitoes. Nat Biotechnol. 2018;36:1062–6.

69. Ryner LC, Baker BS. Regulation of doublesex pre-mRNA processing occurs by 3’-splice site activation. Genes Dev. 1991;5:2071–85.

70. Baker BS, Heinrichs V, Ryner LC. Regulation of Sex-Specific Selection of fruitless 5’ Splice Sites by transformer and transformer-2. Mol Cell Biol. 1998;18:450–8.

71. Hoshijima K, Inoue K, Higuchi I, Sakamoto H, Shimura Y. Control of doublesex alternative splicing by transformer and transformer-2 in Drosophila. Science (80-). 1991;252:833–6.

72. Salvemini M, Polito C, Saccone G. Fruitless alternative splicing and sex behaviour in insects: an ancient and unforgettable love story? J Genet [Internet]. 2010 [cited 2015 Nov 2];89:287–99. Available from: http://www.ncbi.nlm.nih.gov/pubmed/20876995

73. Ruiz MF, Alvarez M, Eirín-López JM, Sarno F, Kremer L, Barbero JL, et al. An unusual role for doublesex in sex determination in the dipteran Sciara. Genetics. 2015;200:1181–99.

74. Price DC, Egizi A, Fonseca DM. Characterization of the doublesex gene within the Culex pipiens complex suggests regulatory plasticity at the base of the mosquito sex determination cascade. BMC Evol Biol. 2015;15.

75. Salvemini M, Mauro U, Lombardo F, Milano A, Zazzaro V, Arcà B, et al. Genomic organization and splicing evolution of the doublesex gene, a Drosophila regulator of sexual differentiation, in the dengue and yellow fever mosquito Aedes aegypti. BMC Evol Biol. 2011;11.

76. Yang Z, Nielsen R. Synonymous and nonsynonymous rate variation in nuclear genes of mammals. J Mol Evol. 1998;46:409–18.

77. Yang Z, Wong WSW, Nielsen R. Bayes empirical Bayes inference of amino acid sites under positive selection. Mol Biol Evol. 2005;22:1107–18.

78. Zhang J, Nielsen R, Yang Z. Evaluation of an improved branch-site likelihood method for detecting positive selection at the molecular level. Mol Biol Evol. 2005;22:2472–9.

79. Kreutzer RD, Modi GB, Tesh RB, Young DG. Brain cell karyotypes of six species of New and Old World sand flies (Diptera: Psychodidae). J Med Entomol. 1987;24:609–12.

80. Cortez D, Marin R, Toledo-Flores D, Froidevaux L, Liechti A, Waters PD, et al. Origins and functional evolution of Y chromosomes across mammals. Nature [Internet]. 2014;508:488–93. Available from: http://www.ncbi.nlm.nih.gov/pubmed/24759410

81. Blackmon H, Ross L, Bachtrog D. Sex Determination, Sex Chromosomes, and Karyotype Evolution in Insects. J Hered [Internet]. 2016;esw047. Available from: http://www.ncbi.nlm.nih.gov/pubmed/27543823

82. Kaiser VB, Bachtrog D. Evolution of Sex Chromosomes in Insects. Annu Rev Genet [Internet]. 2010;44:91–112. Available from: http://www.annualreviews.org/doi/10.1146/annurev-genet-102209-163600

83. Vyskot B. Y Chromosome Evolution. Brenner’s Encycl Genet Second Ed. 2013. p. 372–5.

84. Papathanos P a, Bossin HC, Benedict MQ, Catteruccia F, Malcolm C a, Alphey L, et al. Sex separation strategies: past experience and new approaches. Malar J. 2009;8 Suppl 2:S5.

85. Hendrichs J, Robinson A. Sterile Insect Technique. Encycl Insects. 2009. p. 953–7.

86. Hendrichs JP. Use of the sterile insect technique against key insect pests. Sustain Dev Int. 2000;75–9.

87. Saccone G, Pane A, De Simone A, Salvemini M, Milano A, Annunziata L, et al. New sexing strains for Mediterranean fruit fly Ceratitis capitata: Transforming females into males. Area-Wide Control Insect Pests From Res. to F. Implement. 2007.

88. De Murtas ID, Cirio U, Guerrieri G, Enkerlin D. An experiment to control the Mediterranean fruit fly on the Island of Procida by the sterile-insect technique. Sterile-male Tech Control fruit flies STI/PUB/276 [Internet]. 1970. p. 59–70. Available from: file:///Y:/3772.pdf

89. Volf P, Volfova V. Establishment and maintenance of sand fly olonies. J Vector Ecol. 2011;36.

90. Edgar RC. MUSCLE: Multiple sequence alignment with high accuracy and high throughput. Nucleic Acids Res. 2004;32:1792–7.

91. Suyama M, Torrents D, Bork P. PAL2NAL: Robust conversion of protein sequence alignments into the corresponding codon alignments. Nucleic Acids Res. 2006;34.

92. Kumar S, Stecher G, Tamura K. MEGA7: Molecular Evolutionary Genetics Analysis Version 7.0 for Bigger Datasets. Mol Biol Evol. 2016;33:1870–4.

93. Yang Z. PAML: a program package for phylogenetic analysis by maximum likelihood. Comput Appl Biosci [Internet]. 1997;13:555–6. Available from: http://www.ncbi.nlm.nih.gov/pubmed/9367129

