## Additional_file_2_Figures_S1_S9 for "Genomics and transcriptomics to unravel sex determination pathway and its evolution in sand flies"

```
DmSxl_NP_727160         MYGNNNPGSNNNNGGYPPYGYNN-KSSGGRGFGMSHSLPSGMDTEFSFPS-------SSS      52
CcSxl_NM_001279407      MYGNM------NNGGHAPYGYNGYRPSGGRMWGMSHSLPSGMDTDFTSSYPGPSMN---R      51
MdSxl_NM_001287535      MYGQN----VRNVSYYPPYGYNGYKQSGERMWRMSHSLPSGMDTDFTSSYPGPSAMNPRG      56
AgaSxl_BN000856         MYSKM-----------NGYPSSSFRSSSGRLWGMSHSLPSGMDADF------SSYS---R      40
PpeSxl                  MYNKM-----------NGYA--PFRSSSGRMWGMSHSLPSGMSHYAYQDTDFSLMY--NT      45
                        **.:              *     : *. * : *********.

DmSxl_NP_727160         RRGYNDFPGCGGSGGNGGSANNL-GGGNMCHLPPM-----ASNNSLNNLCGLSLGSGGSD      106
CcSxl_NM_001279407      RGGYNDFSGGGGGGGG--------TMGSMNNMVNA-----ASTNSLNCG--G---GGGRD      93
MdSxl_NM_001287535      RGGYNDFSGGG----S--------AMGSMCNMAPA-----QSTNSLNSG--G---DGGG-      93
AgaSxl_BN000856         PRLYGDMPSSSTFGGGNPSTPHHHHHHHHNHLSQPASHHHTSSSSTSSLTGMNFGP----      96
PpeSxl                  RRGGMNFPASSTFGAV---------------------GHASSSSTS-------------      70
                             :: . .                              *..* .

                                  RRM1
DmSxl_NP_727160         DLMNDPRASNTNLIVNYLPQDMTDRELYALFRAIGPINTCRIMRDYK-TGYSFGYAFVDF      165
CcSxl_NM_001279407      --GHGGGSNGTNLIVNYLPQDMTDRELYALFRTIGPINTCRIMRDYK-TGYSFGYAFVDF      150
MdSxl_NM_001287535      --GDTQAVNGTNLIVNYLPQDMTDRELYALFRTCGPINTCRIMKDYK-TGYSFGYAFVDF      150
AgaSxl_BN000856         ---GSSGTAGTNLIVNYLPQDMTEREMYSMFSAMGPIESCRLMRDLKQTGYSYGFGFVNY      153
PpeSxl                  ---SLSGVAGTNLIINYLPQDMSEREMYSLFSTMGPIESCKIMRDMK-TGYSYGFGFINY      126
                                 .****:*******::**:*::* : ***::*::*:* * ****:*:.*:::

                                                                  RRM2
DmSxl_NP_727160         TSEMDSQRAIKVLNGITVRNKRLKVSYARPGGESIKDTNLYVTNLPRTITDDQLDTIFGK      225
CcSxl_NM_001279407      AAETDSQRAIKSLNGITVRNKRLKVSYARPGGESIKDTNLYVTNLPRTITDDQLDTIFGK      210
MdSxl_NM_001287535      ASEIDAQNAIKTVNGITVRNKRLKVSYARPGGESIKDTNLYVTNLPRTITDDELEKIFGK      210
AgaSxl_BN000856         LNEEAAQRAIKCLNGYPLRNKRLKVSYARPQSDDIKETNLYITNLPRTITEEQLDIIFGK      213
PpeSxl                  STEEAASRAIKCLNGLTVRNKRLKVSYARPSGDDLKETNLYVTNLPRTITEEMLDVIFGK      186
                          *  :..*** :**  :************ .:.:*:****:********:: *: ****

DmSxl_NP_727160         YGSIVQKNILRDKLTGRPRGVAFVRYNKREEAQEAISALNNVIPEGGSQPLSVRLAEEHG      285
CcSxl_NM_001279407      YGMIVQKNILRDKLTGKPRGVAFVRFNKREEAQEAISALNNVIPEGASQPLTVRLAEEHG      270
MdSxl_NM_001287535      YGNIVQKNILRDKLTGRPRGVAFVRFNKREEAQEAISALNNVIPEGASQPLTVRLAEEHG      270
AgaSxl_BN000856         YGTIVQKNILRDKLTGQPRGVAFVRFNKREEAQEAISALNNVIPQGGNQPLIVRVAEDHG      273
PpeSxl                  YGTIVQKNILRDKITGQPRGVAFVRFNKREEAQEAISALNNVIPQGGNQPLTVRVAEEHG      246
                        ** **********:**:********:******************:*..*** **:**:**

DmSxl_NP_727160         KAKAAHFMSQMGVVPANV------------PPPPPQPPAHMAAAFNMMHRGRSIKSQQRF      333
CcSxl_NM_001279407      KAKAQHYMSQLGLIGGGGGGGGGGGGGGGGMGGPPPPP-MNMGYNNMVHRGRQN--KSRF      327
MdSxl_NM_001287535      KMKAHHFMNQLGMG----------------PPAAPIPA-AGPGYNNMVHRGRQN--KMRN      311
AgaSxl_BN000856         RAKAALYVP---------------------------------SYNSIVHNNRGRVRMRNT      300
PpeSxl                  KAKGALYLP---------------------------------TYNSVMHRGRQRMRMNMN      273
                        : *.  ::                                     .::*..*

DmSxl_NP_727160         QNSHPYFDAKKFI 346
CcSxl_NM_001279407      QKMHPYHNAQKFI 340
MdSxl_NM_001287535      HKVHPYHNPQKFI 324
AgaSxl_BN000856         ----PY------- 302
PpeSxl                  NRTLPY------- 279
```

**Figure S1. Multiple sequence alignment of SXL proteins**. Sequence alignment of SXL proteins of *Drosophila melanogaster*, *Ceratitis capitata*, *Musca domestica*, *Anopheles gambiae* and *Phlebotomus perniciosus*. The two RNA-binding domains (RRM1 and RRM2) are highlighted in grey. Gaps were introduced in the alignment to maximize similarity. The protein sequences alignment was performed using the Clustal-Omega software (1.2.4).

```
                                            RS domain 1
DmTRA-2_NM_057416       MDREPLSSGRLHCSARYKHKRSASSSSAGTTSSGHKDRRSDYDYCGSRRHQRSSSRRRSR   60
CcTRA-2_EU999754        MS-------------PRSRSRSISARRSYTKS--PAR-----RSNGRRRH---SREKVYK   37
MdTRA-2_AY847518        MG-------------PRSRSRSRS----Y--S--PED-----R-RGRRRS---YSRDRYS   30
PpeTRA-2                MTSDQ----------KNSMSRSPSEGRSYILKGGSSR-----ENNGYRRSDRSRSRRRYR   45
                        *                . .** *       .             * **      .

DmTRA-2_NM_057416       SRSS-------SESPPPEPRHRSGR-------------------------SSRDRERMH   87
CcTRA-2_EU999754        SRSRSI--SRHPPSPPPPPTGRGGVRCSDASQSSSTSLSPRQGRRMSRSRSRSPYDKR-R   94
MdTRA-2_AY847518        -SPVSK--RRQPPSPPPLPPSRRRSYSRSESPGYNRRNYSSHGNRSKPPRSRSPYDRNYR   87
PpeTRA-2                SRSRSHSQRRHRSRY----------------SGSP----VRNDRKHSHSRSPHSSRRRHV   85
                                                                        ..

                                   RRM
DmTRA-2_NM_057416       KSREHPQASRCIGVFGLNTNTSQHKVRELFNKYGPIERIQMVIDAQTQRSRGFCFIYFEK   147
CcTRA-2_EU999754        GNREKPVQNRCIGVFGLSVYTTQQKIRDIFSRFGPIERIQVVIDAQTGRSRGFCFIYYDD   154
MdTRA-2_AY847518        NNREKPSPCRCLGVFGLSVHTTQQQIREIFSKYGPIERIQVVVDAQTGRSRGFCFIYYKH   147
PpeTRA-2                GSRDHPQQSRCLGVFGLSLITTKQQINDIFSKFGRIKQIQIIMDAKTGRSRGFCFVYFAD   145
                         .*::*   **:*****.  *:::::.::*.::* *::**:::**:* *******:*: .
                                    RNP-1                                RNP-2

                                               Linker Region           RS domain 2
DmTRA-2_NM_057416       LSDARAAKDSCSGIEVDGRRIRVDFSITQRAHTPTPGVYLGRQPRGKAPRSFSPRRGRRV   207
CcTRA-2_EU999754        IADAKAAKDACSGMEIDDRRIRVDYSTTQRPHTPTPGVYMGRHTRREREYNDRYRDD-YR   213
MdTRA-2_AY847518        LADAEVARDQCCGQEVDGRRIRVAYSITERPHSPTPGVYRGRSTRSLSQRNRSQ------   201
PpeTRA-2                ADDAKVARENCSGMEIDGRRIRVDYSITERAHTPTPGKYMGVSTGRGREHNRSNRDDDYY   205
                          **..*:: *.* *:*.***** :* *:* *:**** * *         .

DmTRA-2_NM_057416       YHDRSASPYDNYRDRYDYRNDRYDRNLRRS--PSRNRYTRNRSYSRSRSPQLRRTSSRY   264
CcTRA-2_EU999754        PRRRSGSPFKNRNNYRNDRRRRYDRSRSRSYSPRRARY---------------------   251
MdTRA-2_AY847518        -RRHSPS------PYSRSNRDRYERSRSRSHSPRRYRY---------------------   232
PpeTRA-2                YERESRSRYIERRSPSPYR-RKRYRSRSRSYSPRIRRYR--------------------   243
                         . .* *           .  :  *.  **  *   **
```

**Figure S2. Multiple sequence alignment of TRA-2 proteins.** Sequence alignment of TRA-2 proteins of *D. melanogaster* (DmTRA-2), *Ceratitis capitata* (CcTRA-2), *Musca domestica* (MdTRA-2) and *Phlebotomus perniciosus*. The RNA recognition motif (RRM) is highlighted in black, RS domains are in light grey, and the linker region is in dark grey. RNP-1 and RNP-2 indicate the positions of two ribonucleoprotein identifier sequences, which are highly conserved between RRM proteins. Note the high number of identical amino acids within the RRM domain, whereas both RS domains are less conserved but are abundant in arginine and serine. Gaps were introduced in the alignment to maximize similarity. The protein sequences alignment was performed using the Clustal-Omega software (1.2.4).

```
                                                                 OD1
DmDSX_NP_731198.1      MVSEENWNSDTMSDSDMIDSKNDVCGGASSSSGSSISPRTPPNCARCRNHGLKITLKGHK   60
CcDSX_AAN63598.2       MVSEDNWNSDTMSDSDIHDSKADACGGASSSSGSSISPRTPPNCARCRNHGLKITLKGHK   60
AngDSX_AAZ78363.1      MVSQDRWAE-AMSDSGYDS-R-TDGNGASSSCNNSLNPRTPPNCARCRNHGLKIGLKGHK   57
AeaDSX_ABD96571.1      MVSQDRWMV-KMSEAGYDN-R-ADGSGASS---SSLNPRTPPNCARCRNHGHKIGLKGHK   54
PpeDSX                 MVSEHNTWSDVMTNSGPTDSKDEMCRGASN----SVNPRTPPNCARCRNHGLKIALKGHK   56
                       ***:..     *:::.  . :     ***.    *:.************** ** *****

DmDSX_NP_731198.1      RYCKFRYCTCEKCRLTADRQRVMALQTALRRAQAQDEQRALHMHEVPPANPAATTLLSHH   120
CcDSX_AAN63598.2       RYCKFRYCTCEKCRLTADRQRVMALQTALRRAQAQDEQRVLQIHEVPPGVHAPAALLNHH   120
AngDSX_AAZ78363.1      RYCKYRTCHCEKCCLTAERQRVMALQTALRRAQTQDEQRALNEGEVPPEPVANIHIPKLS   117
AeaDSX_ABD96571.1      RYCKYRNCTCEKCCLTAERQRVMALQTALRRAQTQDEQRLLVDGEVPAEPVHSLQIPKLS   114
PpeDSX                 RYCKFRICSCEKCRLTAERQRVMALQTALRRAQAQDEARHISADEVPPTPPPALAICPIT   116
                       ****:* * **** ***:***************:*** * :   ***        :

DmDSX_NP_731198.1      HHVAAPAHVHAHHVHAHHAHGGHHSHHGHVLHHQQAAAAAAAAPSAPASHLGGSSTAASS   180
CcDSX_AAN63598.2       HLHHHHHLNPN--------------------HHAT-------------------------   135
AngDSX_AAZ78363.1      ELKDLKH---N------------------MIHNSQTRSFDCDSSTGSMASAPGT------   150
AeaDSX_ABD96571.1      DLKEMIH---N------------------S---QQRSLIDCDSSTGSMNSTPGS------   144
PpeDSX                 TPK---------------------------------------------------------   119

DmDSX_NP_731198.1      IHGHAHAHHVHMAAAAAASVAQHQHQSHPHSHHHHHQNHHQHPHQQPATQTALRSPPHSD   240
CcDSX_AAN63598.2       -----------AAAAAAA-AAAH-----------------------HHITTAIRSPPHAE   160
AngDSX_AAZ78363.1      -----------------SSVPLTIHRRSPGVP----------------------------   165
AeaDSX_ABD96571.1      -----------------SLVTLSQHRRSPCSA----------------------------   159
PpeDSX                 ----------------------SQ------------------------------------   121

DmDSX_NP_731198.1      HGGSVGPATSSSGGGAPSSSNAAAATSSNGSSGGGGGGGGGSSGGGAGGGRSSGTSVITS   300
CcDSX_AAN63598.2       LGS-----------------------------------GGGGLAGGIGSAITSVPVSAPP   185
AngDSX_AAZ78363.1      ------------------------------------------------------------   165
AeaDSX_ABD96571.1      ------------------------------------------------------------   159
PpeDSX                 ---------------------------------------------------------CDS   124

                                                                          OD2
DmDSX_NP_731198.1      ADHHMTTVPTPAQSLEGSCDSSSPSPSSTSGAAILPISVS----VN-RKNGANVPLGQDV   355
CcDSX_AAN63598.2       PEHHMTTVPTPAQSLEGSSDTSSPSPSSTSGAAL-PISVVGRKPSL-HPNGVHMPLAQDV   243
AngDSX_AAZ78363.1      --HH-VA---EPQHLGATHSCVSPEPV-----NLL---------------------PDDE   193
AeaDSX_ABD96571.1      --AS-VHPSEAQQNVAGNHSSGTPEPG-----NMVPVGPHMRV----QHHG-PDSGTDDE   206
PpeDSX                 EPPAPITVPLPQRSHEGSCQSSSTTPCSTGEPIPVPSSRKL-APVNPTIRNPAASQSSEQ   183
                                   :   .. .  :  *                               .:

DmDSX_NP_731198.1      FLDYCQKLLEKFRYPWELMPLMYVILKDADANIEEASRRIEE  397
CcDSX_AAN63598.2       FLEHCQKLLEKFRYPWEMMPLMYVILKDAGADIEEASRRIEE  285
AngDSX_AAZ78363.1      LVKRAQWLLEKLGYPWEMMPLMYVILKSADGDVQKAHQRIDE  235
AeaDSX_ABD96571.1      LVKRSQWLLEKLRYPWEMMPLMYVILKGADGDVNKARQRIDE  248
PpeDSX                 FFDNCQKLLDKFNYPWELMPLMYVILENANVDVREAARRIEE  225
                       :.. .* **:*: ****:********:.*. ::.:* :**:*
```

**Figure S3. Multiple sequence alignment of DSX amino-terminal region.** Sequence alignment of non-sex-specific DSX amino-terminal regions of *D. melanogaster* (DmDSX), *Ceratitis capitata* (CcDSX), *Anopheles gambiae* (AngDSX), *Aedes aegypti* (AeaDSX) and *Phlebotomus perniciosus*. The amino-terminal DNA binding DM domain (OD1) and oligomerization domain (OD2) are boxed in grey. The six amino acids boxed in black indicates the positions whose replacement has been shown to abolish DNA-binding activity in *D. melanogaster*; the three amino acids in bold letters indicated residues specific for the DSX DM domain. Gaps were introduced in the alignment to maximize similarity. The protein sequences alignment was performed using the Clustal-Omega software (1.2.4).

```
                            OD2 - Female-specific portion
DmDSXF_NP_731198.1          GQYVVNEYSRQHNLNIYDGGELRNTTRQCG    30
CcDSXF_AAN63598.2           GQHVVNEYSRQHNLNIFDGGELRSTTRQCG    30
AngDSXF_AAZ78363.1          GQAVVNEYSRLHNLNMFDGVELRNTTRQSG    30
AeaDSXF_ABD96571.1          GQAVVNEYSRLHNLNMFDGVELRSTTRQSG    30
PpeDSXF                     GQCVVNEYTRMHNLNMYDGAELRGSTRQCG    30
                            ** *****:* ****::** ***.:***.*
```

**Figure S4. Multiple sequence alignment of DSXF carboxy-terminal regions.** Sequence alignment of female-specific carboxy-terminal region of DSX proteins of *D. melanogaster* (DmDSX), *Ceratitis capitata* (CcDSX), *Anopheles gambiae* (AngDSX), *Aedes aegypti* (AeaDSX) and *Phlebotomus perniciosus*. Gaps were introduced in the alignment to maximize similarity. The protein sequences alignment was performed using the Clustal-Omega software (1.2.4).

```
DmDSXM_NP_731197.1      ------------------------------------------------------------  0
CcDSXM_AAN63597.2       ------------------------------------------------------------  0
AngDSXM_AAZ78362.1      GKRTIKTYEALVKSSLDPNSDRLTEDDEDENISVTRTNSTIRSRSSSLSRSRSCSRQAET  60
AeaDSXM_ABD96573.1      ------GYDIC-CIYFPKRSSRHRHWPDEENISVTRTPSASRSPCADFR----TRSQSSS  49
PpeDSXM                 ------AMSVC-RS---------------------------RFLCPSY------------  14

DmDSXM_NP_731197.1      ------------------------------------------------------------  0
CcDSXM_AAN63597.2       ------------------------------------------------------------  0
AngDSXM_AAZ78362.1      PRADDRALNLDTKSKPSTSSSSGTGCDRDDGDCITFDDS-ASVVRATH-ASRSATRMSRG  118
AeaDSXM_ABD96573.1      PDNNGGALNLDTKSTKATTATT------DDEE-VMYEKRSPKSIESTELRCRLEEALHSG  102
PpeDSXM                 ------------------------------------------------------------  14

DmDSXM_NP_731197.1      ------------------------------------------------------------  0
CcDSXM_AAN63597.2       ------------------------------------------------------------  0
AngDSXM_AAZ78362.1      RSRSQTKRYSQTVESTNAPSRSPGPDEEPSVYKSLAEAASKMARSFIPAREPEDLHTTTH  178
AeaDSXM_ABD96573.1      AAA---------------AAEEPLAGGSGSHWKRESF----GSTEEIPARP-------AH  136
PpeDSXM                 ------------------------------------------------------------  14

DmDSXM_NP_731197.1      ------------------------------------------------------------  0
CcDSXM_AAN63597.2       ------------------------------------------------------------  0
AngDSXM_AAZ78362.1      KSPEREDNPSQPYEAYL--------ESVRRSK-----KSFPHKDAEGVTESAEDCYDKEK  225
AeaDSXM_ABD96573.1      SEPE-----DNGFENGLEAHQSHILHSIHRNVSPKADLAVPSTSAAAVAE--------EQ  183
PpeDSXM                 ------------------------------------------------------------  14

DmDSXM_NP_731197.1      -------------ARVEINRTVAQIYYNYYTP-MALVNGAPMYLTYPSIE---QGRYGAH  43
CcDSXM_AAN63597.2       -------AKRIVNQTISLHWMDRQLYYNYYSS-AALVNTVPTYFPYPIAIGS-NGLLTSQ  51
AngDSXM_AAZ78362.1      EHRIPYSLPKSTFDRLDLLKKPNGLPFPMYKYNELEANNFPLPLLLPGLEAVNRTLYTAH  285
AeaDSXM_ABD96573.1      ------PAPPKNSEPLNLFKR---------------------SHPLPGFEQVPASALAFP  216
PpeDSXM                 ---------------------------------------------------------YYP  17

DmDSXM_NP_731197.1      FTHLPLTQICPPTPEPLALSRSPSSPSGPS-------------AV---------H---NQ  78
CcDSXM_AAN63597.2       FSHLTA-SMRPPSPEQPTLSRMPPSPSKPS-------------RP---------ASILSD  88
AngDSXM_AAZ78362.1      FPTHLLPS----SLYPPVSSESTTAPIFHTHFLGYQPQM-QLPHVEPFYRKEQQQQQLQQ  340
AeaDSXM_ABD96573.1      FPRHPAELMQFLLPYPPIFTQ----------F-GVHPPPRIYNDEASIYRKDLPSD----  261
PpeDSXM                 YLNAA-ATSHFVYNYN--F--------------ETMLSAQL--QNKSFISENVPGT----  54
                        :

DmDSXM_NP_731197.1      KPSRPGSSNGTVHSAASPTMVTTMATTSSTPTLSRRQRSRSATPTTPPPPPPAHSSSNGA  138
CcDSXM_AAN63597.2       TMSPPATAT-SLTSAATATAAT--------------------------------------  109
AngDSXM_AAZ78362.1      TLAEPKEQ-----------------TTSSSPSNNR---------LTPPKGTFFYASAVEN  374
AeaDSXM_ABD96573.1      ---------------------------SRPPSLSR---------TSPPEVPSMYHPSRVD  285
PpeDSXM                 ---------------------------RH-----------------ISESPASYTPLSLP  70

DmDSXM_NP_731197.1      YHHGHHLVSSTAAT-----     152
CcDSXM_AAN63597.2       -------------------     109
AngDSXM_AAZ78362.1      SLTAHQ--ASIATIH----     387
AeaDSXM_ABD96573.1      PSHSHQSSAPVASIH----     300
PpeDSXM                 AALQIKRLIPTSSLTTAST     89
```

**Figure S5. Multiple sequence alignment of DSXM carboxy-terminal regions.** Sequence alignment of male-specific carboxy-terminal region of DSX proteins of *D. melanogaster* (DmDSX), *Ceratitis capitata* (CcDSX), *Anopheles gambiae* (AngDSX), *Aedes aegypti* (AeaDSX) and *Phlebotomus perniciosus*. Gaps were introduced in the alignment to maximize similarity. The protein sequences alignment was performed using the Clustal-Omega software (1.2.4).

```
DmFRUM_AAB96677.1      MMATSQ--DYFGNPYALFRGPPTTLRPRESPLGVGHPHGHGHLHSHAHAHGHGHAHSHYA   58
MdFRUM_XP_019893547.1  MMTTSQ--HFFNNPYAMFHGPPPKMGPPESPHNTY-------------------------   33
AngFRUM_AAU50567.1     -MASSPALPLYASRYPTPNGYPQING-----EV-DAPLDFRKVESLR-------------   40
AeaFRUM_AGC11799.1     -MSSPPAMPNYNNRYPALNGYPQING-----VDPAEPIDCRKLQSHR-------------   41
PpeFRUM                -MMTTP--DIFNSPFQPYRGQPMAIMPPRDDSPPTPALDLKRYSSSDDPP--PPTTPVAA   55
                        * :      : . :   .* *

DmFRUM_AAB96677.1      ALDLQTPHK-RNIETDVRAPPPPLPPPPLPLPPASPRYNTDQGA          101
MdFRUM_XP_019893547.1  ALDLHTTTKPRTLDREE--RPPPYTP---PPPPTSPRFNADLGA           72
AngFRUM_AAU50567.1     ------------------------------------RNSTDTGI           48
AeaFRUM_AGC11799.1     ------------------------------------RNSTDTGT           49
PpeFRUM                HLQLPLHHRERDMVM---------PHQRPPSSQTPPRYTTDQGN           90
                                                           * .:* *
```

**Figure S6. Multiple sequence alignment of FRUM amino-terminal region.** Sequence alignment of male-specific FRU amino-terminal region of FRU proteins of *D. melanogaster* (DmFRU), *Musca domestica* (MdFRU), *Anopheles gambiae* (AngFRU), *Aedes aegypti* (AeaFRU) and *Phlebotomus perniciosus*. Gaps were introduced in the alignment to maximize similarity. The protein sequences alignment was performed using the Clustal-Omega software (1.2.4).

```
                                          BTB
DmFRU_AAB96677.1      MDQQFCLRWNNHPTNLTGVLTSLLQREALCDVTLACEG-ETVKAHQTILSACSPYFETIFLQNQHPHPIIYLKDVRYS  77
MdFRU_XP_019893549.1  MDQQFCLRWNNHPTNLTGVLTSLLQREALCDVTLACEG-ETVKAHQAILSACSPYFETIFLQNHHPHPIIYLKDVRYS  77
AngFRU_AAU50567.1     MDQQYCLRWNNHQSNLTTVLTTLLQDEKLCDVTLACEK-GMVKAHQAILSACSPYFEQIFVENKHLHPIIYLRDVEVN  77
AeaFRU_AGC11799.1     MDQQYCLRWNNHQSNLTTVLRTLLEDEKLCDVTLACDN-GIVKAHQAILSACSPYFEQIFVENKHPHPIIYLRDVEVS  77
PpeFRU                MDQQFCLRWNNHPTNLTGVLTSLLQREALCDVTLACDGGEIVKAHQTILSACSPYFESIFLQNAHPHPIIYMKDVRYS  78
                      ****:******* :*** ** :**: * ********:    *****:********* **::* * *****::**.

DmFRU_AAB96677.1      EMRSLLDFMYKGEVNVGQSSLPMFLKTAESLQVRGLTDNNNLNYRSDCDKLRDSAASSPTGRGPSNYTGGLGGAGGVA 155
MdFRU_XP_019893549.1  EMRSLLDFMYKGEVNVGQSSLPMFLKTAESLQVRGLTDNNNLNYPSELDKHRDADISSPTGRTSYGAGGGAGGPGLGM 155
AngFRU_AAU50567.1     EMRALLDFMYQGEVNVGQHNLQNFLKTAESLKVRGLTESSADRYSADTDSKLRSERIRD------------------- 136
AeaFRU_AGC11799.1     EMRALLNFMYQGEVNVGQHNLQNFLKTAESLKVRGLTESSADRYATESEKSRAERSRVD------------------- 136
PpeFRU                EMRSLLDFMYKGEVNVGQRSLPTFLKTAESLQVRGLTDNNNINYRTDSDRDRDSETNASGG------------AMKHF 144
                      .***:**:***:******* .*  ********:*****:.. .* :: :

DmFRU_AAB96677.1      DAMRESRDSLRSRCERDLRDEL-TQRSSSSMSERSSAAAAAAAAAAAVAAAGGNVNAAAVALGLTTPT--------AA 225
MdFRU_XP_019893549.1  RGERESRDRGRG----EMRDDLHSHRSSSSLSERSSATAAAVAAAVAAASGNASLQSAAATLGLTGGERSPSVGSASA 229
AngFRU_AAU50567.1     -----S---------RDERDSLPNAS---------------------------------------------------- 148
AeaFRU_AGC11799.1     -----S---------RDGRDSAPPTS---------------------------------------------------- 148
PpeFRU                DKTERDRDRDRERLERD------------------------------------------------------------- 161
                           .          :

DmFRU_AAB96677.1      AAAAVAAAVAAAANRSASADGCSDRGS---ERGTLERTDSRDDLLQLDYSNKDNNNSNSSSTGGNNNNNNNN---NNN 297
MdFRU_XP_019893549.1  AAAAVAAVVAAAAGRSASADVLNSRGDAGSDRGSDRG-----------------NDN-SVCGGVD------------ 276
AngFRU_AAU50567.1     -----------------------------------------------------SNNSNNNNNSSGNNNNNTISSNNNNNN 174
AeaFRU_AGC11799.1     -----------------------------------------------------VTNNNTI-----NSNNNNTNNNNNNNN 169
PpeFRU                ------------------------------------------------------------------------------ 161

DmFRU_AAB96677.1      SSSN-NNNSSSNRERNNSGERERERERE-------------------------------------RERDRDRELSTT 336
MdFRU_XP_019893549.1  -------RGGIDERRDDLGQI--DYSNQ-------------------------------------SKRDRDREVSTT 307
AngFRU_AAU50567.1     SLHHGPLRDKELTEHEQLERLQQQQQQQTHHQQQQHPSSHQQQSQQHPSSQHQQPSRSASIDLMQSALVDERDYLAAE 252
AeaFRU_AGC11799.1     TLHHPLQRDKELREQEELRERDRREAHR---ELQRE-----------RELQARDHQRSASAELL-TPMTDDCRYSPSD 232
PpeFRU                ----------------------REENS--------------------------------------ESKDRDRETPVD 178
                                                .                                          *
DmFRU_AAB96677.1      PVEQLSSSKRRRKNSSSNCDNSLSS----SHQDRHYPQDS-------------------------------------- 372
MdFRU_XP_019893549.1  PEHIISN--KRRRKNSSNCDNLLTSTPNANVQDRHYAQDS-------------------------------------- 347
AngFRU_AAU50567.1     DREISTVENKKKRKMSTTCDNSSPSTPSLMN-ERQGGYESQA------------------------------------ 294
AeaFRU_AGC11799.1     DRDLT-VESKKK-RKISTCDNSLPSTPSLMN-DRPGGYESQVSFVGSLFSRQHIYHFFMLICIILNTKLQDSSQRSNK 307
PpeFRU                H-LSSSSSTKRRRKNSLNCDNSMHG---ASVQDRHYSQDS-------------------------------------- 214
                               ::: .   .***   .       :*    :*

DmFRU_AAB96677.1      -------------Q----ANFKSSPVPKT------GGSTSESEDAGGRHDSPLSMTTSVHLGGGGGNVGAASALSGLS 427
MdFRU_XP_019893549.1  -------------QAP--SNFKSSPVPKSSTGGGGGGNTSETEDSGGRRDSPLSASA----LSGGGNVNASSGGMGLN 406
AngFRU_AAU50567.1     ---------------SSHSSFKQSPKPEDEF--------------------KVSSPAPVHPLAG-------------H 324
AeaFRU_AGC11799.1     HFQFNLTVPLILQQASSHSSIKLSPKPDDDF--------------------KVSSPAAMHSLSG-------------P 352
PpeFRU                -------------QASSHSSYKSSPLPKLNPLEGED----------TRRNSP------------------SLNASGAN 251
                                      :. * ** *.

DmFRU_AAB96677.1      QSLSIKQELMDAQQQQQHREHHVALPPDYLPSAALKLHAEDMSTLLTQHALQAADARDEHNDAKQLQLDQTDNIDGSS 505
MdFRU_XP_019893549.1  QSLSIKQELMDAQQQ-QQREHHVSLPPEYLPPGALK-HSEDMASLLSSHSMQAADSREDHNDAKQLPFDQSDNIDAPR 482
AngFRU_AAU50567.1     PLSGIKQELPEL------PVR-HSLSSELMQPIPINMNADDMNNILAPGNMGSLNESG-DSDAHLSHPDHPDNIDGPG 395
AeaFRU_AGC11799.1     ----IKQEYSDL------PSRHHPLHPELLPPIPMNLNPEEMSNMLSQGNMPPLNDSG-DNEGSLSHPDHPDNIDGPG 419
PpeFRU                QSVSIKQELPD-------MGHHPGLPPELLPPTSMSLHPEDMTSLLPAHGLQVESSE---NDSQHPQMDHSDNIDGPG 319
                          ****  :          :   *  : :    :. : ::* .:*    :   .     .:.     *: ****.

DmFRU_AAB96677.1      ARHHLST----PLS--TSSSASPPP---PPFGMHLSAALKREYHPLHYMAAGNGHNGPSALGYGNQGSGNAPNSAGGA 574
MdFRU_XP_019893549.1  GRMSRTSFRRSSFSRASSLSDSPPPHPAATYGLQFDAAIKNEYLLENIMRGANGGHHQNDQ-----A----------H 545
AngFRU_AAU50567.1     GGGGGSRYEHHLSRHASSIL------------------------PSSLVSS-----------------PDG------- 425
AeaFRU_AGC11799.1     GG----QYEHHLSRHASSIL------------------------TSCLVTS-----------------PDG------- 445
PpeFRU                GHR-----------------PPPPLPPPPHLHHLGQHCKSEHPQ---VPGGSAR------------CGD-------S 357
                      .                                                    ZnF-C
DmFRU_AAB96677.1      GSVAGGVGAGGGAGGATGAAGHNSHHTMSYHNMFTPSRD-PGTMWRCRSCGKEVTNRWHHFHSHTAQRSMCPYCPATY 650
MdFRU_XP_019893549.1  HQSLNQLHTLGSATGNMGSSGHNMHHQMSYHNMFTPSRD-PGTMWRCRSCGKEVTNRWHHFHSHTAQRSNCPYCPATY 620
AngFRU_AAU50567.1     -------------TDLPHHTHYQLHHQMSYHNMFTPSRE-PGTAWRCRSCGKEVTNRWHHFHSHTPQRSLCPYCPASY 488
AeaFRU_AGC11799.1     -------PSGGGGGDLSLHPHFSPHHQMSYHNMFTPSRE-PGTAWRCRSCGKEVTNRWHHFHSHTPQRSVCPYCPASY 516
PpeFRU                GSSRGS-PTVAVVA-AALQHHQHHQQQMSYHNMFSPTQELAGTMWRCRTCGKEVTNRWHHFHSHTAQRSMCPYCPATY 433
                                :  .          :: *******:*:::  ** ****:*************** *** ******:*

DmFRU_AAB96677.1      SRIDTLRSHLRVKHPDRLLKLNSSI----------- 675
MdFRU_XP_019893549.1  SRIDTLRSHLRVKHSDRLLKLGSSM----------- 645
AngFRU_AAU50567.1     SRIDTLRSHLRIKHADRLNA---PKFSNPPNCKLPM 521
AeaFRU_AGC11799.1     SRIDTLRSHLRSKHADRLTAPPTPKFGTPPNCKMQM 552
PpeFRU                SRIDTLRSHLRVKHPDRLIKN--------------- 454
                      *********** ** ***
```

**Figure S7. Multiple sequence alignment of FRU proteins.** Sequence alignment of FRU proteins of *D. melanogaster* (DmFRU), *Musca domestica* (MdFRU), *Anopheles gambiae* (AngFRU), *Aedes aegypti* (AeaFRU) and *Phlebotomus perniciosus*. The conserved BTB domain and Zinc Finger type-C domain are boxed in grey. Gaps were introduced in the alignment to maximize similarity. The protein sequences alignment was performed using the Clustal-Omega software (1.2.4).

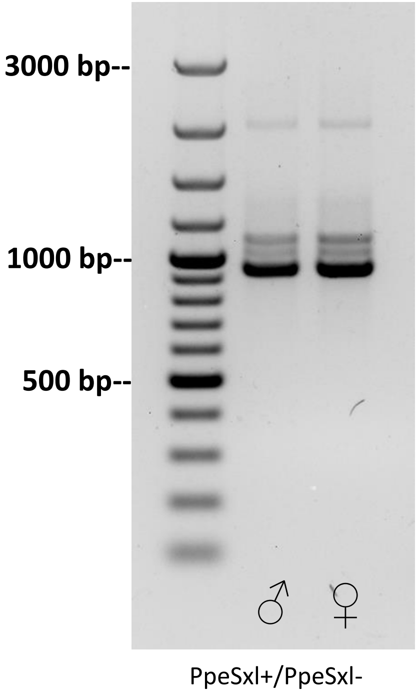


**Figure S8. *Sex-lethal* gene expression at adult stage in *P. perniciosus*.** RT-PCR analysis on total RNA extracted from adult males and females of *P. perniciosus* with primer pair PpeSxl+/PpeSxl-. GeneRuler 100 bp Plus DNA Ladder (ThermoFisher Scientific) was utilized as molecular weight marker.

```
                                                                                                    CAM domain
DmTRAF_AAF49441.1      ------------------------------------------------------------------------------------------------------------------------      0
AoTRAF_ABW04165.1      --MSIPK---------------ASTATRKIQIEQSVPTGSIRKGPYAIERSVDA-SEVVIKRRFGEGSKPLFQRDDIVVNPNSVIKATDSHTEKHLISESKNRAKDVSNQYLKGSRSSTD      102
AgTRAF_ABW04170.1      --MNIPK---------------ASTATRKIQIEQSIPTGSIRKGPYAIERSVDA-NEVVIKRRFGEGSKPLFQRDDIVVNPNSVINVADSHTEKHLISESKKRAKGVSNEYLKRSRSSTD      102
CcTRAF_AAM88673.1      MNMNITK---------------ASATTRKIRIEQNVPSGSVRKGPYAIERSVNP-SEVVIKRRFGEGSKPLFQRDDIVVNPDNVVSNVGAHFETQPKDRSNNSKEEVENQWRKERHKSTD      104
BoTRAF_CAG29243.1      MNSNIPK---------------LFATSSKIQIKQHVPNGSIRKGPHAIERSLVP-DEVVIKRRFGEGSKPLFQRDDIVVNPESVTNTADAQNDKQIQAETRNNTEDISNRRRKERHISTD      104
BtTRAF_AIK25402.1      MNSNTPK---------------LFATSSKIQIKQHVPIGSIRKGPHAIERSLVP-DEVVIKRRFGEGSKPLFQRDDIVVNPDNAINIAEDKSE------KQSTTEDISNRGRKERHISTD      98
MdTRAF_ACY40709.1      MEQTSMGKDSSKTAIAKFVPDDDGVLKHTIKVHQQPTSATEKKGPSMIARSNDLIEEIQIKRRFGEGTKPLFERDDVKVNTVADGESISS--------------KKI---KSH------K      97
ChTRAF_AGE31793.1      MDSNITG-----------LSASSTLGGPKFKIHQSIPSGSVKRGPHAIVRSVELNDCIEIQRRCGEGSIPLFDRHDITVNQTINESNDK-----QINHEHHTSG-KT---KTNRRSSSRE      100
LcTRAF_ACS34689.1      MDSITTG-----------LAASSILEGTKFKIQQSIPSGSIKRGPHAIVRTADLNDGINIQRRFGEGSKPLFDRDDIAVNQTTLDARDTN----NLDHERNTGCKTN---KTNRRSSSTE      102
LsTRAF_AGE31795.1      MDSIATG-----------LAASSTLEGTKFKIQQSIPSGSIKRGPHAIVRTADLNDGINIQRRFGEGSKPLFDRDDIAVNQTTLDARDTN----NLDHERNTGCKTK----TNRRSSSTV      101
PpeTRAF                MS-TI-------------RK--SSSTAHKERYETKHNNG----GKMATKREAER-RPSTRQDSFK-----KSHREESHIPSTSRTPRDGSQHK-------RTGDRRESSKHRK-DASRRD      86

                                                                                                                      Diptera domain
DmTRAF_AAF49441.1      -------------MKMDA-----------DSSGTQHR-----DS--RGS-RSRSRREREYHG-------RS-SE---RDSRKKEHKIPYFADEVREQDRLRRLRQRAHQSTRRTRSRSRS      77
AoTRAF_ABW04165.1      SSSPEGYR--KYHTGPYNDCTTSTNNRSPPRTKPL------------------KSTNEG--KHTIRS-DSSPPF---NHRRRTPEKLPYFIDETRERDRLRRKYETKPKIPSSSPPLSSR      196
AgTRAF_ABW04170.1      SSSPEGYR--KYS-SRYNDCITSTNNSAAPRTKPL------------------KSTNER--KRTIRC-DSPPPF---NHRRRTPEKLPYFIDETRERDRLRRKYETKPKIPSTSPPLTSR      195
CcTRAF_AAM88673.1      SSSPERFR--KHHSSNKSEHSNSGNNITTRHTKTH------------------HPSQENLNTASKRR-DSSPPT---NRRHRTPEKVPYFIDEIRERDRIRRKYGKRSTKSPSPPVMSSK      200
BoTRAF_CAG29243.1      SSSPERYR--KYHTSQKNESEIGSSNNTTRRTKTA------------------KPTSDG--KYAVRR-DVSPPP---NRHRRIPEKIPYFVDEVREQDRIRRKYGSTKNKS---PPASSK      195
BtTRAF_AIK25402.1      SSSPERYR--KYHNNLEK--------ETTRRTKTT------------------NPISDD--KYAARR-DASPPP---NRRRKTSEKIPYFVDQVRERDRIRRKYGSTRNKS---PLASSK      181
MdTRAF_ACY40709.1      ASSPSKGV--N-GLVKQNSPDVTQ--KFTKKYGSSENPDFR----------RHSSYEKDNYHKSN----SKS-GV---HLEGHEGNYRPYFSDPIRERDRLRRLYGSPNERRSRSHSPSSN      195
ChTRAF_AGE31793.1      SSSPERYR--HRDIKKHSSTSMSRRRKTPERTVRSERPSNNRDHHSYNLKHGSNTKTDKYSRKDKSK-TRTPSP----NKKKPGEKVPYYRDEQREKDRLKRLYGRSCSRSPPPTSRS-S      212
LcTRAF_ACS34689.1      SSSPERYR--HRDIKKHSSPT-SGRRKTPERSGRSERPTHSHDKHNYNVKSN-NTMTDKYKRSRRSK-SRSPPH---NANKKTVEKVPYYRDEQREKDRIRRLYGRSRSRTPPPPVGNLS      214
LsTRAF_AGE31795.1      SSSPERYR--HRDIKKHSSPT-SGRRKTPERSGRNERPTHSHDKHNYNVKSN-NTMTDKYKRSRRSK-SRSPPH---NANKKTVEKVPYYRDEQREKDRIRRLYGRSRSRTPPAAAT---      210
PpeTRAF                SSSSSDSEDSDKELGKSS-----RRRENSRR-------------------RRSKSRSRSLDDSRRRMSRRSPPRRRPKSETPPRESKPYYADAQRERDRLREKYGRDEERHRS-------      175
                                                                                                            .  **: *  **:**::.

DmTRAF_AAF49441.1      --------QSS----IRESR-------------HRRHRQ-----------------------------------------------RSRSRNRNRSRSSERKRRQHSRSRSSERRRRQ--      123
AoTRAF_ABW04165.1      PRHRRSRSKSRSRSQSRELEHR-----LKYANL-RRSVSTDRYIGETGRRERGSRSD--------KTSSRYRHRSEDRSKSDRRQRHWRSPSRSPRRSRDRLTRSRKQSRHDERHKNYHS      302
AgTRAF_ABW04170.1      SRHRRSRSKPRSRSHSREFEHR-----LKPANF-RRSVSTDRYIGETSRRERGSRSE--------KTSSRYRHRSEDRSKNDRRQRHWRSPSRSPSRSRDRLTRSHKQSRHDERHKNCHS      301
CcTRAF_AAM88673.1      FRRRRSYSRSISRSRSHSPARSKNRTHV-YGSLSRRSSSVDRYIGGGRKRRRENLR--TERDRDGQYRHHGH-RSE-EQERSRRGRSPRARTRSRTRSRERSK--HVRARNDERNKNLHG      313
BoTRAF_CAG29243.1      FRRRRSISKSCSRSHSRDSMKTKQRSQTRRTYF-RRSISVDRYMGNNSKRERETEKSRTDKDLGGTPSHHYQHTSKDRAKNLRRHRSSRTHSRSRTRSRERSS--RIGTQSSERHRYRHN      312
BtTRAF_AIK25402.1      FRRRRSISRSRSRSHSRDSLKTKQRSPARRTNFRRRSISMDREWGDNSKREREREKSRADKDLHGTPR-HYQHRSDDRGKNVRRSRSSRTHSRSRTRSRDRSS--RIGTQNSERHKYRYN      298
MdTRAF_ACY40709.1      RRRSSHRSRRG----SGSPR-------------SRRY--TSRHRR-------------------RSNS---Q---D--RTSWKHNPEHRTSRRSRTRSPRGNRSRRRSSTSSNEDDERE-      268
ChTRAF_AGE31793.1      GRKYSPITKRR----IESPR-------------RRRSRSKDRY---------------------RKYS---PHRSS--RKDYRTHRSSRSRSRSRTRSPSRRERHKHYSRSSRERDKEH-      288
LcTRAF_ACS34689.1      SSSSSSTAKRR----IESPS-------------RRRRRSTSRDRH-------------------RRHS-PYIHRSV--RRDYR---SRRSRTRSRTRSPQRRERHKHSSRSSRERDKEH-      291
LsTRAF_AGE31795.1      ---LSSSTKRR----IESPT-------------RRRRRSTSRERR--------------------RHS-P-YHRNV--RRDYR---SRRSRTRSRTRSPPRRERYKHSSRSSRERDNEH-      282
PpeTRAF                ----------R-----------------------RRSYS--RE-----------------------------------------------------------RRR-----------------      186
                                                         **

DmTRAF_AAF49441.1      --------RSPHRYNPPPKIINYYVQVPPQD-------FYGMSGMQQSFGYQR--LPRPPPFPPAPYRYRQRPPFIGVPR-----FGYRNAGRPP-------------------Y-----      197
AoTRAF_ABW04165.1      EHGGASSDELAQRNLPQPQIITIPVPVPADF--M----NYTYPTWPT--QWNPPMAHPVRYGPPAAYHIPTILPAAVMPPMRPPLPPY---GPHPPPLRYGVRGF----RLPPQYGASRP      407
AgTRAF_ABW04170.1      EHKGSSNDELAQRNLPPPQIITIPVPVPADF--M----NYTYPTWPT--QWNPPMAHPVRYGPPAAYHIPTILPAAVMPPMRPPLPPY---GPHPPPLRYGVRGF----RLPPQYNASRP      406
CcTRAF_AAM88673.1      NHDELTNAELNQRNLTQPQIITIPVPVPADF--L----NYAYSTWPTQTQWSHPMTPPPRYGAP-AYHMPTILPATVMPPMRPALPPY---GLPPQPMRYGGRGL----RFPQQHG-PRP      418
BoTRAF_CAG29243.1      DNDEK--NGNDERNMPQPQIITIPVPVPADF--M----NYGYPTWPTPTQWSP---QPSRYGAP-PYPMPTFLPA-VLPPLRHPMPPY---GLPPQPIRYGGPGY----RLPSQYGASRP      412
BtTRAF_AIK25402.1      ENEEQ--NGNGERNLSQPQIITIPVPVPADF--I----NYGYPTWPTTTQWAP---QTSRYGTP-TYPMPTFVPA-VLPPLRHPMPPY---GLPPQPIRYGGPGY----RLPVQYGATRP      398
MdTRAF_ACY40709.1      --YRHRHHR--SQERSYPNVLPTLPPALTNYPC----------HYHVAPM-LA--LPGV----------QHRPFLPMVASVRHLPPQALYGGLAGAMPFIPMPMTAYRPHLGHRYPPPRH      361
ChTRAF_AGE31793.1      --KEDVNNRSTAILPPTPQFI-LPVAVPADYAAA----AYTFPGWTTAPQ-LA--WHPGHHRPPAA---APFPMVPMLPALRPPTHQASYAGLPP-ALAYPPITSPYRPHIPQRYPPPRH      394
LcTRAF_ACS34689.1      --KEDVNSLTTAIIPATPQIIPIPVPVPAEYA-A----AYTFPGWT-APQ-PT--WPPSHHRPPATSHFAPFPMWPMMPPLRPPPHQASYGGLPPQH----------------------      377
LsTRAF_AGE31795.1      --KEDANSLTTAIIPATPQIIPIPVPVPAEYA-A----AYTFPGWT-APQ-PT--WPPSHPRPPSASHFAPFPMWTMMPPLRPPPHQASYGGLPPPALAYPPMAASYRPHAPQRYPQPRH      391
PpeTRAF                -------RRSRTPVRETTKIVTVPVPVYPTFYPDGSVYEWYDPAWPGGP---RPMLSQPPMRPPGFYPGGHFMVDPFHRPMRP--------GMAPPRPFRPPMGF----------PQPRF      278
                                        :.:     .                                                                 *

DmTRAF_AAF49441.1      --------------        197
AoTRAF_ABW04165.1      WRPNFRTKNT----        417
AgTRAF_ABW04170.1      WRPNFRTKNT----        416
CcTRAF_AAM88673.1      WRPNFRPKTHK---        429
BoTRAF_CAG29243.1      WRPNFRSKNL----        422
BtTRAF_AIK25402.1      WRPNFRSKNT----        408
MdTRAF_ACY40709.1      KINKKN--------        367
ChTRAF_AGE31793.1      NTNNYHSRPKKPTS        408
LcTRAF_ACS34689.1      --------------        377
LsTRAF_AGE31795.1      DSNNYQTRPKKPSS        405
PpeTRAF                QRPH----------        282
```

**Figure S9. Multiple sequence alignment of TRA proteins.** Sequence alignment of TRA proteins of *D. melanogaster* (DmTRAF), *Anastrepha obliqua* (AoTRAF), *Anastrepha grandis* (AgTRAF), *Ceratitis capitata* (CcTRAF), *Bactrocera oleae* (BoTRAF), *Bactrocera tryoni* (BtTRAF), *Musca domestica* (MdTRAF), *Cochliomyia hominivorax* (ChTRAF), *Lucilia cuprina* (LcTRAF), *Lucilia sericata* (LsTRAF) and *Phlebotomus perniciosus* (PpeTRAF). The putative autoregulation CAM (*Ceratitis-Apis-Musca*) domain is highlighted in yellow while the Diptera domain is highlighted in green. Gaps were introduced in the alignment to maximize similarity. The protein sequences alignment was performed using the Clustal-Omega software (1.2.4).
