## Additional_file_3_S10_S13 for "Genomics and transcriptomics to unravel sex determination pathway and its evolution in sand flies"

### *P. perniciosus tra* gene model

**Clone_G1_Length=1725**

120 EXON1 — 193 — 0.64 gcggaagGTaaggtc — 0.85 attt

DNA: gatccaattgtgcaattgtcgcgaaaagaaaacaccgcaagaaaattccaattttccagtgaatacattaaaaatcagtgaatttctactcacaggtgactcccaagtgtatccagtgtATGTCTACGATACGGAAATCCAGTTCAACAGCGCACAAAGAGCGATATGAGACGAAACACAACAATGGCGGAAAGTAAGGTCGCCATGAGAAGGAATGATTTATGGCTTTTGCGATGCATTT
+3fr: ·M··S··T··I··R··K··S··S··S··T··A··H··K··E··R··Y··E··T··K··H··N··N··G··G··K

259 EXON2 — aatgggttattttgcAGaatggcgacaaagagggagg — 392 EXON MS1 — 0.42 cggacggGTaatgaa

DNA: AATGGGTTATTTTGCAGAATGGCGACAAAGAGGGAGGCGGAGAGACGTCCATCAACGCGTCAAGACAGCTTCAAAAAGTCCCACAGGGAGGAAAGTCATATTCCCTCCACTTCCCGGACACCCCGGGACGGCTCTCAACACAAACGGACGGGTAATGAATCTCTCAACAGGCTGAGGATTTGATGAgtatttgagatatcaaagtcaggatagaattcatgagaatttattgagttacta
+2fr: ··M··A··T··K··R··E··A··E··R··R··P··S··T··R··Q··D··S··F··K··K··S··H··R··E··E··S··H··I··P··S··T··S··R··T··P··R··D··G··S··Q··H··K··R··T··G··N··E··S··L··N··R··L··R··I··*··*·

624 — 0.89 gattgatGTaagtgt — 0.94 ggtaattaaatctacttgcAGgtgttcaggagctgtggaag — 669 EXON FS1

DNA: agatcaacaagcaacactgaagacatccctcgcagccggaaaccgagggattagcaattTCATCAATCAACActcgatcaacaagtagaaaagccaaaggcggaagcagctgggagttccgggatggatgcaaatgattgatGTaagtgtgaatactttttttactggtaattaaatctacttgcAGGTGTTCAGGAGCTGTGGAAGAGCAAAACAGATCGGAAATTCTTCAATCAACACAA
G··V··Q··E··L··W··K··S··K··T··D··R··K··F··F··N··Q··H··N

DNA: TCAACATTGATAAAAACATCCAATTGGCGGAAATAGGGATTCAAAATTAAATTTCTTGCTCAATTAATCTTCAATCAACAcgatataagcaacaagcaacaTCTTCAATCAACAcaaaccctaaggatcTCTTCAATCAACAcgtagtttagattttaaatcaaatttttaaatattggaatcaattgtagatttttgagagaaattatcagaatatttccttaaaaaatgttttcaagg
··Q··H··*··*·

1012 EXON3 — 0.91 tgtactaaaatccccccacAGgtgatcgcagggaatcctcc — 1067 — 0.58 ttcatcaGTgagtag — 1126 — 0.92 tctaatcagaaattcttgcAGagctcctcggactcagagga

DNA: actttttgagagtcaaaatgtaaatttgtactaaaatccccccacAGGTGATCGCAGGGAATCCTCCAAGCATCGCAAAGATGCGTCAAGGAGAGATTCATCAGTGAGTAGATTCCTATTGTAATTCTCTAGAAAAAAAAATCTAATCAGAAATTCTTGCAGAGCTCCTCGGACTCAGAGGATTCAGATAAGGAGCTTGGGAAGTCATCACGGAGACGTGAGAACTCTCGCCGGAGACGCTC
+3fr: ·G··D··R··R··E··S··S··K··H··R··K··D··A··S··R··R··D··S··S ·S··S··S··D··S··E··D··S··D··K··E··L··G··K··S··S··R··R··R··E··N··S··R··R··R··R··S

DNA: AAAGAGTCGTAGTCGGAGTCTCGATGACTCGCGGCGTCGCATGAGTCGTCGATCTCCCCCACGACGTCGACCAAAGTCAGAGACGCCTCCGCGAGAGTCCAAGCCCTACTATGCTGATGCTCAGCGCGAGAGGGATCGCTTGAGGGAGAAATATGGACGTGATGAGGAGCGCCATAGGAGTCGCAGGCGAAGCTACAGTCGCGAGAGACGCCGGCGACGCTCGAGGACGCCCGTGAGGGAG
+1fr: ··K··S··R··S··R··S··L··D··D··S··R··R··R··M··S··R··R··S··P··P··R··R··R··P··K··S··E··T··P··P··R··E··S··K··P··Y··Y··A··D··A··Q··R··E··R··D··R··L··R··E··K··Y··G··R··D··E··E··R··H··R··S··R··R··R··S··Y··S··R··E··R··R··R··R··R··S··R··T··P··V··R··E·

DNA: ACTACAAAGATCGTGACGGTTCCCGTGCCAGTCTATCCCACCTTCTATCCCGATGGCTCAGTGTATGAGTGGTATGATCCCGCGTGGCCAGGGGGCCCGCGTCCCATGCTGTCTCAGCCACCAATGCGACCACCAGGATTCTATCCCGGCGGTCATTTCATGGTGGATCCCTTTCATAGGCCCATGAGACCCGGAATGGCACCTCCAAGGCCATTCCGGCCGCCTATGGGCTTCCCCCAGC
+1fr: ·T··T··K··I··V··T··V··P··V··P··V··Y··P··T··F··Y··P··D··G··S··V··Y··E··W··Y··D··P··A··W··P··G··G··P··R··P··M··L··S··Q··P··P··M··R··P··P··G··F··Y··P··G··G··H··F··M··V··D··P··F··H··R··P··M··R··P··G··M··A··P··P··R··P··F··R··P··P··M··G··F··P··Q··

1707

DNA: CGCGATTTCAAAGACCCCATTGAAAATTGACTGAATAG
+1fr: P··R··F··Q··R··P··H··*·

**Figure S10. Manually-curated *P. perniciosus transformer* gene model.** Exonic sequences are indicated by black cases; black upper cases indicate coding sequences. Intronic sequences are indicated by gray lower cases. Start and stop codons are highlighted in green and red, respectively. In yellow boxes, putative TRA/TRA-2 binding sites. Exon start/end positions in the scaffold/s are indicated. Intron boundaries were predicted using transcripts vs genome alignments and confirmed by *de novo* prediction with Berkeley BDGP Splice Site Prediction Tool with default parameters ([http://www.fruitfly.org/seq_tools/splice.html](http://www.fruitfly.org/seq_tools/splice.html)); prediction scores are indicated in red. Azure box indicates the male-specific exon 1 sequence; Purple box indicates the famel-specific exon 1 sequence.

### *P. papatasi tra* gene model

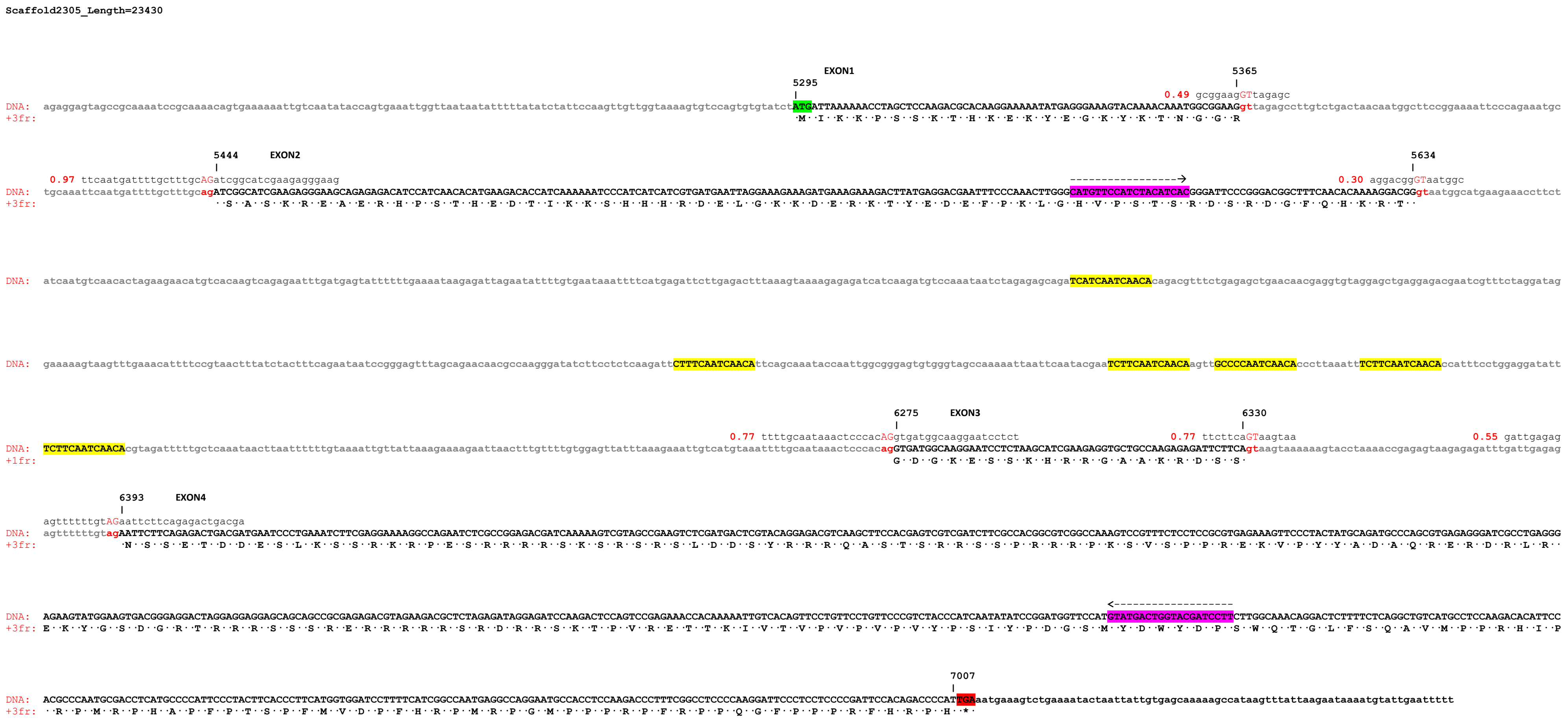


**Figure S11. Manually-curated *P. papatasi transformer* gene model.** Exonic sequences are indicated by black cases; black upper cases indicate coding sequences. Intronic sequences are indicated by gray lower cases. Start and stop codons are highlighted in green and red, respectively. In yellow boxes, putative TRA/TRA-2 binding sites. Exon start/end positions in the scaffold/s are indicated. Intron boundaries were predicted using transcripts vs genome alignments and confirmed by *de novo* prediction with Berkeley BDGP Splice Site Prediction Tool with default parameters (http://www.fruitfly.org/seq_tools/splice.html); prediction scores are indicated in red. In purple boxes are indicated the positions of the primers utilized for *PpatraF* amplification.

### *P. bergeroti tra* gene model

```
Contig_32222_414377_7522_Length=2705

DNA:  caaatatctggcggctttcccgaaatgaaacgtgggaggtttttttttcactttaatatcacgagactctttgtaacacacctcaataaacacctgaaacagacacaagatgtgacttcagacactgggtcaaggtttaacgaatatttaaaggcaatttttattgaattcagtgtgccagatgactttccagataagaaaatttagagatttctcaatgaattcccttccggaaaccc

DNA:  atccgccattgctttcttcccgtcgttgataaaccacgccacaatgtccactcaccgacttgcagcaaaatgtcccccgacgattatttatcacactcacaattataaatagagtgaaaggaggaaaaaggtcagagatttgagggaaaatataaatgcaaatgtagcgagaaatgtgacaagtctgcggccatgaagagactccactttttccacctcaaatcacactctgaataccac

DNA:  aaatttttcacttatttcaaatgcagaagatacttggacactttaatttgaggttcattgtatttttttccactaatttttagtatttttaggcgaaaattgcatttattccgcagttgcatattccgatttctgcaccttttacttgtcaaaccagctgtttttttagccaccaggcggcgctaattgccatattgtgaaataaaaaaaaggcctattagcaagaggagtagccgcaaaa

                                                                                                         EXON1
                                                                                                     827                                                              897
                                                                                                     |                                                                 |
                                                                                                                                                        0.82 gcggaagGTtagtgt                                                       0.97 tcaatgatt
DNA:  tccgcaaaactgtgaaaaaattgacaaattaccagtgaaagttacttaaatattatttttatatctattccaagttgttggtaaaagtatccagtgtgtatctATGATTAAAAAACCTAGCTCCAGGACGCACAAGGAAAAATATGAGGGAAAATACAAAACAAATGGCGGAAGgttagtgtatagacagcttaacaatggcttccggaaaattcccagaaatgctgcaaattcaatgatt
+3fr:                                                                                                        ·M··I··K··K··P··S··S··R··T··H··K··E··K··Y··E··G··K··Y··K··T··N··G··G··R

          977     EXON2                                                                                                                                                                    1167
          |                                                                                                                                                                                 |
      tttgcagatcAGcatcgaagagggagg                                                                                                                                                0.41 agaacggGTaatggc
DNA:  tttgctttgcagATCAGCATCGAAGAGGGAGGCAGAGAGACATCCATCAACACATGAAGACACCATCAAAAAATCCCATCATCATCGTGATGAATTAGGAAAGAAAGATGAAAGGAAGTCTTATGAGGATGATTTTCCCAAACTTGGGCATGTTCCATCTACATCACGGGATACCCGGGACGGCTTTCAACACAAAAGAACGGgtaatggcatgaagaaatcttctatcaatgtcaacact
+1fr:             ··S··A··S··K··R··E··A··E··R··H··P··S··T··H··E··D··T··I··K··K··S··H··H··H··R··D··E··L··G··K··K··D··E··R··K··S··Y··E··D··D··F··P··K··L··G··H··V··P··S··T··S··R··D··T··R··D··G··F··Q··H··K··R··T··

DNA:  agaagaacatgtcacaagtcagagaattttgatgagtattttattgaaataaagagattagaatattttgtgaaataattttcatgagattcttgagattttaaagtaaaagagagatcatcaagatgcccaaataatctagagagcagaTCATCAATCAACAcagacgtttctgagagctgaacaacgatgtgtaggagctgaggagaagaatcgtttctaggattgaaaaaagtaagtt

DNA:  tgaaatattttccgtaactttatctactttcagaataatccaggagcttagcagaacgccaagggatatttttctctcaagattCTTTCAATCAACAttcagcaaattccaattggcgggagagtgggtagccaaaaagtaattcaatacgaaTCTTCAATCAACAagttGCCCCAATCAACAcacttaaattTCTTCAATCAACAcaatttcctgaaggatattTCTTCAATCAACAcgt

                                                                                                   1807     EXON3                                      1862                                                       1926
                                                                                                   |                                                   |                                                          |
                                                                               0.85 ttgcaataaactcccccacAGgtgatggcagggaatcctct                 0.77 ttcttcaGTaagtaa                                  0.69 gatttagggagtttttttgtAGaat
DNA:  agattttgctcaattaacttttttttgaaaaattttattaaagaaaagaattaactttgttttctggagttatttaaagaaattgtcatgtaaattttgcaataaactcccccacagGTGATGGCAGGGAATCCTCTAAGCATCGAAGAGGTGCTGCCAAGAGAGATTCTTCAgtaagtaaaaagaaaagtacttaaaatcgggtgtagagatttgatttagggagtttttttgtagAAT
+2fr:                                                                                                  G··D··G··R··E··S··S··K··H··R··R··G··A··A··K··R··D··S··S·                                                ·N·

      tcttcagagactgacag
DNA:  TCTTCAGAGACTGACAGTGAATCCCTCAAATCTTCGAGGAAAAGGCCAGAATCTCGCCGGAGACGATCAAAAAGTCGTAGCAGAAGTCTCGATGACTCGTACAGGAGACGTCAGACCTCCACGAGTCGTCGATCTTCACCACGACGCCGGCCAAAGTCTGTTTCTCCACCGCGGGAGAAAGTGCCCTACTATGCAGATGCCCAGCGTGAGAGGGATCGCCTGAGAGAGAAGTATGGAAGTG
+3fr: ·S··S··E··T··D··S··E··S··L··K··S··S··R··K··R··P··E··S··R··R··R··R··S··K··S··R··S··R··S··L··D··D··S··Y··R··R··R··Q··T··S··T··S··R··R··S··S··P··R··R··R··P··K··S··V··S··P··P··R··E··K··V··P··Y··Y··A··D··A··Q··R··E··R··D··R··L··R··E··K··Y··G··S··

DNA:  ATGGGAGGATGAGGAGGAGGAGCAGCAGCCGCGAGAGACGCAGAAGACGCTCTAGAGATAGAAGATCCAAGACTCCAGTCCGAGAAACCACAAAAATTGTCACAGTTCCTGTTCCTGTTCCCGTCTACCCATCAATTTATCCAGATGGTTCCATGTATGACTGGTACGATCCCTCTTGGCAAACAGGGCTCTTTCCACAGGGTGTAATGCCTCCAAGACACATTCCACGCCCAATGCGACC
+3fr: D··G··R··M··R··R··R··S··S··S··R··E··R··R··R··R··R··S··R··D··R··R··S··K··T··P··V··R··E··T··T··K··I··V··T··V··P··V··P··V··P··V··Y··P··S··I··Y··P··D··G··S··M··Y··D··W··Y··D··P··S··W··Q··T··G··L··F··P··Q··G··V··M··P··P··R··H··I··P··R··P··M··R··P

                                                                                                                                          2540
                                                                                                                                          |
DNA:  TCATGCCCCATTCCCTACTTCACCCTTCATGGTGGATCCTTTTCATCGGCCAATGAGGCCAGGAATGCCACCTCCAAGACCCTTTCGACCTCCCCAAGGATTCCCTCCTCCCCGATTCCACAGACCCCATTGAaatacaaagtctgaaaatactattgtgagcaaaagccattttaagaataaaatgtattgaatttttattgtttttttttttagagaggaattctcgttatagagaaat
+3fr: ··H··A··P··F··P··T··S··P··F··M··V··D··P··F··H··R··P··M··R··P··G··M··P··P··P··R··P··F··R··P··P··Q··G··F··P··P··P··R··F··H··R··P··H··*·

DNA:  tattaataggggaaagtggggcatatttgaaagtcgagcacctttgaaattggg
```

**Figure S12. Manually-curated *P. bergeroti transformer* gene model.** Exonic sequences are indicated by black cases; black upper cases indicate coding sequences. Intronic sequences are indicated by gray lower cases. Start and stop codons are highlighted in green and red, respectively. In yellow boxes, putative TRA/TRA-2 binding sites. Exon start/end positions in the scaffolds are indicated. Intron boundaries were predicted using transcripts vs genome alignments and confirmed by *de novo* prediction with Berkeley BDGP Splice Site Prediction Tool with default parameters (http://www.fruitfly.org/seq_tools/splice.html); prediction scores are indicated in red.

### *P. duboscqi tra* gene model

```
Contig_184858_395575_582582_842104_Length=3119

DNA:  tttcatttttccgaattcgaacatttagccaattttcttgccattggacttaaaattaaattttgactttaaaatttaaatgcctctagtcagaatttgttttcattgatagattcacagtgtgttcatagaaatcagacaccaatcctagctacgccactgaatggggatggagaaagtgttcataaacttaccaggaagacgggatcgagaataccatcgtctactaggggctcagca

DNA:  aacaaacgatcacagtccatatctggaaattttcatgagggaaaacagagagagaaaacgttttttcatgagcttactgggcagtttacaagattgtgcacgagacattaacctggacgttctgatgtctcaggtgtgtccacggttgtcgattgaaacattgcggcaacattgagaaaattccggagggaaacactcagtgaatcatgatcgcgctgacgatctgatccatctgtatttt

DNA:  ccatgtttcagcacaaaatggggcacagcagaatctctgccctcatttgatttatacacatacactgacctgcaaagcctctctcttcccgcacaaatatctggcggctttcccgaaatgaaacgtgggaggtttttcacttatatatcacgagactcttctaacacacctcaataaacacctgaaacagacacaagatttgacttcagacactgggtcaaggtttaaagaatctttagg

DNA:  ggtaatttccattgaattcaattagcttagatgatttccagataagaaaattcagagatttctcaaacgaattccgttccggaaacttatccgccattgctttcttccctgttgtagataaaccacgccacaagtccactcaccgatttgcagcaaaatgtcccccgacgattgtttatcacactcacaattataaatagagtgaaaggaggaaaaaggtcagagatttgagggaaaataa

DNA:  atgcaaatgtagcaagaaatgtgacaagtctgcggccatgagagactcactttttcccacttcaactcacactctgaataccacaaatttttcacttatttcaaatgcagaagatacttggacactttaacttgaggttcattgtgtttctttccactaattttgagtatttttaggtgaaaaattgcatttattccgcacttgtatattcgatttctgcaccttttacttgtcaaaccag

                                                                                                                                                                                                    1387     EXON1
                                                                                                                                                                                                    |
DNA:  ctgtttttttagccaccaggcggcgcttattgccattgtgaaataaaaaaaggcctattgacaagaagagtacagccgcaaaattagcaacagtgaaaaaataccaattgaccagtgaaattgcttaatattatttttatatctatttcaagttgttgggaaaggtgtccagtgtgtatctATGATTAAAAAACCTAATTCCAAGACGCATAAGGAAAAATATGAGGGAAAATACAAAACA
+2fr:                                                                                                                                                                                                ·M··I··K··K··P··N··S··K··T··H··K··E··K··Y··E··G··K··Y··K··T·

              1457                                                                1535     EXON2
              |                                                                   |
 0.41 gcggaagGTtggtgc                                          0.98 tttaattatttcaccttgcAGattggcatcgaagcgagaag
DNA:  AATGGCGGAAGgttggagcttgtctgactcacaatggcttccggaaaattcccagaaatgctgcaaatttaattatttcaccttgcagATTGGCATCGAAGCGAGAAGCAGAGAGACATCCATCAACACATGAAGACACCATTAAAAAATCCCATCATCATCGTGATGAATTAGGGAAGAAAGATGAAAGGAAGGCTTATGAAGAAGAATTTCCTAAATTTGGGCATGTTCCATCTACATC
+2fr: ·N··G··G··R                                                                         ··L··A··S··K··R··E··A··E··R··H··P··S··T··H··E··D··T··I··K··K··S··H··H··H··R··D··E··L··G··K··K··D··E··R··K··A··Y··E··E··E··F··P··K··F··G··H··V··P··S··T··S
                                         1725
                                         |
                              0.30 aggacggGTaatggc
DNA:  ACGGGATTCCCGGGACGGCTTTCAACACAAAAGGACGGgtaatggcatgaagaaatcttctatcaatgacgatattaggagaacatgtcacaagtcagagaatttgatgagtattttttgaaataagaaattggaatattttgtgaattttcatgagatttttgagacttttaaagtaaaaagagagatcatcaatatgcccaagtaatcttgagagcagaTCATCAATCAACAaagacat
+2fr: ··R··D··S··R··D··G··F··Q··H··K··R··T··

DNA:  ttcagagctggacaaccgaagctgaaggaaccgaggagatgagtagtttccaggatgtaaaaaggtaagtttaaaatatttatcgtaacttcgtctactttcagaatatccaggaattacacagaacatgtccaagggataccttcctctcaagattCTTTCAATCAACAttcaacaaattccaattggcgggagaaagggtagtgaaaaattaatccaatgcacatcaaatctccaatca

                                                                                                                                                                                                           2363     EXON3
                                                                                                                                                                                                           |
                                                                                                                                                               0.85 ttttgcaataaactcccacAGgtgatggtagggaatcctct
DNA:  acaagttGCCCCAATCAACAcatttaaattTCTTCAATCAACAccatttcctggaggacgttTCTTCAATCAACAcgtagattttgctcaaataacttaatttttattaaatttttatcaaacaaaaatttaaatttttgtggagttatttaaagaagttctcatgtaaattttgcaataaactcccacagGTGATGGTAGGGAATCCTCTAAGCACCGAAGAGGTGCTGCCAAGAGAG
+3fr:                                                                                                                                                                                                            G··D··G··R··E··S··S··K··H··R··R··G··A··A··K··R··

            2418                                                          2480     EXON4
            |                                                             |
 0.77 ttcttcaGTaagtaa                                     0.66 taattgtgagagtttttgtAGaattcttcagagactgacga
DNA:  ATTCTTCAgtaagtaaaaaaaagtacttaagccgggagtagagagatttaattgtgagagtttttgtagAATTCTTCAGAGACTGACGATGAATCCCTCAAATCTTCGAGGAAAAGGTCAGAATCTCGCCGGAGACGATCTAAAAGTCGTAGCCGAAGTCTGGATGACTCGTACAGGAGACGTCAAACCTCCACGAATCGTCGATATTCGCCACGGCGCCGGCCAAAGTCTGTTTCTCCTC
+3fr: D··S··S·                                                              ·N··S··S··E··T··D··D··E··S··L··K··S··S··R··K··R··S··E··S··R··R··R··R··S··K··S··R··S··R··S··L··D··D··S··Y··R··R··R··Q··T··S··T··N··R··R··Y··S··P··R··R··R··P··K··S··V··S··P··

DNA:  CTCGTGAGAAAGTTCCCTACTATGCAGATGCCCAACGCGAGAGGGATCGCCTGAGGGAGAAATATGGAAGTGATGGGAGGGCTAGGAGAAGGAGCAGCAGTCGCGAGAGACGCAGAAGGCGTTCTAGAGAAAGGAGATCCAAGACTCCAGTCCGAGAAACCACAAAAATTGTTACAGTTCCTGTTCCTGTTCCTGTCTACCCGTCAATATATCCGGATGGTTCCATGTATGACTGGTACGA
+3fr: P··R··E··K··V··P··Y··Y··A··D··A··Q··R··E··R··D··R··L··R··E··K··Y··G··S··D··G··R··A··R··R··R··S··S··S··R··E··R··R··R··R··R··S··R··E··R··R··S··K··T··P··V··R··E··T··T··K··I··V··T··V··P··V··P··V··P··V··Y··P··S··I··Y··P··D··G··S··M··Y··D··W··Y··D

                                                                                                                                                                                                                  3094
                                                                                                                                                                                                                  |
DNA:  TCCTTCTTGGCAAGCAGGACTCTTTTCTCAGGGTGTCATGCCTCCAAGACACATTCCACGCCCAATGCGTCCTCATGCCCCATTCCCTGCCTCACCCTTCATGGTGGATCCTTTCCATCGGCCAATGAGACCAGGAATGCCACCTCCAAGACCCTTCCGACCTCCCCAAGGATTCCCTCCTCCCCGATTCCACAGACCCCATTGAaatgaaagtctgaaaatactac
+3fr: ··P··S··W··Q··A··G··L··F··S··Q··G··V··M··P··P··R··H··I··P··R··P··M··R··P··H··A··P··F··P··A··S··P··F··M··V··D··P··F··H··R··P··M··R··P··G··M··P··P··P··R··P··F··R··P··P··Q··G··F··P··P··P··R··F··H··R··P··H··*·
```

**Figure S13. Manually-curated *P. duboscqi transformer* gene model.** Exonic sequences are indicated by black cases; black upper cases indicate coding sequences. Intronic sequences are indicated by gray lower cases. Start and stop codons are highlighted in green and red, respectively. In yellow boxes, putative TRA/TRA-2 binding sites. Exon start/end positions in the scaffolds are indicated. Intron boundaries were predicted using transcripts vs genome alignments and confirmed by *de novo* prediction with Berkeley BDGP Splice Site Prediction Tool with default parameters (http://www.fruitfly.org/seq_tools/splice.html); prediction scores are indicated in red.
