## Additional_file_4_Figure_S14 for "Genomics and transcriptomics to unravel sex determination pathway and its evolution in sand flies"

```
                                                                                DIPTERA domain
PpeTRA      MSTIRKSSSTAHKERYETK-HNNGGK|MATKREAERRPSTRQDSFKKSHRE----------      49
PduTRA      --MIKKPNSKTHKEKYEGKYKTNGGR|LASKREAERHPSTHEDTIKKSHHHRDELGKKDER      58
PpaTRA      --MIKKPSSKTHKEKYEGKYKTNGGR|SASKREAERHPSTHEDTIKKSHHHRDELGKKDER      58
PbeTRA      --MIKKPSSRTHKEKYEGKYKTNGGR|SASKREAERHPSTHEDTIKKSHHHRDELGKKDER      58
               *:* .* :***:** * :.***:   *:******:***::*::****:.

PpeTRA      ---------ESHIPSTSRTPRDGSQHKRTGDRRESSKHRKDASRRDSSSSSDSEDSDKEL      100
PduTRA      KAYEEEFPKFGHVPSTSRDSRDGFQHKRTGDGRESSKHRRGAAKRDSSNSSETD---DES      115
PpaTRA      KTYEDEFPKLGHVPSTSRDSRDGFQHKRTGDGKESSKHRRGAAKRDSSNSSETD---DES      115
PbeTRA      KSYEDDFPKLGHVPSTSRDTRDGFQHKRTGDGRESSKHRRGAAKRDSSNSSETD---SES      115
                      .*:*****  *** ******* :******:.*::****.**:::   .*

PpeTRA      GKSSRRRENSRRRRSKSRSRSLDDSRRRM----SRRSPPRRRPKSETPPRESKPYYADAQ      156
PduTRA      LKSSRKRSESRRRRSKSRSRSLDDSYRRRQTSTNRRYSPRRRPKSVSPPREKVPYYADAQ      175
PpaTRA      LKSSRKRPESRRRRSKSRSRSLDDSYRRRQASTSRRSSPRRRPKSVSPPREKVPYYADAQ      175
PbeTRA      LKSSRKRPESRRRRSKSRSRSLDDSYRRRQTSTSRRSSPRRRPKSVSPPREKVPYYADAQ      175
             ****:* :**************** **     .**  ******* :****. *******

PpeTRA      RERDRLREKYGRDEERHRSR--RRSYSRERRRRRSRTPVRETTKIVTV--PVPVYPTFYP      212
PduTRA      RERDRLREKYGSDGRARRRSSSRERRRRRSRERRSKTPVRETTKIVTVPVPVPVYPSIYP      235
PpaTRA      RERDRLREKYGSDGRTRRRSSSRERRRRRSRDRRSKTPVRETTKIVTVPVPVPVYPSIYP      235
PbeTRA      RERDRLREKYGSDGRMRRRSSSRERRRRRSRDRRSKTPVRETTKIVTVPVPVPVYPSIYP      235
            *********** * . :*    *.   *. * ***:************  ******::**

PpeTRA      DGSVYEWYDPAWPGGPRPM------LSQPPMRPPGFYPGGHFMVDPFHRPMRPGMAPPRP      266
PduTRA      DGSMYDWYDPSWQAGLFSQGVMPPRHIPRPMRPHAPFPASPFMVDPFHRPMRPGMPPPRP      295
PpaTRA      DGSMYDWYDPSWQTGLFSQAVMPPRHIPRPMRPHAPFPTSPFMVDPFHRPMRPGMPPPRP      295
PbeTRA      DGSMYDWYDPSWQTGLFPQGVMPPRHIPRPMRPHAPFPTSPFMVDPFHRPMRPGMPPPRP      295
            ***:*:****:*  *              **** . :* . ************** ****

PpeTRA      FRPPMGFPQPRFQRPH  282
PduTRA      FRPPQGFPPPRFHRPH  311
PpaTRA      FRPPQGFPPPRFHRPH  311
PbeTRA      FRPPQGFPPPRFHRPH  311
            **** *** ***:***
```

**Figure S14. Multiple sequence alignment of TRA proteins in *Phlebotomus* spp.** Sequence alignment of TRA proteins of *P. perniciosus* (PpeTRA), *P. duboscqi* (PduTRA), *P. papatasi* (PpaTRA) and *P. bergeroti* (PbeTRA). The DIPTERA domain is highlighted in grey. The position of the sex-specifically regulated splicing site between exon2 and exon3 is indicated by the pipe symbol (|). Gaps were introduced in the alignment to maximize similarity. The protein sequences alignment was performed using the Clustal-Omega software (1.2.4).
