## Additional_file_5_Figure_S15 for "Genomics and transcriptomics to unravel sex determination pathway and its evolution in sand flies"

**A)**

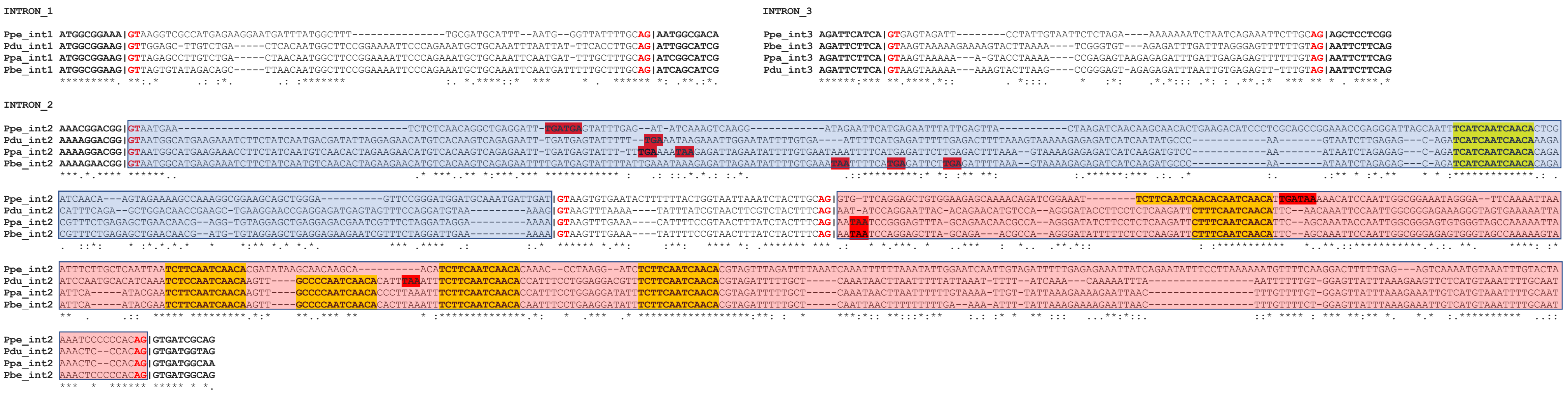


**B)**

| Intron 1 | | | | | | | |
|---|---|---|---|---|---|---|---|
| Species | 5' Donor SS GTRAGY | Identity | BDGP SS Pred. Score | 3' Acceptor SS YYYYYYYYYYYYNYAG | Identity | Cons. n° of py = 8.01+/-1.98 n° o py | BDGP SS Pred. Score |
| Ppe | GTAAGG | 5/6 | 0.64 | ATGGGTTATTTTGCAG | 10/16 | 7 | 0.85 |
| Ppa | GTTAGA | 4/6 | 0.49 | TGATTTTGCTTTGCAG | 13/16 | 9 | 0.97 |
| Pbe | GTTAGT | 5/6 | 0.82 | GATTTTTGCTTTGCAG | 13/16 | 9 | 0.98 |
| Pdu | GTTGGA | 3/6 | 0.41 | TTATTTCACCTTGCAG | 14/16 | 10 | 0.97 |

| Intron 2 | | | | | | | |
|---|---|---|---|---|---|---|---|
| Species | 5' Donor SS GTRAGY | Identity | BDGP SS Pred. Score | 3' Acceptor SS YYYYYYYYYYYYNYAG | Identity | Cons. n° of py = 8.01+/-1.98 n° o py | BDGP SS Pred. Score |
| Ppe | GTAATG | 5/6 | 0.42 | TAAAATCCCCCCACAG | 12/16 | 8 | 0.91 |
| Ppa | GTAATG | 5/6 | 0.30 | CAATAAACTCCCACAG | 11/16 | 7 | 0.77 |
| Pbe | GTAATG | 5/6 | 0.41 | ATAAACTCCCCCACAG | 12/16 | 8 | 0.85 |
| Pdu | GTAATG | 5/6 | 0.30 | CAATAAACTCCCACAG | 11/16 | 7 | 0.85 |

| Intron 3 | | | | | | | |
|---|---|---|---|---|---|---|---|
| Species | 5' Donor SS GTRAGY | Identity | BDGP SS Pred. Score | 3' Acceptor SS YYYYYYYYYYYYNYAG | Identity | Cons. n° of py = 8.01+/-1.98 n° o py | BDGP SS Pred. Score |
| Ppe | GTGAGT | 6/6 | 0.58 | TCAGAAATTCTTGCAG | 11/16 | 7 | 0.92 |
| Ppa | GTAAGT | 6/6 | 0.77 | AGAGAGTTTTTTGTAG | 10/16 | 6 | 0.55 |
| Pbe | GTAAGT | 6/6 | 0.77 | AGGGAGTTTTTTGTAG | 10/16 | 6 | 0.69 |
| Pdu | GTAAGT | 6/6 | 0.77 | GTGAGAGTTTTTGTAG | 10/16 | 6 | 0.66 |

| ms1_male-specific | | | |
|---|---|---|---|
| Species | 5' Donor SS GTRAGY | Identity | BDGP SS Pred. Score |
| Ppe | GTAAGT | 6/6 | 0.89 |
| Ppa | GTAAGT | 6/6 | 0.96 |
| Pbe | GTAAGT | 6/6 | 0.96 |
| Pdu | GTAAGT | 6/6 | 0.99 |

| fs1_female-specific | | | | |
|---|---|---|---|---|
| Species | 3' Acceptor SS YYYYYYYYYYYYNYAG | Identity | Cons. n° of py = 8.01+/-1.98 n° o py | BDGP SS Pred. Score |
| Ppe | TTAAATCTACTTGCAG | 12/16 | 8 | 0.94 |
| Ppa | CTTCGTCTACTTTCAG | 14/16 | 10 | 0.91 |
| Pbe | CTTTATCTACTTTCAG | 14/16 | 10 | 0.91 |
| Pdu | CTTTATCTACTTTCAG | 14/16 | 10 | 0.84 |

**Figure S15. Multiple alignment of *tra* introns in *Phlebotomus* spp. A)** All introns exhibit GT/AG consensus terminal dinucleotides (in red). Exonic flanking sequences are indicated by bold cases; exon/intron boundaries are indicated by the pipe | symbol. Azure box represents the male-specific ms1 exon. Pink box represents the female-specific fs1 exon. In frame stop codons are highlighted in red. Putative TRA/TRA-2 binding sites are highlighted in yellow. Intron boundaries were predicted using transcripts vs genome alignments for *P. perniciosus* and *P. papatasi* and *de novo* predicted by Berkeley BDGP Splice Site Prediction Tool with default parameters (http://www.fruitfly.org/seq_tools/splice.html) for *P. bergeroti* and *P. duboscqi*. **B)** Splicing site (SS) score values and consensus match for all the splicing sites are reported. Bold letters indicate match with the consensus sequences. Strong SS were highlighted in green and weak SS in red. Consensus number of pyrimidine of 3' Acceptor SS in sand flies has been calculated by the tabulation of 25.000 random acceptor SS extracted from the *P. papatasi* PpapI1.4 gene set using the Biomart tool of VectorBase. (M=A or C; W=A or T; R=A or G; Y=C or T; N=any nucleotide).
