## Additional_file_6_Figure_S16-S17 for "Genomics and transcriptomics to unravel sex determination pathway and its evolution in sand flies"

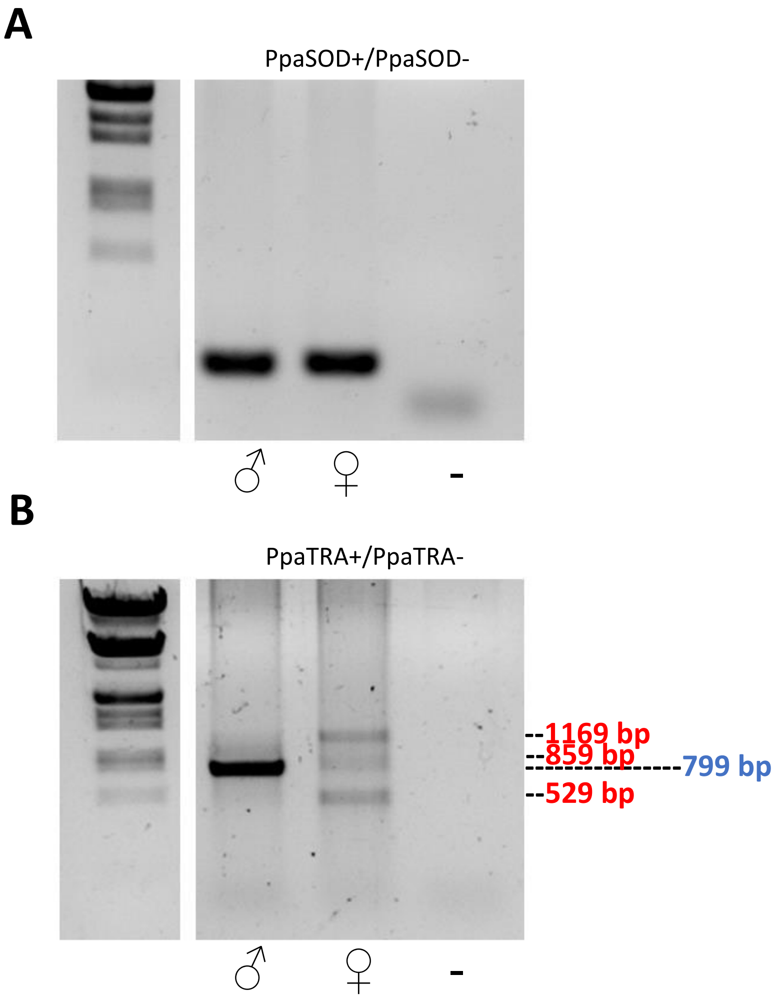


**Figure S16. *tra* gene expression at adult stage in *P. papatasi*.** RT-PCR analysis on total RNA extracted from adult males and females of *P. papatasi*. (A) Control RT-PCR with primer pair PpaSOD+/PpaSOD-. (B) RT-PCR with primer pair PpaTRA+/PpaTRA-. λDNA digested with EcoRI and HindIII endonucleases (Marker III – Sigma Aldrich) was utilized as molecular weight marker.

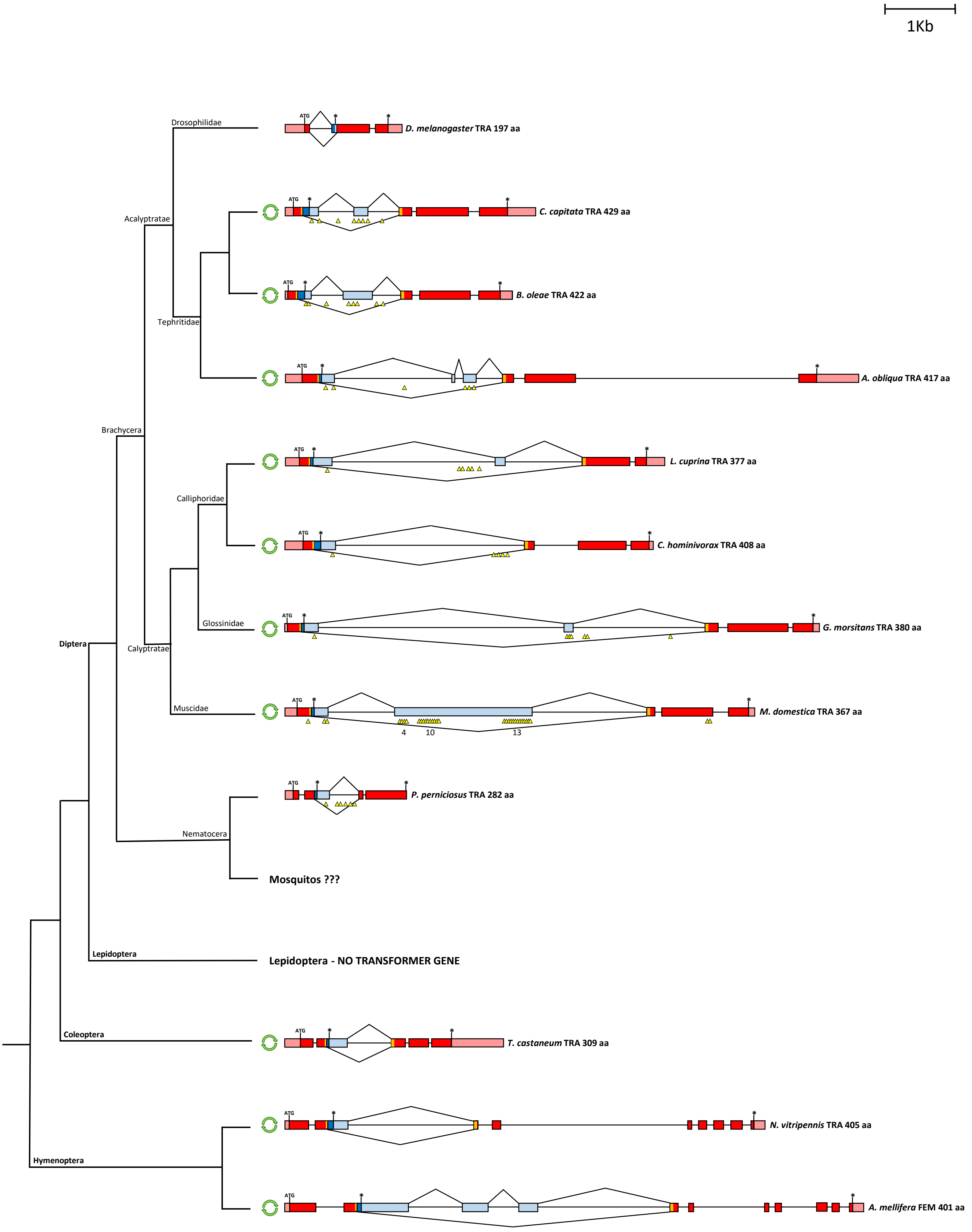


**Figure S17. Phylogenetic relationship, genomic structure and sex-specific splicing regulation of *transformer* orthologues in insects.** Pink boxes indicate untranslated regions; azure boxes indicate untranslated males-specific regions; red boxes indicate female-specific exon/regions; blue boxes indicates male-specific coding regions, yellow boxes indicate the position of the *tra* region encoding for the TRACAM domain. Translational start (ATG) and stop (*) sites are indicated. Yellow triangles indicate the position of the putative TRA/TRA-2 binding sites (for *M. domestica* is reported also the number of TRA/TRA-2 bindings sites). Green double circular arrows indicates autoregulation of the *tra* ortholog.
