## Additional_file_7_Figures_S18-S20 for "Genomics and transcriptomics to unravel sex determination pathway and its evolution in sand flies"

### *P. papatasi tra-2* partial gene model

```
Scaffold1530_Length=36934

                                                                                                                                                                                       32651
                                                                                                                                                                32624                  |
                                                                                                                                                                |              0.99 ctataagGTaagttc
DNA:  atttcgaaaaaatgtgcatacacatccgagatacgcagccattttcgaaatgtcactagtgtgtttttgtaaatttttgcatctctagagccaaaaaagccaattttctctcatttttcacacatctcataatagtttttcccactaaaatagctacattgtgATGAGTACACGTACCTACTATAAGGTAAGTTCAGAAAATTCAATTGAGTTTGCAAATAAAATTCCCAAACAACCGGC
+2fr:                                                                                                                                                            ·M··S··T··R··T··Y··Y··K··V··S··S··E··N··S··I··E··F··A··N··K··I··P··K··Q··P··A

                                                                                                                                                                                                                                       32933
                                                                                                                                                                                                                                       |
                                                                                                                                                                                                                             ??? ccaaaaaGTacaaat
DNA:  TTGCTGTGCGGTCGCCATGGCGAAAAAGTGCATCAAAGACTGCGATGCATTTTGTCCCCAAGGTTCTCTTTTGTTTTTTTTTTCCTTGCCTTGAAGAAGCACACCTATCTGATCGATCAAACAATCCCTTCGTTAGTTCAATATCTCAGCTAAAGGGAAAGGGTTAGACAGTGAGTGTCCTTGAGGAAAAAAAAGAGTCCAAAATCTTACTCATGACATCTGACCAAAAAgtacaaataag
+2fr: ··C··C··A··V··A··M··A··K··K··C··I··K··D··C··D··A··F··C··P··Q··G··S··L··L··F··F··F··S··L··P··*··R··S··T··P··I··*··S··I··K··Q··S··L··R··*··F··N··I··S··A··K··G··K··G··L··D··S··E··C··P··*··G··K··K··E··S··K··I··L··L··M··T··S··D··Q··K·

                                                          33002
                                                          |
                                       0.90 cccttttctcccattttccAGaactcaatgtcaagaagtcc
DNA:  tgaaaaaaaatataaaacaatttcataactaaattacccttttctcccattttccagAACTCAATGTCAAGAAGTCCCAGCGAGGAACGACACTATATCTCTCGAGCTGGTTCATCGCGAGAGAATGGCGACTACAGGAAGTCCCGCAAATCGTCTTCCCGCAGTCGAGCCCGCTACAGATCTCGCTCACGTTCACGGTCTCGTTCTGAGAAGCGCTACAGAAGCAGGTCACCGTCCCGC
+1fr:                                                     ·N··S··M··S··R··S··P··S··E··E··R··H··Y··I··S··R··A··G··S··S··R··E··N··G··D··Y··R··K··S··R··K··S··S··S··R··S··R··A··R··Y··R··S··R··S··R··S··R··S··R··S··E··K··R··Y··R··S··R··S··P··S··R·

                                                                 33250                                       33303
                                                                 |                                           |
                                                        0.77 aagtcgaGTaagtgc                ??? ttataaatgacatagaaccAGgtcaattcacaggatcatc
DNA:  CACGATAGGAAGCGCTCCTATTCCAGGAGCCCCAATTCCTCTCGTCGCAGACACGTAGCAAGTCGAgtaagtgctgatgcaaatcaattggatgtgattataaatgacatagaaccagGTCAATTCACAGGATCATCCCCCAAAGTCCAGATGTCTAGGAGTCTTTGGACTCAGTGATGCAACAACTGAAGACCAGATTTATCAAATTTTCTCCAAATTTGGAACAATACACCGTACTCA
+1/2f:·H··D··R··K··R··S··Y··S··R··S··P··N··S··S··R··R··R··H··V··A··S··R·                                       ·V· N··S··Q··D··H··P··P··K··S··R··C··L··G··V··F··G··L··S··D··A··T··T··E··D··Q··I··Y··Q··I··F··S··K··F··G··T··I··H··R··T··Q

                           33443
                           |
                0.76 tgcaatgGTaattcc
DNA:  AATAATTATGGATGCAATGgtaattcctaattcaataatacaaaatatttcccccagaatcccctcaacttcccagtgatgagttagccgggaagttgaaaaccacatacgcgagattatctgattttgggtagataaatatccggaaaatttagaatgatatccacactcatggcttgctaagagtttcaattgggttttcattaaaggcattccggaacattttctctcaatctctat
+1fr: ··I··I··M··D··A··M·

                                               33708
                                               |
                       0.64 ggggatttgttttcgttgcAGacgggatgttcacggggctt
DNA:  tccagagtagattttaattatagggggatttgttttcgttgcagACGGGATGTTCACGGGGCTTCTGTTTTGTCTACTTTGCCAATGCTGATGATGCCAAAGTGGCAAAGGATAATTGCTCAGGAATGGAACTCGATGGGAGACGTATTCGAGTGGATTATTCCATTACTCAGAGACCTCATACTCCAACTCCAGGAGTCTACATGGGACAACCAACGGGCAGAGGACGAGAGAATCGAGA
+2fr:                                          ·T··G··C··S··R··G··F··C··F··V··Y··F··A··N··A··D··D··A··K··V··A··K··D··N··C··S··G··M··E··L··D··G··R··R··I··R··V··D··Y··S··I··T··Q··R··P··H··T··P··T··P··G··V··Y··M··G··Q··P··T··G··R··G·· R··E··N··R··

                          33922
                          |
               0.87 gaaatgaGTaagtac
DNA:  TCGTAGTCGCAGAAATGAgtaagtactgccttttaaatcaactttgtttattttcataatcgattcagttcaagtgaatccaataacactgcaaagatgattttgtccctggattctcatgctatttttttctattgttagagtcaaatgatttacgaaaatacttatcattttgatttttaagtattaaataatataataaaagctaaattaaataaataaattttagtattaaataat
+2fr: D··R··S··R··R··N··

DNA:  ataataatagcttgtactgcctaattgcgagaaaaaaaaaacaataacactcgaaattgtgaagtttttctttagttggctgagtgctcgttttccgtttgaacctcaatatcgaggttaaaaaacaatctgagacaaaagttgtacatggacgaaaatgtagctgattaaattctctatcaaacccttaaaacagattgttaggaacagtcctaattttcgagaaatttttgattaaat

DNA:  ttgctaaaaagttcgattttccatgttttctgacatttatgagactacagcgcccctggtgtcagttgtacgaacttcaactgttcagaggatttgtcgggcatgtaaaggagaccaaattattccatagtgagaaataaatcggttcagcagaatcggagatataggcacccaagtatcaaaaaacgacaaaaacggttttgcctagattccagnnnnnnnnnnnnnnnnnnnn

DNA:  nnnnnnnnnnnnnnnnnnnnnnnnnnnnnnnnnnnnnnnnnnnnnnnnnnnnnnnnnnnnnnnnnnnnnnnnnnnnnnnnnnnnnnnnnnnnnnnnnnnnnnnnnnnnnnnnnnnnnnnnnnnnnnnnnnnnnnnnnnnnnnnnnnnnnnnnnnnnnnnnnnnnnnnnnnnnnnnnnnnnnnnnnnnnnnn
```

**Figure S18. Manually-curated *P. papatasi tra-2* partial gene model.** Exonic sequences are indicated by black cases; black upper cases indicate coding sequences. Intronic sequences are indicated by gray lower cases. Start and stop codons are highlighted in green and red, respectively. Underlined cases indicate putative alternatively spliced exon, predicted on the basis of sequence conservation with the corresponding exon of *P. perniciosus tra-2* gene. It is interesting to observe the sequence of the putative alternative N-terminus (MTSDQK) which is identical between the two *Phlebotomus* species. Exon start/end positions in the scaffold/s are indicated. Intron boundaries were predicted using transcripts vs genome alignments and confirmed by *de novo* prediction with Berkeley BDGP Splice Site Prediction Tool with default parameters (http://www.fruitfly.org/seq_tools/splice.html); prediction scores are indicated in red. Interestingly, the BDGP Splice Site Prediction Tool fails to predict the 5' alternative donor splicing site downstream the putative alternatively spliced exon of *P. papatasi tra-2*, indicating that this splicing site is suboptimal and it could need an activation by unknown factors.

### *P. bergeroti tra-2* partial gene model

**Contig_173972_Lenght=4440**

```
DNA:  ataataatagcttgtactgcctaattgcgagaaaaaaaaaacaataacactcgaaattgtgaagtttttctttagttggctgagtgctcgttttccgtttgaacctcaatatcgaggttaaaaaacaatctgagacaaaagttgtacatggacgaaaatgtagctgattaaattctctatcaaacccttaaaacagattgttaggaacagtcctaattttcgagaaatttttgattaaat

                                                                                                        2942
                                                                                    2966                  |
                                                                                    |          0.99 ctataagGTaagttc
DNA:  tgcatctctagagccaaaaaagccaattttctcccatttttcacacttttcataatagtttttcccactaaaatagctacattgtgATGAGTACTCGTACCTACTATAAGGTAAGTTCACAAAAGTCAATTTGTCTTTCCATGAATAAAATTCCCAAACAACCGGCTTGCCGTGTGGTCGCCATGGCGAAAAAGTGCATCAAAGACTGCGATGCATTTTGTCCCCAAGGTTCTCTTTTGT
-3fr                                                                                    ·M··S··T··R··T··Y··Y··K··V··S··S··Q··K··S··I··C··L··S··M··N··K··I··P··K··Q··P··A··C··R··V··V··A··M··A··K··K··C··I··K··D··C··D··A··F··C··P··Q··G··S··L··L··

                                                                                                                                        2656                                                         2590
                                                                                                                                          |                                                            |
                                                                                                                              ??? ccaaaaaGTacaaat                                     0.94 ctcttttctcccattttccAGaactcaatgtcgagaagtc
DNA:  TTTTTTTTCCTTGCCTTGAAGAAGTACACCTATCTGATCGATCAAACAATCCCTTCGTTAGTTCAATATCTCAGCTATAGGAAAAGGGTTAAACTGTGAGTGTCCTTGAGGAAAAAAGAGGAGTCCAAAATCTTGCTCATGACATCTGACCAAAAAgtacaaataaactacgaaaaaatatgaaataatttctgaaattgctcttttctcccattttctagAACTCAATGTCGAGAAGTC
-1/3f: ·F··F··F··L··A··L··K··K··Y··T··Y··L··I··D··Q··T··I··P··S··L··V··Q··Y··L··S··Y··R··K··R··V··K··L··*··V··S··L··R··K··K··E··E··S··K··I··L··L··M··T··S··D··Q··K·                                                                    ·N··S··M··S··R··S··

                                                                                                                                                                                                        2341
                                                                                                                                                                                                          |
      c                                                                                                                                                                                          0.77 aagtcgaGTaagtgc
DNA:  CCAGCGAGGAACGACACTACATCTCTCGAGCAGGTTCATCCCGAGAGAATGGAGACTACAGAAAGTCCCGCAAATCGTCTTCTCGTAGTCGAGGACGCTACAGATCTCGCTCACGTTCACGATCTCGTTGTGAGAAGCGTTACAGAAGCAAGTCACCGCCCCGCCACGATAGGAAACGCTCCTACTCCAGGAGCCCCAATTCTTCCCGTCGCAGACACGTAGCAAGTCGAgtaagtgctg
-3fr: P··S··E··E··R··H··Y··I··S··R··A··G··S··S··R··E··N··G··D··Y··R··K··S··R··K··S··S··S··R··S··R··G··R··Y··R··S··R··S··R··S··R··S··R··C··E··K··R··Y··R··S··K··S··P··P··R··H··D··R··K··R··S··Y··S··R··S··P··N··S··S··R··R··R··H··V··A··S··R·

                                               2288                                                                                                                                     2148
                                                 |                                                                                                                                        |
                        ??? ttataaatgacatagaaccAGgtcaattcacaggatcatc                                                                                                     0.72 tgcaatgGTaattca
DNA:  atgcaaatcaatttgatgcgattataaatgacataggaccagGTCAATTCACAGGATCATCCTCCAAAGTCCAGATGTCTAGGGGTCTTTGGACTCAGTGATGCAACCACTGAAGACCAGATTTATCAAATTTTCTCCAAATTTGGAACTATACACCGTACTCAAATAATTATGGATGCAATGgtaattcaataatacaatatatttctccccagaatcccctcaacttcccagtgatta
-1fr:                                            ·V··N··S··Q··D··H··P··P··K··S··R··C··L··G··V··F··G··L··S··D··A··T··T··E··D··Q··I··Y··Q··I··F··S··K··F··G··T··I··H··R··T··Q··I··I··M··D··A··M·

                                                                                                                                                                                                1884
                                                                                                                                                                                                  |
                                                                                                                                                                   0.66 gggaaattgtaatcatttcAGacggggtgttcacggggctt
DNA:  cctagccgggaagttgaaatctcatatgtgagatatctgtctgattttggtagataaatatccggaaaatttagaatgatacccacactcatagcttgctaagagtttcaattgggttttcattaaaggcattccgggacatttttcactcaatcaattccacagagtagtagattttaattattgggaaattgtaatcatttcagACGGGGTGTTCACGGGGCTTTTGTTTTGTCTACT
-3fr:                                                                                                                                                                                             ·T··G··C··S··R··G··F··C··F··V··Y··

                                                                                                                                                                          1670
                                                                                                                                                                            |
                                                                                                                                                               0.87 gaaatgaGTaagtat
DNA:  TTGCCAATGCTGATGATGCCAAAGTGGCAAAGGATAATTGCTCAGGAATGGAACTCGATGGGAGACGTATTCGAGTGGATTATTCCATTACTCAGAGACCTCATACTCCAACTCCAGGAGTCTACATGGGACAACCAACGGGCAGAGGACGAGAGAATCGAGATCGTAGTCGCAGAAATGAgtaagtatcactctgaatcaattttgtttattttcgtatcttattcggttcaagtgaat
-3fr: F··A··N··A··D··D··A··K··V··A··K··D··N··C··S··G··M··E··L··D··G··R··R··I··R··V··D··Y··S··I··T··Q··R··P··H··T··P··T··P··G··V··Y··M··G··Q··P··T··G··R··G··R··E··N··R··D··R··S··R··R··N··

DNA:  ccaatattctctaaatcaaataaaaaatacgcctcctaagatttttaattcggaaaaagaaacagtggtaaacccagatagacttttatgtggcttactgtgttatcacacttgcacatt
```

**Figure S19. Manually-curated *P. bergeroti tra-2* partial gene model.** Exonic sequences are indicated by black cases; black upper cases indicate coding sequences. Intronic sequences are indicated by gray lower cases. Start and stop codons are highlighted in green and red, respectively. Underlined cases indicate putative alternatively spliced exon, predicted on the basis of sequence conservation with the corresponding exon of *P. perniciosus tra-2* gene. It is interesting to observe the sequence of the putative alternative N-terminus (MTSDQK) which is identical between the two *Phlebotomus* species. Exon start/end positions in the scaffold/s are indicated. Intron boundaries were predicted using transcripts vs genome alignments and confirmed by *de novo* prediction with Berkeley BDGP Splice Site Prediction Tool with default parameters (http://www.fruitfly.org/seq_tools/splice.html); prediction scores are indicated in red. Interestingly, the BDGP Splice Site Prediction Tool fails to predict the 5' alternative donor splicing site downstream the putative alternatively spliced exon of *P. bergeroti tra-2*, indicating that this splicing site is suboptimal and it could need an activation by unknown factors.

### *P. duboscqi tra-2* partial gene model

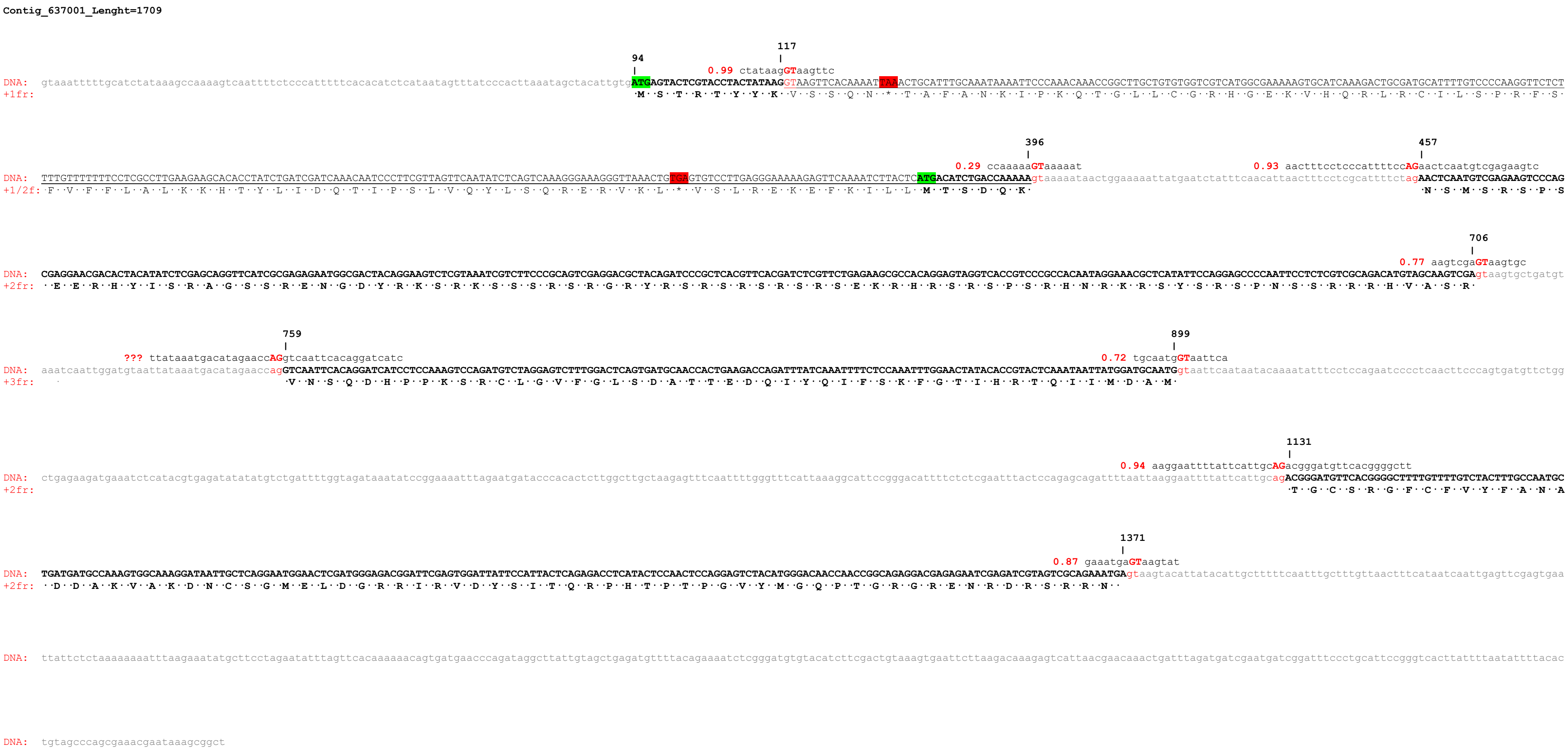


**Figure S20. Manually-curated *P. duboscqi tra-2* partial gene model.** Exonic sequences are indicated by black cases; black upper cases indicate coding sequences. Intronic sequences are indicated by gray lower cases. Start and stop codons are highlighted in green and red, respectively. Underlined cases indicate putative alternatively spliced exon, predicted on the basis of sequence conservation with the corresponding exon of *P. perniciosus tra-2* gene. It is interesting to observe the sequence of the putative alternative N-terminus (MTSDQK) which is identical between the two *Phlebotomus* species. Exon start/end positions in the scaffold/s are indicated. Intron boundaries were predicted using transcripts vs genome alignments and confirmed by *de novo* prediction with Berkeley BDGP Splice Site Prediction Tool with default parameters (http://www.fruitfly.org/seq_tools/splice.html); prediction scores are indicated in red. Interestingly, the BDGP Splice Site Prediction Tool predicts the 5' alternative donor splicing site downstream the putative alternatively spliced exon of *P. duboscqi tra-2* as very weak, indicating that this splicing site is suboptimal and it could need an activation by unknown factors.
