## Additional_file_8_Figures_S21-S22 for "Genomics and transcriptomics to unravel sex determination pathway and its evolution in sand flies"

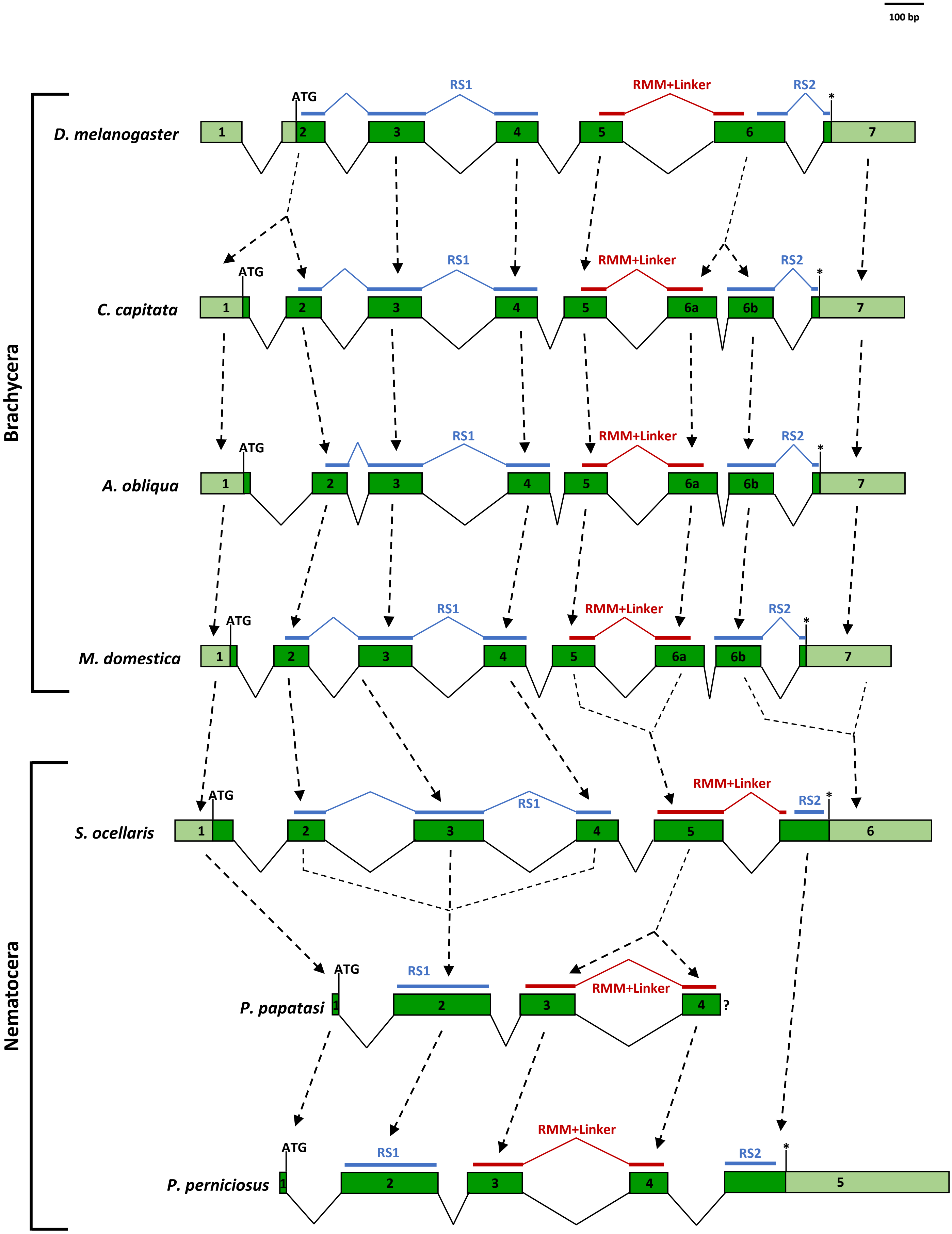


**Figure S21. Comparison of genomic structures of diptera *tra-2* genes.** Green boxes represent coding regions while light green boxes represents untranslated regions. Asterisks indicate the position of stop codons. The exon portions encoding for RS domains, RRM domain and linker region are indicated. Introns not to scale. *P. perniciosus tra-2* intron-exon organization has been predicted by comparison with *P. papatasi* partial *tra-2* gene.

```
                              RS domain 1
LumTRA-2      ?--RNLFPQSSD-SSGYDR-RHSRTKKKRENGHHKKSSKS------RRRHRSRSHSK---  46
LneTRA-2      ?--RNLFPQSSD-SSGDDR-RYSRTKKKRENGHHKKSSKS------RRRHRSRSHSK---  46
PpeTRA-2A     --MTSDQKNSMSRSPSEGRSYILKGGSSRENNGYRRSDRSRSRR--RYRSRSRSHSQRRH  56
PbeTRA-2      MSTRTYYKNSMSRSPSEERHYISRAGSSRENGDYRKSRKSSSRSRGRYRSRSRSRS--RC  58
PpaTRA-2      MSTRTYYKNSMSRSPSEERHYISRAGSSRENGDYRKSRKSSSRSRARYRSRSRSRS--RS  58
PduTRA-2      MSTRTYYKNSMSRSPSEERHYISRAGSSRENGDYRKSRKSSSRSRGRYRSRSRSRS--RS  58
                  .   :* . * .  *    :  ..***. :::* :*      * * ****:*

                                                          RRM
LumTRA-2      -RRHYSRSPTRTTRKRHISRSPVPT--------YKKYEQPEEGRCLGVFGLSMHTTEKHV  97
LneTRA-2      -RRHYSRSPTRTIRKRHISRSPVPS--------YKKYEQPEEGRCLGVFGLSMHTTEKHV  97
PpeTRA-2A     RSRY-SGSPVRNDRKHSHSRSPHSSRRRHVG----SRDHPQQSRCLGVFGLSLITTKQQI  111
PbeTRA-2      EKRYRSKSPPRHDRKRSYSRSPNSSRRRHVASRVNSQDHPPKSRCLGVFGLSDATTEDQI  118
PpaTRA-2      EKRYRSRSPSRHDRKRSYSRSPNSSRRRHVASRVNSQDHPPKSRCLGVFGLSDATTEDQI  118
PduTRA-2      EKRHRSRSPSRHNRKRSYSRSPNSSRRRHVASRVNSQDHPPKSRCLGVFGLSDATTEDQI  118
                *: * ** *  **:  ****  :          . ::* :.*******   **:.::
                                                         RNP-1

LumTRA-2      HEIFSKFGPIERTQIIVDSKTGRSRGYCFVYFENTEDAKVAKDQCTGIEIDDRRIRVDYS  157
LneTRA-2      HEIFAKFGPIERTQIIVDSKTGRSRGYCFVYFKNTEDAKVAKDQCTGIEIDDRRIRVDYS  157
PpeTRA-2A     NDIFSKFGRIKQIQIIMDAKTGRSRGFCFVYFADADDAKVARENCSGMEIDGRRIRVDYS  171
PbeTRA-2      YQIFSKFGTIHRTQIIMDAMTGCSRGFCFVYFANADDAKVAKDNCSGMELDGRRIRVDYS  178
PpaTRA-2      YQIFSKFGTIHRTQIIMDAMTGCSRGFCFVYFANADDAKVAKDNCSGMELDGRRIRVDYS  178
PduTRA-2      YQIFSKFGTIHRTQIIMDAMTGCSRGFCFVYFANADDAKVAKDNCSGMELDGRRIRVDYS  178
               :**:*** *.: ***:*: ** ***:*****  :::*****:::*:*:*:*.********
                                       RNP-2

              Linker Region         RS domain 2
LumTRA-2      LTARPHTPTPGIYMGKASQRCGD---SSTSRSNRYEDTS-YEKTRNRSPSPYKRR-RRSR  212
LneTRA-2      LTARPHTPTPGVYMGKASQRCGDRGDSSTSRSNRYEDSS-YEKTRNRSPSPYKRR-RRSR  215
PpeTRA-2A     ITERAHTPTPGKYMGVSTGRGREHNRSNRDDDYYYERESRSRYIERRSPSPYRRKRYRSR  231
PbeTRA-2      ITQRPHTPTPGVYMGQPTGRGRENRDRSRRN-----------------------------  209
PpaTRA-2      ITQRPHTPTPGVYMGQPTGRGRENRDRSRRN-----------------------------  209
PduTRA-2      ITQRPHTPTPGVYMGQPTGRGRENRDRSRRN-----------------------------  209
              :* * ****** ***  : *  :    .

LumTRA-2      SRSHSPRVRRYH        224
LneTRA-2      SRSYSPRVRRYH        227
PpeTRA-2A     SRSYSPRIRRYR        243
PbeTRA-2      -----------?        209
PpaTRA-2      -----------?        209
PduTRA-2      -----------?        209
```

**Figure S22. Multiple sequence alignment of sand fly TRA-2 proteins.** Sequence alignment of TRA-2 proteins of *Phlebotomus perniciosus* (PpeTRA2-A), *Phlebotomus papatasi* (PpaTRA2), *Phlebotomus bergeroti* (PbeTRA2), *Phlebotomus duboscqi* (PduTRA2), *Lutzomyia umbratilis* (LumTRA2-A) and *Lutzomyia neivai* (LneTRA2). The RNA recognition motif (RRM) is highlighted in black, RS domains are in light grey, and the linker region is in dark grey. Question marks indicate uncompleted sequences at N- or C- termini. Bold letters indicate amino acids identical between *P. perniciosus*, *L. umbratilis* and *L. neivai*. RNP-1 and RNP-2 indicate the positions of two ribonucleoprotein identifier sequences, which are highly conserved between RRM proteins. Note the high number of identical amino acids within the RRM domain, whereas both RS domains are less conserved but are abundant in arginine and serine. Gaps were introduced in the alignment to maximize similarity. The protein sequences alignment was performed using the Clustal-Omega software (1.2.4).
