## Additional_file_9_Figures_S23-S28 for "Genomics and transcriptomics to unravel sex determination pathway and its evolution in sand flies"

#### *P. papatasi dsx* gene model

```
Scaffold7_Length=1359615

                                                                    994630      EXON1
                                                                    |
DNA:  ctaaattctctgtatatctagactaagtgtcagtcttcggttaaaagttcaattaagtgggacgctgttgATGGTGTCGGAGCACAATACGTGGAGCGACGTGATGTCCAATTCGGACCTCACAGACTCGAAAAACGAGTTGTGCGGAGGAGCCTCGAGCTCCAGCGGCAACTCATTGAGCCCGAGGACGCCACCAAATTGCGCCCGCTGCCGCAATCACGGCCTCAAGATAGCCCTTAA
+2fr:                                                                       ·M··V··S··E··H··N··T··W··S··D··V··M··S··N··S··D··L··T··D··S··K··N··E··L··C··G··G··A··S··S··S··S··G··N··S··L··S··P··R··T··P··P··N··C··A··R··C··R··N··H··G··L··K··I··A··L··K

DNA:  GGGTCACAAACGCTACTGCAAGTACCGTTACTGCAACTGCGAAAAGTGTCGTCTAACGGCCGAACGTCAGCGGGTGATGGCCTTGCAGACGGCCCTAAGACGTGCTCAGGCTCAGGACGAGGCACGTCAGATCTCCGCGGATGAGGTGCCACCGACGCCACCGCCAGCCCCGTCCATTTGCACCCTTGCCACTCCTAAGAGTCAGTGCGATAGCGAACCACCGGCACCCATTACGGTGCC
+2fr: ··G··H··K··R··Y··C··K··Y··R··Y··C··N··C··E··K··C··R··L··T··A··E··R··Q··R··V··M··A··L··Q··T··A··L··R··R··A··Q··A··Q··D··E··A··R··Q··I··S··A··D··E··V··P··P··T··P··P··P··A··P··S··I··C··T··L··A··T··P··K··S··Q··C··D··S··E··P··P··A··P··I··T··V··P

                                                                                                                          995176
                                                                                                                          |
                                                                                                                0.91 actgcaaGTaagtac
DNA:  CACACCGGCACGTTCCCTAGAGGGTTCCTGCCACTCATCATCGGCCACACCCTGTTCTGCTTCCGGAGAACCCATCTCCGTGCCATCCAGCCGCAAATTGGCCCCAGTCAATCCAACCATCAGAAATCCAACTGCAAgtaagtaccattcaaattatccttactacgtatatggtaccctctacaacagctaccctacttatggccacatacaccaaaaaaaaatataagcttaatcct
+2fr: ··T··P··A··R··S··L··E··G··S··C··H··S··S··S··A··T··P··C··S··A··S··G··E··P··I··S··V··P··S··S··R··K··L··A··P··V··N··P··T··I··R··N··P··T··A··

                                                   ----////----

                                      1114154   EXON2                                                                                    1114294
                                      |                                                                                                  |
                      0.90 ttcctcttgtcacgctttcAGgtcaaagctccgaagtcttc                                                              0.90 gaagaagGTgagatg
DNA:  cttgattttaatgttcaactcttcctcttgtcacgctttcagGTCAAAGCTCCGAAGTCTTCTTTGACAATTGCCAGAAATTGCTGGACAAATTCAATTATCCTTGGGAACTGATGCCCCTAATGTATGTGATCTTGGAAAATGCCCCGGACGTGAGGGAAGCGGCACGGAGAATTGAAGAAGgtgagatggtattttattgatttttatagaaattgaaaaatatctttttgggtttac
+3fr:                                         S··Q··S··S··E··V··F··F··D··N··C··Q··K··L··L··D··K··F··N··Y··P··W··E··L··M··P··L··M··Y··V··I··L··E··N··A··P··D··V··R··E··A··A··R··R··I··E··E··

                                                   ----////----

                                                        1118126     EXON F
                                                        |
                                      0.76 atttaccacttcacgatctAGgtcaatgcgttgtgaatgag
DNA:  gcatttgtgtgaggattaaactttgacaaatgattgctaaacaacatttaccacttcacgatctagGTCAATGCGTTGTGAATGAGTATACTCGAATGCATAATCTCAATATGTATGATGGAGCCGAATTGAGGGGCTCCACCCGGCAATGCGGATGATAAtttactgccagcagtttcatatgttagtaaaaaaaaaacaaacgtTCATCAATCAACAttgagataatatgcaaatgtt
+3fr:                                                               G··Q··C··V··V··N··E··Y··T··R··M··H··N··L··N··M··Y··D··G··A··E··L··R··G··S··T··R··Q··C··G··*··*·

DNA:  ggcactcgtacacacctaagttcaaactaagatatatcgtaacttaacccaagaaatgcgtatttgaaagaaagatcaaccccgttaagcttacacaaaactcttttattgattagatttactaattaattgtagatgtacataatatgagtatggccacttcttttcattctttttatatatttagtaattaatatcgtttaaaacttcttggagtgcgcatttttaattgcttaaaaa

DNA:  gaatatttgtggataattattattttaaatgattatttatgattgttcgcggtagtggctattggtgttaaagtgatTCTTCAATCAACAtttTCTTCAATCAACAttTCTTCAATCAACACCTTCAATCAACAaatctgcaatggaaacaaaaaaatcaagctcaaaaatcacaaaacatatcttccataatttttgactcaattataatgatgtttttttggcgggaaatgccaaaag

                                                                              1118868
                                                                              |
                                                                    0.70 tcacaatgtgagaaa
DNA:  gacccacttaaatcagactctcaagagagaagaaaatcaatcaaccatttggatttagccacaatcttaacagtacatccattcacaatgtgagaaatgggcagcgtttcatcaaattgacaacaatatagatgtggatgccagaagattgcaacagcgttaaaaaaagtggaacaacagcaaaaaaagtgcaaggagaaaacacattaatgttcaaaggcgaatggtaagaagttgatgt

                                                   ----////----

                                                                                  1141155      EXON M
                                                                                  |
                                                            0.57 atcaggatgtttattttgcagctgcgagtataataaaatgt
DNA:  tatcggccactgaagagcccgcgagtcattggttaatcatgattaccggatattacaatattattagttaagtaatcaggatgtttattttgcagCTGCGAGTATAATAAAATGTCGATATGGACATGGACTCCCAATCCCAGCACACCAGTACTACACATACCTATCTACGGCGGCAGCGAGAGTGCAGACGTCGATATCCATATCCTTCGTACTACTACACATACTGGGCAGCCGGTA
+2fr:  ·                                                                                                A··A··S··I··I··K··C··R··Y··G··H··G··L··P··I··P··A··H··Q··Y··Y··T··Y··L··S··T··A··A··A··E··C··R··R··R··Y··P··Y··P··S··Y··Y··Y··T··Y··W··A··A··G··S

                                                                                                                                                                       1141498
                                                                                                                                                                       |
DNA:  TACAAGTCCCTTCCTATATAACTTCTATAACGAGACAGTACTGTCGCCGCATTTGCACAGCAAGTACCTCAATTCGGAAAGTACTCCTGCGGTACGGAATATATCGGAATCTCCGTCAGCCTATTCATCTCTGCCACTTCCTGCCGCTCTACAAATTACTCGTCTGATACCAACGAATAGCTTATCTGGTGCATCGACGTAGcgatgtatactctatgtgattcaactaatagtgattgt
+2fr: ··T··S··P··F··L··Y··N··F··Y··N··E··T··V··L··S··P··H··L··H··S··K··Y··L··N··S··E··S··T··P··A··V··R··N··I··S··E··S··P··S··A··Y··S··S··L··P··L··P··A··A··L··Q··I··T··R··L··I··P··T··N··S··L··S··G··A··S··T··*·
```

**Figure S23. Manually-curated *P. papatasi doublesex* gene model.** Exonic sequences are indicated by black cases; black upper cases indicate coding sequences. Intronic sequences are indicated by gray lower cases. Start and stop codons are highlighted in green and red, respectively. In yellow boxes, putative TRA/TRA-2 binding sites. Exon start/end positions in the scaffold/s are indicated. Intron boundaries were predicted using transcripts vs genome alignments and confirmed by *de novo* prediction with Berkeley BDGP Splice Site Prediction Tool with default parameters (http://www.fruitfly.org/seq_tools/splice.html); prediction scores are indicated in red.

### *L. longipalpis dsx* gene model

```
Scaffold136_Length=197883

                                                                                 60489      EXON1
                                                                                 |
DNA:  caacaactaaaaaagaacgaaacttggctaaattcactccaagtggcacgctgttgatggtgtcggagcacaatacgtggagcgacgtgATGTCCAATTCGGACCTCACAGACTCGAAGAATGAGTTGTGCGGGGGAGCCTCGAGCTCCAGCGGCAACTCATTGAGCCCGAGGACGCCGCCAAATTGCGCCCGCTGCCGCAATCACGGCCTCAAGATTGCCCTAAAGGGTCACAAGCGTT
+3fr:    ·                                                                                       ·M··S··N··S··D··L··T··D··S··K··N··E··L··C··G··G··A··S··S··S··S··G··N··S··L··S··P··R··T··P··P··N··C··A··R··C··R··N··H··G··L··K··I··A··L··K··G··H··K··R··

DNA:  ACTGCAAATTTCGCTACTGCAACTGCGAAAAGTGCCGCCTCACGGCCGAACGACAGCGGGTGATGGCCCTACAGACGGCCCTAAGACGTGCTCAGGCTCAGGACGAGGCGCGTCAAATCTCCGCGGATGAGGTGCCACCGACACCACCGCCACTTACTGGTCAAATTGCAACAACGCCCAAAAGTCAGTGCGACGGTGAACTACCGCGATCGAATACAGTGCCAACTCCGGCACGTTCCC
+3fr: Y··C··K··F··R··Y··C··N··C··E··K··C··R··L··T··A··E··R··Q··R··V··M··A··L··Q··T··A··L··R··R··A··Q··A··Q··D··E··A··R··Q··I··S··A··D··E··V··P··P··T··P··P··P··L··T··G··Q··I··A··T··T··P··K··S··Q··C··D··G··E··L··P··R··S··N··T··V··P··T··P··A··R··S··
·

                                                                                                    60999
                                                                                                    |
                                                                                  0.94 gcagcaaGTaagttt
DNA:  TCGAGGGCTCCTGCCACTCATCTTCGGCAACACCATGCTCCGCTTCTGGTGAACCCATCACCGTGCCATTAAGCCGTAAGCCTCCAGCAGTTAATCCAACCGTTAGAAGCCCAGCAGCAAgtaagtttttctttatcttttctacacacttacatacctatataatattaaatatatatatttcttatctctatttgccatctcagaagatttattctctttattgtcacaatttcctt
a
+3fr: L··E··G··S··C··H··S··S··S··A··T··P··C··S··A··S··G··E··P··I··T··V··P··L··S··R··K··P··P··A··V··N··P··T··V··R··S··P··A··A··


Scaffold377_Length=75672

                                                                      40065      EXON2                                                                                                                    39922
                                                                      |                                                                                                                               |
                                                 0.97 tttcttttcatgtactttcAGgtcaaagctccgaggtgttc                                                                                              0.24 gaggaagGTttgtcg
DNA:  gcaaattgttaatgtgccatttgctttctttaatggtcacctccttattctttctttcttttcatgtactttcagGTCAAAGCTCCGAGGTGTTCTTTGACAATTGTCAGAAGTTACTGGATAAATTCAATTACCCATGGGAGTTGATGCCCCTAATGTATGTGATCTTAGAAAATGCCAACGTAGATATGCAGGAAGCAGCACGAAGGATTGAGGAAGgtttgtcgtgtcaatctcata
-3fr:                                                                         S··Q··S··S··E··V··F··F··D··N··C··Q··K··L··L··D··K··F··N··Y··P··W··E··L··M··P··L··M··Y··V··I··L··E··N··A··N··V··D··M··Q··E··A··A··R··R··I··E··E··

----////----

          37672      EXON F
          |
 0.51 aaatttAGgtcaatgtgttgtgaatgaa
DNA:  aaatttagGTCAATGTGTTGTGAATGAATACACCCGAATGCACAATCTCAATATGTACGATGGAGCCGAACTGCGGGGTTCAACGAGACAATGTGGATGATAAtacatttcgaattatttcacgtgttagaactttaattaaaaataaaaaaaaaacaaacaaagagataatttgcaagtgaatttgaatacttgtcctcacacctaagttcaaattacaaagtaataacttatccaaga
-2fr:         ·G··Q··C··V··V··N··E··Y··T··R··M··H··N··L··N··M··Y··D··G··A··E··L··R··G··S··T··R··Q··C··G··*··*·

DNA:  acaatttattttttatttttttattattttgtaaaacaacaattaaagccctaagctttgaaattcaaactcaaaaaaactgtttttttatatataattttctttgaattattaaaatgtgtgtagtgtaaataaagcctttaaatttctactaagaaattcatcgtaaataattaatttttttttaatgtgtgacattaaaaagtgttcaagtgtaaggcattttggtttgctgaaaagaaa

                                                                                                                                                                 37014
                                                                                                                                                                 |
                                                                                                                                                      0.74 atcaactGTgagtta
DNA:  atcttttatggataattattgttttaaatgattatttatgattggatttgcattcgcgtgaaagaaaaaaaaaacgttggtgttaaaatgatTCTTCAATCGAAAtggattttcattgtcttcaacacggTCTTCAATCAGCAttTCATCAATCAACAcTCTTCAATCAAGAttgTCTTCAATCAACTgtgagttaaatcgattcttttacaaaattccagcatcgcagccttgctcactc

DNA:  tttcacattaaaaatttggttttaattctgaaaaaccaaaaaggaacttaagaggggcaaaaaagttgctcaattaaaccattgagcggaagtaatttcataaataaaatcaatttctaacaataaaatacgggatgtattaaaaaaaacaacaaccagcaatatttaatcaaaatggcaacaagaagttgctggcaaggttgataccaatatatgtatataaaaggttgtgtgtggatg

----////----

                                                  19946      EXON M
                                                  |
                              0.89 gtaagtaaatatttttttacAGcaatgggagagtgcagacgt
DNA:  atttttttccatttaattcatgctaaaattatgtaagtaaatatttttttacagCAATGGGAGAGTGCAGACGTCGATATCCATATCCTTCGTACTACTACACATACTGGGCAGCCGGTAGTAGTCCCTACCTGTTCAATTACAACATCCTCTCGTCGCAATTGCAAAATAAGAGCTTTAGCTGTGACAGTATTTCATCAACACGTGGCATCTCAGAGGCACCATCGTCCTACCCCTCAA
-3fr:                                                   A··M··G··E··C··R··R··R··Y··P··Y··P··S··Y··Y··Y··T··Y··W··A··A··G··S··S··P··Y··L··F··N··Y··N··I··L··S··S··Q··L··Q··N··K··S··F··S··C··D··S··I··S··S··T··R··G··I··S··E··A··P··S··S··Y··P··S··

                                                                 19690
                                                                 |
DNA:  TATCACTACCTGCCGCCGCCCTACAAATCAGACGTCTGATACCAACTGGTAACCTAGCCGGTGCATCGACGTAGcgatgtgttttctatgtgattcaactagtagttatagtaagtacag
-3fr: I··S··L··P··A··A··A··L··Q··I··R··R··L··I··P··T··G··N··L··A··G··A··S··T··*·
```

**Figure S24. Manually-curated *L. longipalpis doublesex* gene model.** Exonic sequences are indicated by black cases; black upper cases indicate coding sequences. Intronic sequences are indicated by gray lower cases. Start and stop codons are highlighted in green and red, respectively. In yellow boxes, putative TRA/TRA-2 binding sites. Exon start/end positions in the scaffolds are indicated. Intron boundaries were predicted using transcripts vs genome alignments and confirmed by *de novo* prediction with Berkeley BDGP Splice Site Prediction Tool with default parameters (http://www.fruitfly.org/seq_tools/splice.html); prediction scores are indicated in red.

#### *P. papatasi fru* gene model

### A)

```
Scaffold43920_Length=20651

                                                                                                                                                        1050  EXON S2
       827      EXON S1                                                                                                                                  |
       |                                                                                                                                      0.95 cacctagGTaggttc
DNA:  gcatagaATGATGACTACTCCGGACATCTTCGGCAGCCCCTTTCCGCCGTATAGGTGTCAGCCGTTGGCTGTACTTCCACCACGTGATGATAGTCCGCCAATGCCGGCACTTGACCTCAAACGCTATCCCAGCACTGAGGTTCCTCCGGCATCAACAACACCTGTCCGTTTACCTGAACGTGAAATGCCGATGCCGCACCTGCGTCCATCTTCGGCACAAACACCACCTAGgtaggttct
+2fr:        ·M··M··T··T··P··D··I··F··G··S··P··F··P··P··Y··R··C··Q··P··L··A··V··L··P··P··R··D··D··S··P··P··M··P··A··L··D··L··K··R··Y··P··S··T··E··V··P··P··A··S··T··T··P··V··R··L··P··E··R··E··M··P··M··P··H··L··R··P··S··S··A··Q··T··P··P··R··*·

DNA:  acctcctgggcgctagccaaattagcaggacacagtgcgacagaccacatctgcatctctcaggagaaaagTCTGCAATCAACTtgtctcgctgtgtcttatgtacagaatggcatggggaagtagtgcattgatcaattccctgtagaacaaaaataaaactataccactgctttaaggcttttcatttcttcgggtgcactgttcaccttttgtactcataaagatttcagtaactct

                                                                                                                                                   ??? 0.76 caaggtgG
DNA:  cctagaaaatcatcgagatttaattcaattatgcttttgcagtaaatgcttctgcagagaaatttagaagaacatgatgaaaatcggaagttaaaatatgaggatctcaagatcacagtcttgatgaaagattcactcgcgagggaatctgacgaagattcacattggaaaatgatattaaatgaatacgagatggtgatgtattattcagtggcatcatgaagtacaagtgcaaggtgg

      Tgaggct ???
DNA:  tgaggctcaaaaacgttgaaacatacgggggtgttgtacgtttgataccgtccaaaattacactcaagcccctattgtatgattcttgggcgcaactcttggcattttacatccatctcaagagagtgcttttatgcttggcggcgattttcccttcctccacctccatgaaatgttcaaaattgtttcgtgaaaattctgcaggaagctgcatttcaatatatataaacctttggccat

DNA:  tttctggagaatccttttgatatacagagaccgtgaggaccttgggcatcaggggtgtgcttgcatcttgtacaagtcaaacatgccccaagattgaaaagaattcctttttatgggaagtcaccggaatacagatacaatgtactacttttcagaaaagtggaaaatgccttacttctaacaatgcacttaccaaattccatagtgtgtttctttcgaggcacgtggccgttattgatg

DNA:  aatgttgattgtagatttgtttttgttcaaatgcatgttaatcgtggcaaatgttgattgaaaatataatgattctttacgactattgcaataaataggtattactggcaaaaaagcatttctatttgaatttcttcaagtcatttcttgtttatctaattcgactttcattagcaaacttttctactcttctactattatatttttgctcctgttgctctacatttcttccttattctg

DNA:  gcggtagaaatttttctatattgcatacaaaagttactatcggagcaatatattgaaaactccgcgattgcaattttttttttggttactttttttgcaaaggaaaaatctttgtagattttccatactcggaaacttctttattttgtactgtatcacttgctcatatcctggatctcgtatttcttttaaacagtgagaatgacgcatgcttataatttgaaagaaaccttctggt

DNA:  gttagatttatgtttaagaaatttgtatgttaggttgttgctcacgcagggatggcggattggatcagggttttcacagtggatgaggaaataaggcaggggagtatatgtaatctcttaatttcaaagtaacaccttggtccaatcacaccagaattttatttctgcttccacattttcatcactggcctttgaacatcttttttttgagtgaaaacaaaaatattcaaaaaatgaata

                  ??? 0.75 cattttgGTcagtaa ???
DNA:  tacattttggacagtaaattattccattttggtcagtaaataacgtaacattaacgacaaattgctattaactttaaacattcctgagatatcaaaatatgatttataacttctctatttcccattgccaagaatattttattaattctatgcattgagaatgtgtgaatcccattttaattacttgccgaaattattttttatattttactttaaccctttaacgacgagacagt

DNA:  ttttgcggaccgaaaatcagcaataaaaatgaaaccgataaacaaaattagtaaaaagtcttagataccttagaaaatctatcagaatctgattcgaagtatttttgccctgtatgagcgatagggacaaaaatagccaaaaaaatttagtaaattttctgacaataccaatgacagataattttcactctaacgaaaaatattatgtttgagtattgtagtttctttttcctaaacggc
```

**Figure S25A. Manually-curated *P. papatasi fruitless* gene model**. The genomic region regulated by sex-specific alternative splicing, which includes the two sex-specific exons S1 and S2, is reported. Exonic sequences are indicated by black cases. Black upper cases indicate coding sequences. Intronic sequences are indicated by gray lower cases. Start codons are highlighted in green. In yellow boxes, putative TRA/TRA-2 binding sites. Exon start/end positions in the scaffolds are indicated. Intron boundaries were predicted using transcripts vs genome alignments and confirmed by *de novo* prediction with Berkeley BDGP Splice Site Prediction Tool with default parameters (http://www.fruitfly.org/seq_tools/splice.html); prediction scores are indicated in red. Red question marks indicate putative predicted female-specific donor splice sites. The absence of the expected TRA/TRA-2 binding site cluster in the proximity of female-specific donor splicing site in Scaffold 43920 indicates a possible incorrect assembly of this genomic region.

#### *P. papatasi fru* gene model

**B)**

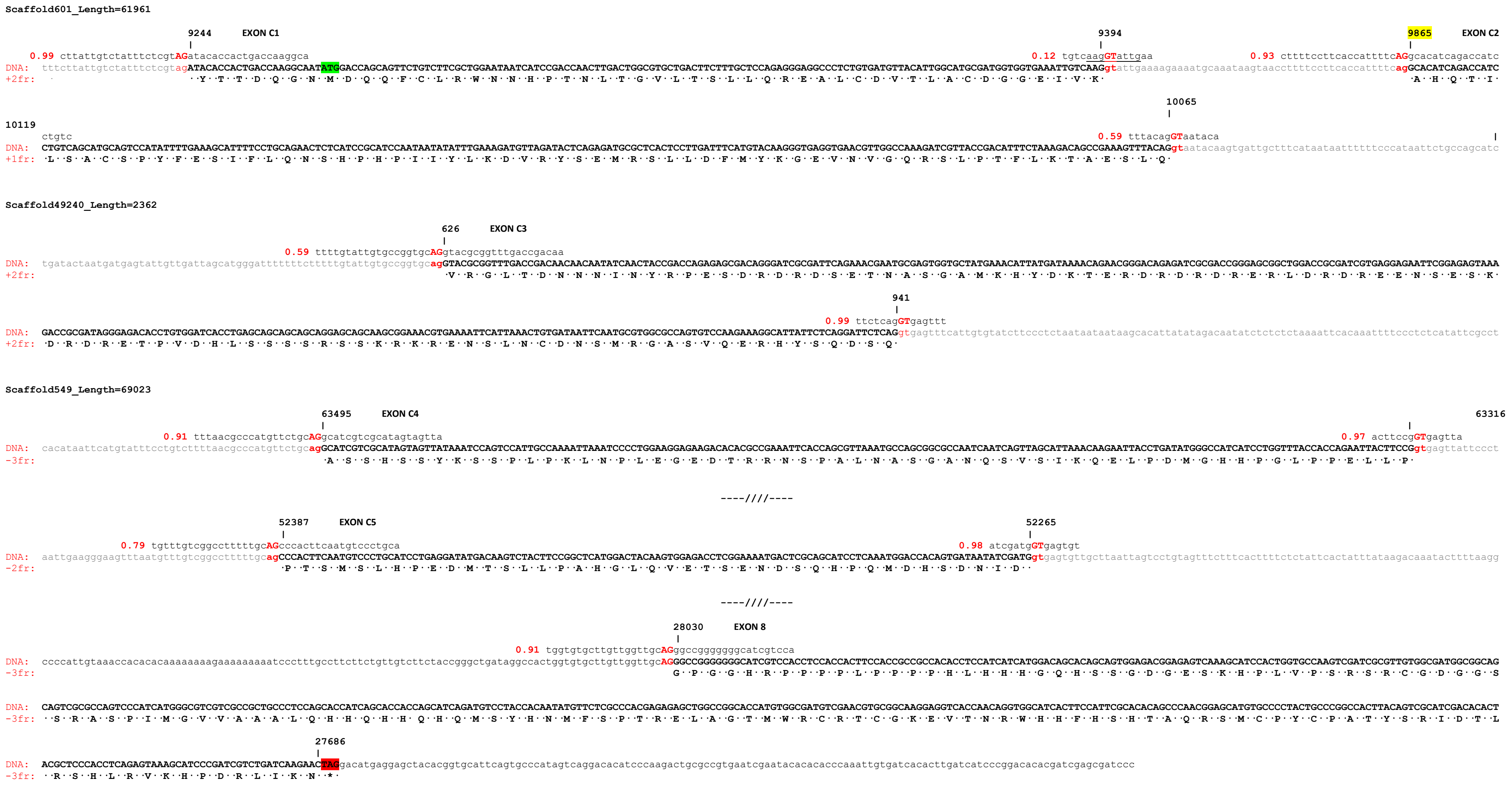


**Figure S25B. Manually-curated *P. papatasi fruitless* gene model**. The genomic region which includes common exons 1-5 and the terminal exon, encoding for the Zinc Finger domain of type C, is reported. Exonic sequences are indicated by black cases. Black upper cases indicate coding sequences. Intronic sequences are indicated by gray lower cases. Start and stop codons are highlighted in green and red, respectively. In yellow boxes, putative TRA/TRA-2 binding sites. Exon start/end positions in the scaffolds are indicated. Intron boundaries were predicted using transcripts vs genome alignments and confirmed by *de novo* prediction with Berkeley BDGP Splice Site Prediction Tool with default parameters (http://www.fruitfly.org/seq_tools/splice.html); prediction scores are indicated in red. Red question marks indicate putative predicted female-specific donor splice sites. The absence of the expected TRA/TRA-2 binding site cluster in the proximity of female-specific donor splicing site in Scaffold 43920 indicates a possible incorrect assembly of this genomic region.

#### *L. longipalpis fru* partial gene model

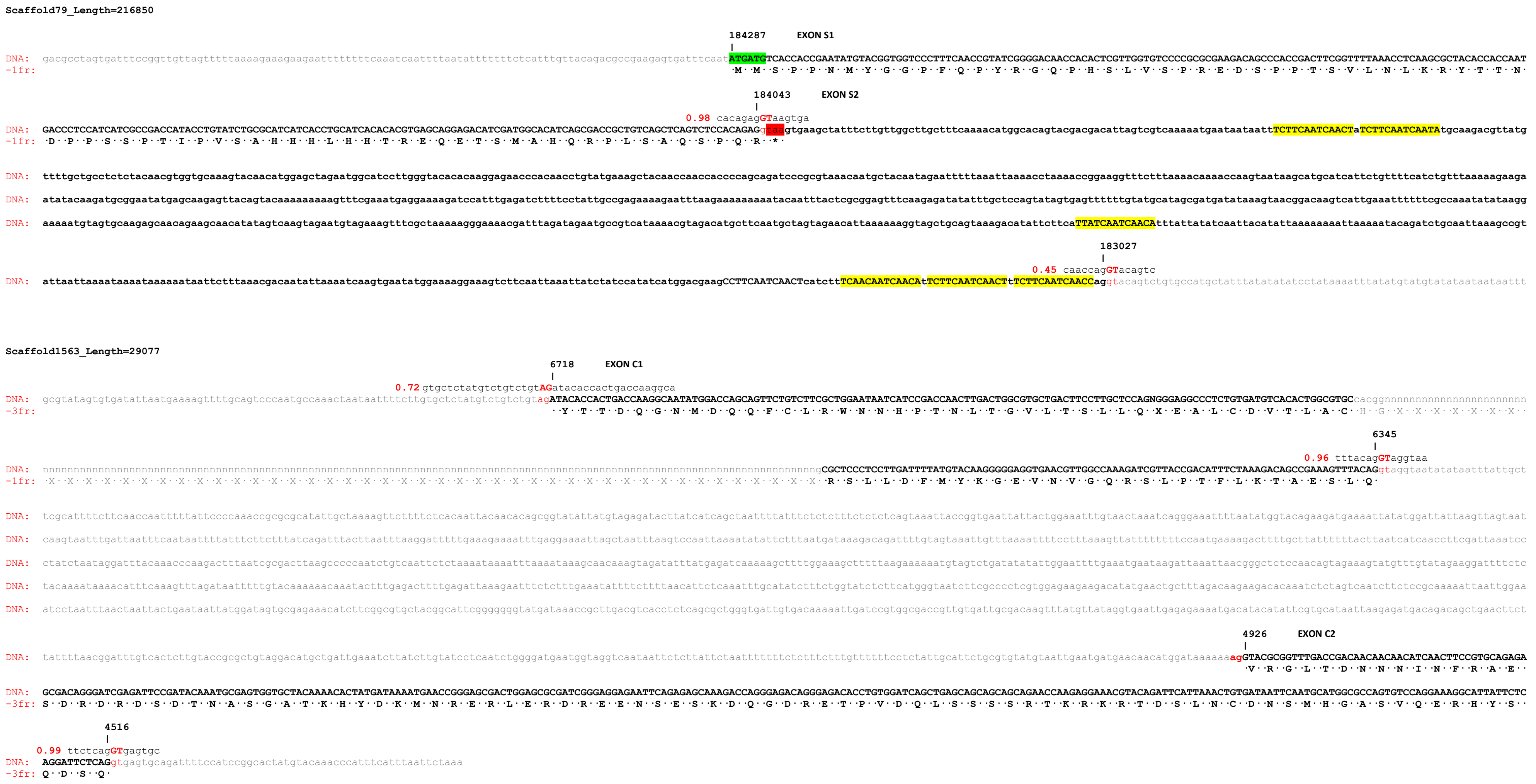


**Figure S26. Manually-curated *L. longipalpis fruitless* partial gene model**. The genomic region regulated by sex-specific alternative splicing is included. Exonic sequences are indicated by black cases. Black upper cases indicate coding sequences. Intronic sequences are indicated by gray lower cases. Start and stop codons are highlighted in green and red, respectively. In yellow boxes, putative TRA/TRA-2 binding sites. Exon start/end positions in the scaffolds are indicated. Intron boundaries were predicted using transcripts vs genome alignment and confirmed by *de novo* prediction with Berkeley BDGP Splice Site Prediction Tool with default parameters (http://www.fruitfly.org/seq_tools/splice.html); prediction scores are indicated in red.

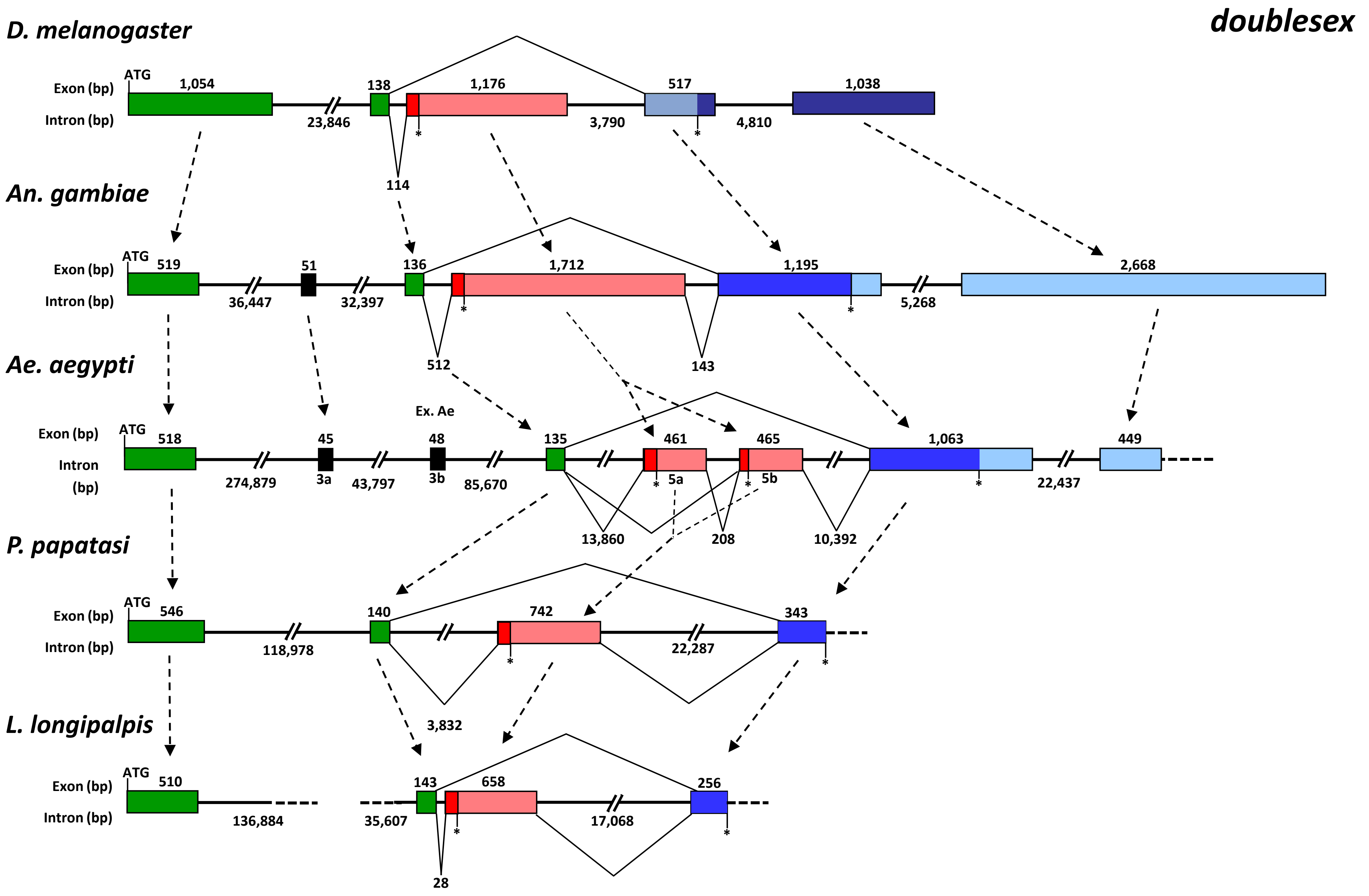


**Figure S27. Comparison of genomic structures of diptera *dsx* genes.** Green boxes represent the OD1 and OD2 domains encoding common exons. Black boxes represent exons encoding protein regions conserved in mosquitoes but not in fruit flies and sand flies. Alternative male-specific and female-specific exons are represented as blue boxes and pink boxes, respectively. Asterisks indicate the position of stop codons. In *Drosophila*, the *dsx* gene encompasses a 45 Kb-long region on chromosome 3R and is organized into 6 exons and 5 introns, with three common exons followed by a female-specific and two male-specific exons. DmDSXF translation initiates at the ATG within exon 2 and terminates within the female-specific exon 4, while in the case of DmDSXM, translation begins at the same ATG and terminates within the first male-specific exon 5.

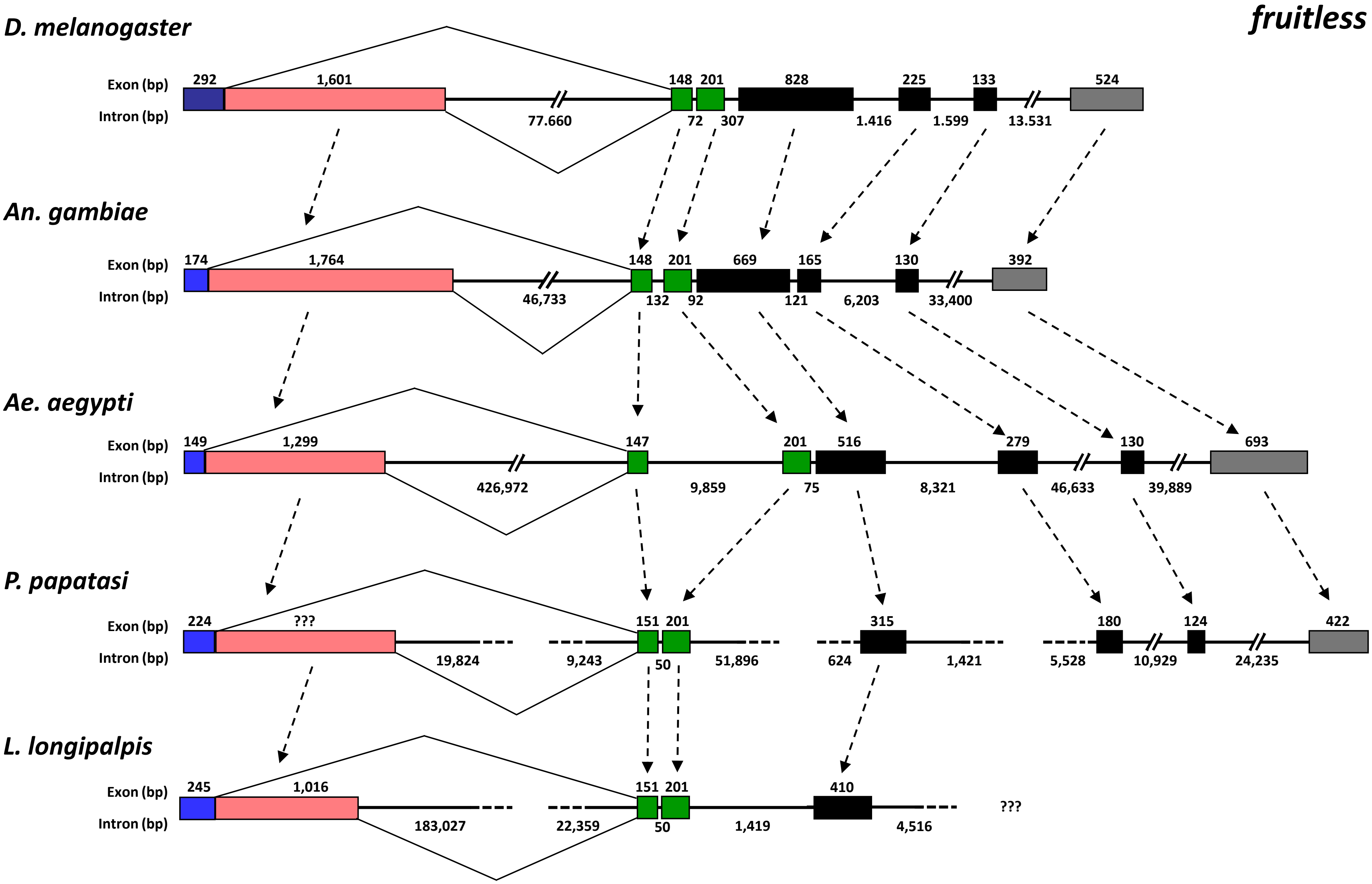


**Figure S28. Comparison of genomic structures of diptera *fru* genes.** Fru S1 common (but encoding the male-specific N-terminus) and S2 female-specific exons are represented as blue and pink boxes, respectively. Green boxes represent the non-sex-specific C1 and C2 exons, encoding the BTB domain. Black boxes represent the non-sex-specific C3-C5 exons encoding the Connector region of FRU proteins. The terminal grey boxes represent the ZnF-C domain encoding exons. The *Drosophila fru* gene encompasses a 98 Kb-long region and is organized in 7 exons and 6 introns with 6 common exons preceded by the sex-specific regulated region, with a common exon and a female-specific exon.
