## Additional_file_10_Figure_S29 for "Genomics and transcriptomics to unravel sex determination pathway and its evolution in sand flies"

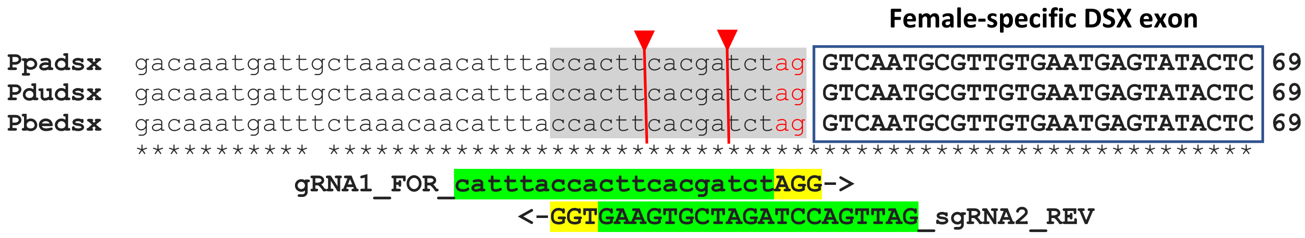


**Figure S29. Crispr/Cas9 target sites in sand fly *dsx* genes.** ClustalW multiple alignment of the genomic region surrounding the 3' acceptor female-specific splicing site (highlighted in light grey) of *dsx* gene in *P. papatasi* (*Ppadsx*), *P. duboscqi* (*Pdudsx*) and *P. bergeroti* (*Pbedsx*). Last intronic AG di-nucleotides are indicated in red. Exonic sequences are indicated in upper cases. As observed for *Anopheles* mosquito species (Kyrou et al., 2018) this genomic region is highly conserved also in *Phlebotomus* sand fly species, most probably because of the critical sex-specific splicing regulation required for the *dsx* gene. Two putative sgRNA target sites are indicated, with PAM NGG sequence highlighted in yellow. sgRNA target sites were predicted using the CHOPCHOP tool with default parameters (http://chopchop.cbu.uib.no/). The two putative Cas9 cutting sites are located into the polypyrimidine tract of the 3' acceptor female-specific splicing site and could led to the disruption of the correct splicing mechanisms in female individuals.
