## Additional_file_11_Figures S30-S33 for "Genomics and transcriptomics to unravel sex determination pathway and its evolution in sand flies"

### *P. bergeroti fru* partial gene model

**Contig_256375_Lenght=2168**

```
                                                                                                                                                   1948
DNA:  gctgaacactaacaatagttcttcacaagaaaaaattaccaaataaattatcaaattgtaaatttctgtatttctttaaaagatttgaactgtataaaagatactaatttttttgaaaaatagcagtaaaatatcatcatacaattatatttttaactaaaatttgcatgtgctatctctctatttgcagagcttgagcagagtaagtgatcgcatagaATGATGACTACTCCGGACA
-2fr:                                                                                                                                              ·M··M··T··T··P··D··

                                                                                                                        1724
                                                                                                          0.96 cacctagGTaggttc
DNA:  TCTTCGGCAGCCCGTTTCCGCCGTATAGGTGTCAGCCGTTGCCTGTACTTCCACCACGTGATGATAGTCCGCCAATGCCGGCGCTTGACCTTAAACGCTATCCCAGCACTGAGGTCCCACCGGCATCAACAACACCTGTCCGTTTATCTGAACGTGAAATGCCGGTGCCGCACCTGCGTCCATCTTCAGCACAAACACCACCTAGgtaggttctacctcctgggcgctagccaaattag
-2fr: I··F··G··S··P··F··P··P··Y··R··C··Q··P··L··P··V··L··P··P··R··D··D··S··P··P··M··P··A··L··D··L··K··R··Y··P··S··T··E··V··P··P··A··S··T··T··P··V··R··L··S··E··R··E··M··P··V··P··H··L··R··P··S··S··A··Q··T··P··P··R··*·

DNA:  caggacacagtgcgacagatcacatctgcatctctcaggagaaaagTCTGCAATCAACTtgtctcgctgtgtcttatgtacagaatggtatggggaagtagtgcgctcccctgccgtatttcctcatccactgtgaaaaccctgatccaacccgccatccctgcgtgagcaacaacctaacatacaaatttcttaaacataaatctaacaccagaaggtttctttcaaattataagcat

DNA:  gcgtcattctcactgtttaaaagaaatacgagatccaggatatgagcaagtgatacagtacaaaaataaagaagtttccgagtatggaaaatctagaaagattttccttagcaaaaaaaagtaaccaaaaaaaaaaatttgcaatcgcggagttttcaatatattgctccgatagtaacttttgtatgcaatatagaaaaatttctaccgccagaataaggaagaaatgtagagcaaca

DNA:  ggagcaaaaatataatagtagaagagtagaaaagtttgctaaagaaagtcgaattagataaacaagaaatgacttgaagaaattcaaatagaaatgctttttgccagtaataactatttattgcaatagtcgtaaagaatcattataTTTTCAATCAACAtttgccacaattaacatgcatttgaacaaaaatataTCTACAATCAACATtcatcaataacggccacgtgcctcgaaa

                   953
                   |
          1.00 gaatttgGTaagtgc
DNA:  gaaacacactatggaatttggtaagtgcattgttagaagtaaggcattttccacttttctgaaaagtagtacattatatctgtattccggtgacttcccataaaaaggaattcttttcaatcttcgggcatgtttgacttgtacaagatgcaagcacacccctgatgcccaaggtcctcacagtccctgtatatcaaaaggattctccagaaaatggccaaaggtttatatatattgaa

DNA:  atgcagctttctgtagaattttcacgaaacaattttgaacattcatggggggaggaggaagggaaaatcgccgccaagcataaaagcactctcttgagatggatgtaaaatgccaagagttgcgcccaagaatcatacaataggggcttgagtgtaattttggacggtatcaaacgtacaacacccctgtatgtttcaacgtttttgagcctcaccaccttgcacttgtacttcatgatg

DNA:  ccactgaataatacatcaccatctcgtattcatttaatatcattttccaatgtgaatcttcgtcagattccctcgcgagtgaatctttcatcaagactgtgatcttgagatcctcctattttaacttccgatttcatcatgttcttctaaatttctctgtagaagcatttactgctaaagcataattgaattaaatctcgatgattttctagcagagttatactgaaatttttatgagt

DNA:  acaaaaggtgaacagttcacccaaagaaatgaaaagtcttaaagcagtggtagttttatttttgtgctacagggaattgatcagtttatgtaatctcttaatttcaaattaacacatttgtcattcgtcacacactagaattttatttctgcttccacatattaatcactgtctttgaatgaaagactgcctttgagtgaaaataaaaatatgcagttttggtcagtaggggaaagtac

DNA:  tctctctccgaacgttc
```

**Contig_16679_Lenght=1684**

```
DNA:  tatatatatatatatatatatatatgttctatgccttatataatatatactctatatacattatatagtaaaatcaataatttttaatacattcgatcttttacaaatctttttaaataaaattaaaagaggctttgacacatctattaaacaaaatacttttattcatatttttctattcatttgggagactttttttgagacaatcttcctgggtcaccctgttcataaactttg

DNA:  agtgttaatgaaggctttttttgcctcttccatgtgtgttctccgacgcgaatagaaaaagacagatatgacagtatgttgcaaaattttagaataaatttattttctagatgctcatttttcgcaaacacctcgcccatgtataagcgccaaaaggtttttgcaggtggaactggacatggtcaataaaaaattagtctttctaccttgatacaatccgtgtgcaaactttccatt

                                                                                  1105
                                                                                  |
                                                          0.99 cttattgtctatttctcgtAGatacaccactgaccaaggca
DNA:  aggttttttttttggtaaaatagtcgaggagtgatattaatgaaaagttttgcactttcaatgacaaactaataaattttcttattgtctatttctcgtagATACACCACTGACCAAGGCAATATGGACCAGCAGTTCTGTCTTCGCTGGAATAATCATCCGACCAACTTGACTGGCGTGCTGACTTCTTTGCTCCAGAGGGAGGCCCTCTGTGATGTTACATTGGCATGCGATGGTGG
-1fr:                                                                                                      ··Y··T··T··D··Q··G··N··M··D··Q··Q··F··C··L··R··W··N··N··H··P··T··N··L··T··G··V··L··T··S··L··L··Q··R··E··A··L··C··D··V··T··L··A··C··D··G··G
              954
              |
 0.89 tgtcaagGTatttat
DNA:  TGAAATTGTCAAGgtatttattttattacactgcaattactgaaataattatcttttttttttgttgataagttcatccctcttgagtgtcataaaacatgttcgcgaatttcttttagtatagtgcaaaatgtaggaaaaacaaagaatagtaattatatagtacccgagtcttaatgttttcgtcaccggttcctaatttaattatactattatactattagtatcaaaaaattacc
-1fr: ··E··I··V··K·

                                                                                                                                                              0.98 ctttt
DNA:  ttaatagaagaaaagacagacagatattgacgtcaatcgatatgtggagctagaaaccttgggcaacgcgttaaaaagattcaaaaactcgctttttgctccatctcactctcgcctacatcaccatcaatgtcttggactatctcactttgtcttgtcttgtgagacacaatattctaagatgtagaaaaaaaaattcttggtgtcattgaaaagaaaatgcaaatacgtaacctttt

                   450                                                                                                                                    250
                   |                                                                                                                                      |
      ccttcaccattttcAGgcacatcagaccatcctgtc                                                                                                          0.72 tttacagGTaataca
DNA:  ccttcaccattttcagGCACATCAGACCATCCTGTCAGCATGCAGTCCATATTTTGAAAGCATTTTCCTGCAGAACTCTCATCCGCATCCAATAATATATTTGAAAGATGTTAGATACTCAGAGATGCGCTCACTCCTTGATTTCATGTACAAGGGTGAGGTGAACGTTGGCCAAAGATCGTTACCGACATTTCTAAAGACAGCCGAAAGTTTACAGgtaatacaagtgattgctttca
-1fr:                 ·A··H··Q··T··I··L··S··A··C··S··P··Y··F··E··S··I··F··L··Q··N··S··H··P··H··P··I··I··Y··L··K··D··V··R··Y··S··E··M··R··S··L··L··D··F··M··Y··K··G··E··V··N··V··G··Q··R··S··L··P··T··F··L··K··T··A··E··S··L··Q·

DNA:  taataattttttcccataattctgccagcatcttaaattgttgtttagtgtcagaattcagcataattctcacaattacatcccagcggtagcacatataattccaattaaagcttctttcctgagaattggcagagaaatgatcaagaattagtattagagttgaagatttttctctagatggaatttttcaaatacaattgcgaaaattcctttcgtaattttgacataaaatcta

DNA:  aaattaaaaaa
```

**Figure S30. Manually-curated *P. bergeroti fruitless* partial gene model**. The genomic region regulated by sex-specific alternative splicing is included. Exonic sequences are indicated by black cases. Black upper cases indicate coding sequences. Intronic sequences are indicated by gray lower cases. Start and stop codons are highlighted in green and red, respectively. In yellow boxes, putative TRA/TRA-2 binding sites. Exon start/end positions in the scaffolds are indicated. Intron boundaries were predicted using transcripts vs genome alignment and confirmed by *de novo* prediction with Berkeley BDGP Splice Site Prediction Tool with default parameters (http://www.fruitfly.org/seq_tools/splice.html); prediction scores are indicated in red.

### *P. duboscqi fru* partial gene model

**Contig_1876045_Length=2257**

```
DNA:  gtgaaagattacacactataatagtgtaaagttaaattaaatttcagatgaattgttcttttgcgttattttgtattaaaaaccatttgaattgatagattttcgcttatatctctgtaaaaatatataaaattttgaaattttctatgttatttttagccctaaatatatcaaaataaaaatttctgtatttcgttaaaatattcaaaattttgttaaaaaggaacaggaaaattcca

                                                                                           1935
                                                                                           |
DNA:  taatattataatattttaactaaaacttgtgtttcttatctgtctatttacagagcttgagaagagtaagtgaccgcatagaATGATGACTACTCCGGACATCTTCGGCAGCCCCTTTCCACCGTATAGGGGTCAGCCGTTGGCTGTACTTCCACCACGTGATGACAGTCCGCCAACGCCGGCACTTGACCTCAAACGCTATGCCAACACTGAGGTCCCTCCGGCATCAACAACACCT
-2fr:                                                                                      ·M··M··T··T··P··D··I··F··G··S··P··F··P··P··Y··R··G··Q··P··L··A··V··L··P··P··R··D··D··S··P··P··T··P··A··L··D··L··K··R··Y··A··N··T··E··V··P··P··A··S··T··T··P·

                                                                                                   1685
                                                                                                   |
                                                                                       0.96 cacctagGTaggttc
DNA:  GTCGCACCACACCTCCGTTTACCGCTGCATCATGCACATGAACGTGAAATGCCGATACCGCACCAGCGTCCATCTTCGGCACAAACACCACCTAGgtaggttctacctcctgggcgctagccaaattagcaggacacagtgcgacagatcacatctgcatctctcaggagaaaagTCTGCAATCAACAtgtctcgctgtgtcttatgtacagaatgggtatggggaagtagtgcgctcc
-2fr: ·V··A··P··H··L··R··L··P··L··H··H··A··H··E··R··E··M··P··I··P··H··Q··R··P··S··S··A··Q··T··P··P··R··*·

DNA:  cctgccttatttcctcatccactgtgaaaaccctgatccaacccgtcatccctgcgtgagcaacaacctaacatacaaatttcttaaccataaatctaacaccaaaaggtttctttcaaattataagcatgcgtcattctcactgtttaaaagagatacgagatccaggatatgagcaagtgatacagtacaaaaaaagaagtttccgagtatggaaaatctagaaagattttccttt

DNA:  gcaaaaaaaagtaaccgaaaaaaattgcaatcgcggagtttttaatatattgctccgatagtaacttttgtatgcaatatagaaaaatttctaccgccagaataaggaagaaatgtagagcaacaggagcaaaaatataatagtagaagagtagaaaagtttgctaaagaaagtcgaattagataaacaagaaatgacttgaagaaattcaaatagaaatgctttttgccaataatac

                                                                                                                                        920
                                                                                                                                        |
                                                                                                                                 1.00 gaatttgGTaagtgc
DNA:  ctatttattgcaatagtcgtaaagaatcattataTTTTCAATCAACAtttgccacgattaacatgcatttgaacaaaaataaaTCTACAATCAACAttcatcaataacggccacgtgcctcgaaagaaacaaactggaatttggtaagtgcattgttagaagtaaggcattttccactgttttctgaaaagtgtagtacattgtatctgtattccggtgacttcccataaaaggaatt

DNA:  cttttcaatcttcgggcatgtttgacttgtgcaagatgcaagcacacccctgatgcccaaggtccttacggtctctgtatatcaaaaggattctccagaaaatggccaaaggtttatatatattgaaatgcagctttctgcagaattttcacgaaacaattttgaacattcatggtggaggaggagaaaatcgccgccaagcataaaagcactctcttgagatggatgtaaaatgccaa

DNA:  gagttgcgcccaagaatcatacaatagggcctgagtgtaattttggacggtatcaaacgtacaacacccccgtagtatgtttcaacgtttttgagcctcaccaccttgcacttgtacttcatgatgccactgaataatacattaccatctcgtattcatttaatatcattttccaatgtgaatcttcgtcagattccctcgcgagtgaatctttcatcaagactgtgatcttgagatc

DNA:  ctcctattttaacttccgatttcatcatgttcttctaaatttctctgcagaagcatttactgcaaaagcataattgaattaaatctcgatgattttctaggagagttactgaaattttatgagtacaaaaggtgaacagtacacccaatgaaatgaaaagtctttaagcactgatagttttatttttgttctacagggaattgatcaatttatgtaatctcttaattacaaagtaaca

DNA:  ccttggtttaattccatcagaattttatttctccttccacattttcaccactgcttttaaatagaagtctgtctttgataaataataaataataataataatgctg
```

**Contig_84267_Lenght=896**

```
DNA:  aaaaatatgtatcttaactatttgcaatgaaagatagtttctgggatttaattgatcaataaaaatatattcacttaaacaatattataaggctgtgtaatatatttatatagcatcagtcttttaatctttaacatattcgatcttttacaaatatttttaactgaaattgaaaaaggctttggcaaattaatttaacaaaatactttttttcgtactctcctattcatttgggagact

DNA:  tttttgagacaatcttcctgggtcaccctgttcataaaactttgagtgttaatgaaggtttttttgcctcttccatgtgtgttctccgacgcgaatagaaaaagacagatatgacagtatgttgcaaaatttagaataaatttatttctagatgctcatttttcgcaaacacctcgcccatgtataagcgccaaaaggtttttgcaggtggaactggacatggtcaataaaaaat

                                                                                                                                              624
                                                                                                                                              |
                                                                                                                   0.99 cttattgtctatttctcgtAGatacaccactgaccaaggca
DNA:  tagtctttctaccttgatacaatccgtgtgcaaaactttccattaggtgcttttttttggtaaaatagtcgaggagtgatattaatgaaaagttttgcactttcaatgacaaactaataaatttcttattgtctatttctcgtagATACACCACTGACCAAGGCAATATGGACCAGCAGTTCTGTCTTCGCTGGAATAATCATCCGACCAACTTGACTGGCGTGCTGACTTCTTTGCT
-1fr:                                                                                                                                              ··Y··T··T··D··Q··G··N··M··D··Q··Q··F··C··L··R··W··N··N··H··P··T··N··L··T··G··V··L··T··S··L··L

                                                        121
                                                        |
                                             0.89 tgtcaagGTatttat
DNA:  CCAGAGGGAGGCCCTCTGTGATGTTACATTGGCATGCGATGGTGGTGAAATTGTCAAGgtatttattttattaaattgaaatacttaagtttttaaataagttcatccctcttgagtgccataaaacatgttcgcaaatttcttctagcgttgtgcaaaatgtagaaaaacgaaaaaa
-1fr: ··Q··R··E··A··L··C··D··V··T··L··A··C··D··G··G··E··I··V··K·
```

**Contig_34439_Length=735**

```
DNA:  tctccggttcctaatttaattatactattatactattagtatcaaaaaattaccttaatagaagaaaagacagacagatattgacgtcaatcgatatgtggagctagaaaccttgggcagcgcgttaaaaagattcaaaaacttgctttttgctccatctcactctcgcctacatcaccatcaatgtcttggactatctcactttgtcttgtcttgggagacacaatattctaagatgt

                                                                                310
                                                                                |
                                                       0.98 cttttccttcaccattttcAGgcacatcagaccatcctgtc
DNA:  attaaaaaaaaattcttggtgtcattgagaagaaaatgcaaataagtaaccttttccttcaccattttcagGCACATCAGACCATCCTGTCAGCATGCAGTCCATATTTTGAAAGCATTTTCCTGCAGAACTCTCATCCGCATCCAATAATATATTTGAAAGATGTTAGATACTCAGAGATGCGCTCACTCCTTGATTTCATGTACAAGGGTGAGGTGAACGTTGGCCAAAGATCGTTA
+3fr:                                                                          ·A··H··Q··T··I··L··S··A··C··S··P··Y··F··E··S··I··F··L··Q··N··S··H··P··H··P··I··I··Y··L··K··D··V··R··Y··S··E··M··R··S··L··L··D··F··M··Y··K··G··E··V··N··V··G··Q··R··S··L·

                                   511
                                   |
                          0.72 tttacagGTtaataca
DNA:  CCGACATTTCTAAAGACAGCCGAAAGTTTACAGgtaatacaagtgaatagctttcataatttttcccataattctgccagcatctaaattgttgtttagtgtaagacattcagcataattctcacaattacatcccagcggtagcacatataagtccaattaaagcttctttgctgagaattggcagagaaatgatcgaaaattagtattagagttgaaatgaattattaatagatttt
+3fr: ·P··T··F··L··K··T··A··E··S··L··Q·

DNA:  ttttctctagatggaatt
```

**Figure S31. Manually-curated *P. duboscqi fruitless* partial gene model**. The genomic region regulated by sex-specific alternative splicing is included. Exonic sequences are indicated by black cases. Black upper cases indicate coding sequences. Intronic sequences are indicated by gray lower cases. Start and stop codons are highlighted in green and red, respectively. In yellow boxes, putative TRA/TRA-2 binding sites. Exon start/end positions in the scaffolds are indicated. Intron boundaries were predicted using transcripts vs genome alignment and confirmed by *de novo* prediction with Berkeley BDGP Splice Site Prediction Tool with default parameters (http://www.fruitfly.org/seq_tools/splice.html); prediction scores are indicated in red.

### *P. bergeroti dsx* partial gene model

**Contig_33004_Length=2707**

DNA: gcccgaatttgagagagtctactgtatggtgagcattttactgtcaatttgattcttctctgaactgaatttagaacttctttttccgagtgtatgtcgactcgattttgcaaataaaggggagcttctggacatatatctctctcactttcatcgatacggtttttcttttcgcaattattgcacacactttcatgttccatgtgcgccactgcaacaccaacaagcaaatggtcagac

DNA: ggtgggtagtaacattgcatatgctatatgttgatgcatagcagacatttagttctgaacttgcacacactttcatgttccatgtgcgccactgcaacaccaacaagcaaatggtcagacatgtgtgagtaatgattgcaaatatatattcaagtgaatgacgttgagcggcgaaatatcttggcattgacgcgatttagccgcgaagagagaactcaagcactcgtcagtgtgtcgata

DNA: cgtatgggattgtattatatgatctgtatgcactatcgcaaccagaccgaaagacccatggttatgtattcagttgcaacccgtcctccgttcgtgtcacactggctttcaaattcctgtcgctacaattcgatatcctcagtttgtttgacaccaccatcggcaacagttgcactgaatcccaaaaataagattccgccgagattttcctaatcagcttgtgcatttcgataaatcgtg

DNA: taccacgcggcatagcgctgcatggtgctttgtgctcgagtaataatttctttttcttctttggacaatttactgcttgtgtgaatattgagaaggtgaaattgagagattgttagttgagaataattgtttcacatgaaaatattctcctgtgaaatttctgcaaaagacataggagtgagacacaaaaaaagagggaagtttcattccttgtacttttaataaaatactgactgaatt

DNA: ccaatgttactaccccatggtgttgattgaatgaagtgcaaagcagaaaatgttgattaattgcctgtaataccacaaacgttcattggtgaatcttcacgtaatgatcatcaaaaataaactcaagtcctaaagcattatatcggcataacgtctgaatctgacgaagaaagaaaaaaaaaactataaatactccattagctcatttgattgtgaaggtgaatttattaaatgtcttct

DNA: ttgattgcaacattggcgcactttcttacccaagagagcttctcaactgagaagaacctggggtagtcggttgcatgtgcactaaacttatatgccaacaattgtatacattttgtgacttgttgcttgctacgaagtacataaatacagtggaagtggcaattatataggcacttttatcatacagctctcaccacgaaaagaaaaaacaatatagcagaggaggtagacctcttagc

DNA: ttggtactgaagaaaaaagtttgagtttgtgtagaattgcaagctttcgacctgctactaccaagactatttatttttattttaattgtgcaaactttgtgcagaaaacgtgccatctgcgagagactcacagaggttaaatctcttccgtgatccacatcagtgacaattaatgattctaacttagaagattatcatactgatttctatataagatcaatccatttaaactgttcaag

DNA: acataccttaggttttgatcaaatacccatcgctagcaaagtgttggggtcttcaatcatatcgcatttgaaatccctaccctcgagacattggccagaaagtcttatcgttaattgacgatattttcttgagtcatacagagttatgccagaagacaatctgtataagaaggaatgagagaaagaagtggcaaaaagtgatgaatgtttgcctttggttggggtaacttcggcagatgt

2091
|
DNA: ctacagacctgtaaaattgcaataagagacaatttaaagacattaaaacatctgacaagtgattattgcaaattcagtaacattgtgtgcaaaaaagtgactaaattctctgtataatctagactaagtttcagtcttcggtcaaaggttcaattaagtgggacgctgttg**ATG**GTGTCGGAGCACAATACGTGGAGCGACGTGATGTCCAATTCGGACCTCACAGACTCGAAAAACGAG
+1fr: ·M··V··S··E··H··N··T··W··S··D··V··M··S··N··S··D··L··T··D··S··K··N··E·

DNA: **TTGTGCGGAGGAGCCTCGAGCTCCAGCGGCAACTCATTGAGCCCGAGAACGCCACCAAATTGCGCCCGCTGCCGCAATCACGGCCTCAAGATAGCCCTTAAAGGTCACAAACGCTACTGCAAGTACCGTTACTGCAACTGCGAAAAGTGTCGTTTAACGGCCGAACGTCAGCGGGTGATGGCCTTGCAGACGGCCCTAAGACGTGCTCAGGCTCAGGACGAGGCACGTCAGATCTCCGCG**
+1fr: ·L··C··G··G··A··S··S··S··S··G··N··S··L··S··P··R··T··P··P··N··C··A··R··C··R··N··H··G··L··K··I··A··L··K··G··H··K··R··Y··C··K··Y··R··Y··C··N··C··E··K··C··R··L··T··A··E··R··Q··R··V··M··A··L··Q··T··A··L··R··R··A··Q··A··Q··D··E··A··R··Q··I··S··A·

2578
|
1.00 gctgcaaGT
DNA: **GATGAGGTGCCACCGACGCCACCGCCAGCCCCGTCCATTTGCACCCTTGCCACTCCTAAGAGTCAGTGCGATAGCGAACCACCGGCACCAATTACTGTGCCCACACCGGCACGTTCCCTAGAGGGTTCCTGCCACTCATCATCGGCCACACCCTGTTCTGCTTCCGGAGAACCCATCTCCGTGCCATCCAGCCGCAAATTGGCCCCAGTCAATCCAACCATCAGAAATCCAGCTGCAA**gt
+1fr: ·D··E··V··P··P··T··P··P··P··A··P··S··I··C··T··L··A··T··P··K··S··Q··C··D··S··E··P··P··A··P··I··T··V··P··T··P··A··R··S··L··E··G··S··C··H··S··S··S··A··T··P··C··S··A··S··G··E··P··I··S··V··P··S··S··R··K··L··A··P··V··N··P··T··I··R··N··P··A··A··

aagtac
DNA: aagtaccattcaaattatccttacaacatataaggtactctctacaacagctaccctacttatggcttcaaacattaaaaaagaaatatgcttaatcctaagagttcaggaattttttgtaaaacttt

**Contig_197637_Length=1025**

DNA: tccttctattcttagtcattctatttccttatcaattagtcaacagacatagtaaaagtacaggtgaagtgtcttgaggtgtgtagtttttttattcgcaacttgatttcatcaaaaggcactcaagatgatggtgaaggaaaaagtctttgccaatcaatgacttcaacaattatttcaatttaatattatttaccctttttggtatcgggaaagatgcttttaaaagatatcatcgtgt

551
|
0.93 ttcctcttgtcacgctttcAGgtcaaa
DNA: ggcagagtccgtggagatcacagattcgcaataattttatattgacgcaccttttgatatataatatttgattttaatggaagctaatgatagactgcaacacgcaaatatttcctttttactattcattaaataccatgccataaaaaagagccaacacttaaactagattctgtaattgatggaatattttcttgattttaatgtacaactcttcctcttgtcacgctttcag**GTCAAA**
-3fr: G··Q··

407
|
gctccgaagtcttc 0.99 gaagaagGTaagatg
DNA: **GCTCCGAAGTCTTCTTTGACAATTGCCAGAAATTGCTGGATAAATTCAATTACCCTTGGGAACTGATGCCCCTAATGTATGTGATCTTGGAAAATGCCAATGTTGACGTGAGGGAAGCGGCACGGAGAATTGAAGAAG**gtaagatggatttttattgatttaaaaaaaaatgaaaaatatcttttcggttttacgtgaaacccattgcattgtgattcctgaaatgattttgagatgaaa
-3fr: S··S··E··V··F··F··D··N··C··Q··K··L··L··D··K··F··N··Y··P··W··E··L··M··P··L··M··Y··V··I··L··E··N··A··N··V··D··V··R··E··A··A··R··R··I··E··E··

DNA: gacataaatttggcatgagatcctactagataatccaatcaaatcaaatttattaagaagaaggttccgttgttaagaactagtttagctagattccatcgactaataaaattgaagaaaataaatcttcctgttgtagagacaattacttttatgtcaaattagatcaatttcaatttagtagtctataggcagtacaagacttttaactattaaaatttttatttactgatgaagaac

DNA: ttaatatgcaagttttatgtgaatagaacattagttaaattacgtctaagaaattgcataatggt

**Contig_120456_Length=776**

148
|
0.24 atttaccacttcacgatctAGgtcaatgcgttgtgaatgag
DNA: aagttacttaagacacctaaggcaatgccctttctaattcgattcagtttcaaatgttactcattttcaatgggatttgagtgcattttagcgaggattaaactttgacaaatgatttctaaacaacatttaccacttcacgatctag**GTCAATGCGTTGTGAATGAGTATACTCGAATGCATAATCTCAATATGTATGATGGAGCCGAATTAAGGGGCTCCACCCGGCAATGCGGATGA**
-1fr: G··Q··C··V··V··N··E··Y··T··R··M··H··N··L··N··M··Y··D··G··A··E··L··R··G··S··T··R··Q··C··G··*·

DNA: **TAA**tttactgccagcagtttcatatgttagtaaaagaaaaaaacgt**TCATCAATCAACA**ttgagagaatatgctaatgttggcactcgtacacacctaagttcaaactaagatatatggtaacttaacccaagaaacgcgtatttgaaagaaagatcaaccccgttaagcttacacaaaactcttctattgattagatttactaattaattgtagatgtacataatatgagtacggccac
-1fr: ·*·

DNA: ttcttttcattctttttatatatttagtaattaatatcgtttaaaacttcttggagtgcgcatttttatttgcttaaaaagaatatttgtggataattattgttttaaatgattatttatgattgttcgcggtagtggctattggtgttaaagtgat**TCTTCAATCAACA**ttt**TCTTCAATCAACA**tt**TCTTCAATCAACACTTTCAATCAACA**aatctgcaatggaaataaaaaatcaa

DNA: actcaaaaatcacaaaacatatctttcataatttttgactcaattataatgatgtt

**Figure S32. Manually-curated *P. bergeroti doublesex* partial gene model**. The genomic region regulated by sex-specific alternative splicing is included. Exonic sequences are indicated by black cases. Black upper cases indicate coding sequences. Intronic sequences are indicated by gray lower cases. Start and stop codons are highlighted in green and red, respectively. In yellow boxes, putative TRA/TRA-2 binding sites. Exon start/end positions in the scaffolds are indicated. Intron boundaries were predicted using transcripts vs genome alignment and confirmed by *de novo* prediction with Berkeley BDGP Splice Site Prediction Tool with default parameters (http://www.fruitfly.org/seq_tools/splice.html); prediction scores are indicated in red.

### *P. duboscqi dsx* partial gene model

```
Contig_42106_Length=2817

DNA:  ttgtttttctgagtgtatctcgactcgattttgcaaataaaggggagcttctggacatatatctatctcactttcatcgatacggtttttcttttcgcaattattgcacacactttcatgtcccatgtgcgccactgcaacaccagcaagcaaatggtcagacggtgggtagtaacattgcatatgctatatgttgatgcatagcagacatttagttttgagcatgtgtgagtaatgatt
DNA:  gcaaatatatattcaagtgaatgacgttgagcggcgaaatatcttggcattgacgcgatttagccgcgaagagagaactcaagcactcgtcagtgtgtcgatacgtatgggattgtattatacgatctgtatgcactatcgcaaccagaccgaaagacccatggttatgtattcagttgcaacccgtcctccgttcgtgtcacactggctttcaaattcctgtcgctacaattcgatatc
DNA:  ctcagtttgtttgacaccaccatcggcaacagttgcactaaatcccaaaactaagattccgccgagattttcctaatcagcttgtgcatttcgataatcgtgtaccacgcggcatagcgctgcgcggtgctttgtgcttgagtaaaaatttctttttcttctttcaacaatttactacttgtgtgaatattgagacggtgaaattgagagattgttaggttgagaataattgtttcacct
DNA:  gaaaatataattctgtgaaatttgtgcaagagacataaagtgagactcaaaataataaaataagaaactttcaatccttcaacttttgctaaaatctgactgaattccaatgcactaccccatggtgttgattgaattgaagtgcaaagcagaaaatgttgattaattgcctataagtccacaagcttgctcattggtgaatcttcgcgtaatggtcatcagaaataaactcaagtccta
DNA:  aagcgttacattggcataacgtctgaatctgaagaggaaagaaaaaaagactataaatactccattagctcattttgattgtgaaggtgaatttattaaatgtcttctttgattgcaacattggcgcactttcttacccaagagagcttctcaactgagaagaacctggggtagtcggttgcatgtgcactaaacttatatgccaacaattgtatacattttgtgacttgttgcttgct
DNA:  acgaagtacataaatacagtggaagtggcaattatataggcacttttatcatatagctctcaccacgaaaaaaaaaacactgtagcagaggaggtagacttcttagcttggtattgaaaaaaaaagtttgagtttgtgtagaattgcaagctttcgacctgttacaaccaagactttatttcttattattgaactgtgcaaagtttgtgcagaaaacgtgccatctgcaagagactcaca
DNA:  gaggtaaaatctcttccgtgatccacatcagtgacaattaatgattctaacttagaagtttatcatactgacttctatataagattgatccatttaaactgcttaagacataccttaaggtttgatcaaatacccgtcactaacgaagtgttgggatcttcaatcatatcgcatttgaagtccctaccctcgagacattggtcagagagtcctatagttaattgacgatattttcttggg
DNA:  ttatacagagttatgccagaagacaatctttgagaaggaatgagagaaagaagtggcaaaaagtgatgaatgtttgcctttggttggggtaacttcggcagatgcaaagacttgtaaaattgcaataagagacaatttaaagacattaaaacatctgatcagtgattattgcaatttctgtaacattgtgtgaaagaagtgactaaagtctctgtatatctaaactaaatctcagtcttc

                                  1953
                                  |
DNA:  ggtcaaaagttcaattaagtgggacgctgttgATGGTGTCGGAGCACAATACGTGGAGCGACGTGATGTCCAATTCGGACCTCACAGACTCGAAAAACGAGTTGTGCGGAGGAGCCTCGAGCTCCAGCGGCAACTCATTGAGCCCGAGGACGCCACCAAATTGCGCCCGCTGTCGCAATCACGGCCTCAAGATAGCCCTTAAGGGTCACAAACGCTACTGCAAGTACCGCTACTGCAACT
+3fr:                                   ·M··V··S··E··H··N··T··W··S··D··V··M··S··N··S··D··L··T··D··S··K··N··E··L··C··G··G··A··S··S··S··S··G··N··S··L··S··P··R··T··P··P··N··C··A··R··C··R··N··H··G··L··K··I··A··L··K··G··H··K··R··Y··C··K··Y··R··Y··C··N··

DNA:  GCGAAAAGTGTCGTCTAACGGCCGAACGTCAGCGGGTGATGGCCTTGCAGACGGCCCTAAGACGTGCTCAGGCTCAGGACGAGGCACGTCAGATCTCCGCGGATGAGGTGCCACCGACGCCACCGCCAGCCCCGTCCATTTGCACCCTTGCCACTCCTAAGAGTCAGTGCGATAGCGAACCACCGGCACCTATTACTGTGCCCACACCGGCACGTTCCCTAGAGGGTTCCTGCCACTCAT
+3fr: C··E··K··C··R··L··T··A··E··R··Q··R··V··M··A··L··Q··T··A··L··R··R··A··Q··A··Q··D··E··A··R··Q··I··S··A··D··E··V··P··P··T··P··P··P··A··P··S··I··C··T··L··A··T··P··K··S··Q··C··D··S··E··P··P··A··P··I··T··V··P··T··P··A··R··S··L··E··G··S··C··H··S··

                                                                                                         2499
                                                                                                         |
                                                                                            1.00 actgcaaGTaagtac
DNA:  CATCGGCCACACCCTGTTCTGCTTCCGGAGAACCCATCTCCGTGCCATCCAGCCGCAAATTGGCCCCAGTCAATCCAACCATCAGAAATCCAACTGCAAgtaagtaccattcaaattgtccttgctacgcccatagtatcctctacgacagctatcctacttatggccatatatactaaaaaaaaataagcttaatactattctctgagttaaagaaagactttaaaaaccttggatgaa
+3fr: S··S··A··T··P··C··S··A··S··G··E··P··I··S··V··P··S··S··R··K··L··A··P··V··N··P··T··I··R··N··P··T··A··

DNA:  gactgtaatttgacgtaaagtagatgcgagtgactttttttcccagtcgatgactttgaccccttcctgtttataagctttcctttaattctagtgtttaagaatgattttcaccgatcagatatgattaataggatcagtaaagaccatccttgggcactcaatgggtcaagtggta


Contig_146905_Length=1363

DNA:  aataaagtcgaaagacatgaaaactgtcagcaaaaattcttctattcttggttattctacttccttatcaattagtcaacagacatagttagagtgcaggtgaagtgtcttgaggtgtgcagtttttttactcacaacttgatttcatcaaaaggcactcaagatgatggtgaaggaaaaagtctttgccaatcaatgacttcaacaattatttcaatttaatatgatttaccctttttt
DNA:  gtatcggaaaagatgcttttaaaagatatcatcgagtggcaaagtccgtggagatcacagatttgcaataattttatattgacgcaccttttgatatataatatttgattttaatggaagctaatgatagactgcaacacgcaaatatttccttttactattcattaaataccatgccataaaaaagagccaacacttaaactagattctgtaattgatggaatattttcttgattttaa

                                852                                                                                                                    708
                                |                                                                                                                      |
       0.93 ttcctcttgtcacgctttcAGgtcaaagctccgaagtcttc                                                                                              0.99 gaagaagGTaagatg
DNA:  tgttcaactcttcctcttgtcacgctttcagGTCAAAGCTCCGAAGTCTTCTTTGACAATTGCCAGAAATTGCTGGACAAATTCAATTATCCTTGGGAACTGATGCCCCTAATGTATGTGATCTTGGAAAATGCCAATGTGGATGTGAGGGAAGCGGCACGGAGAATTGAAGAAGgtaagatggaattttattgatttttataaaaattgaaaaatatctttttcgggtttacgagaacc
-1fr:                                 G··Q··S··S··E··V··F··F··D··N··C··Q··K··L··L··D··K··F··N··Y··P··W··E··L··M··P··L··M··Y··V··I··L··E··N··A··N··V··D··V··R··E··A··A··R··R··I··E··E··

DNA:  ccattgcaatgtgattcctgaaatgattttgagatgaaagacataaatttggcattggatcctactagataatccaatgaaatcaaatttattaagaagaaggttccgttgttaagaactagtttagctggattccatcgactaataaaattgaagaatataaatcttcctgttgtagaatcaatttcttttatgtcaaattagatcaatttcaatttagtagtctataggcaggacaa
DNA:  gacttttaactattaaaattttttatttactgatgaagaacttaatatgcaagttttatgtgagtagaacattagttaaattacatctaagagattgcataatggttgcaatatcaaattttatagtactggacaataaaatgttgagattcattaagaaaatattacattacacacttttctcaggcaattgtcaagctctgacgagaatatcttctttcaacatattaatttattatga
DNA:  tagttgaataataaagtatgttgtacatattatgtttatcaaaggtgtacgaatatagttgtttaaactgaaaagtgtaatttaaaaagaaaatcttggagattttcaaatagtttttttagaatttttttaagtcctctaataagaaacataacttttaaaa


Contig_213005_Length=2151

DNA:  attataaagtttccaaggatttgcagtttcttaaatttgctaatttgttatttttttatttttaaaattaatgatttccgttttgtgtatattattttgtacagaccattttaattaaaaaaaactaatcagtttgctgtcaccttttaaatgaagtaaaatatacaaaagtgttacatttcaattctgagataattgatgtgtggattattgcgttccattaatcaagctttacatttt
DNA:  tactttattaagaagcaaaattgagatatttgaattaagactagcgacagttagattttaattgtcgttgtaaaattttaacatcatatttttggactgaaagttttattttaaatttaaatcgaatatttgatatatgtttaacatataaattcagtatgttgtagagttcaattgtggtaacattagattaattgacctaagtaagatgatactggaatgaaatccttagttatttg

                                                                                                                                                            0.24 atttaccacttca
DNA:  gatttggaatatcacaggaatatttgatatctttcagtcttcatcaaaatcatttcaggaatcattgaagataaaatttctcctcaattgggagagtttaagttacttaagacaccttaaggcaatgccttttctaattcgattcagtttcaaatgtaactgattctcaatgagatttgagtgcatttgagcgaggattaagctttgacaaatgattgctaaacaacatttaccacttca

              1423
              |
      cgatctAGgtcaatgcgttgtgaatgag
DNA:  cgatctagGTCAATGCGTTGTGAATGAGTATACTCGAATGCATAATCTCAATATGTATGATGGAGCCGAATTGAGGGGCTCCACCCGGCAATGCGGATGATAAtttactgccagcaaaaaaaaatgtTCATCAATCAACAttgagagaatatgcaaatgttggcactcgtacacacctaagttcaaactaagatatatggtaacttaacccaagaaacgcgtatttgaaagaagatcaac
-2fr:         G··Q··C··V··V··N··E··Y··T··R··M··H··N··L··N··M··Y··D··G··A··E··L··R··G··S··T··R··Q··C··G··*··*·

DNA:  cccgttaagcttacacaaaactcttctattgattagatatactaattaattgtagatgtacataatatgagtacggccacttcttttcattctttttatatatttttttagtaattaatatcgtttacaacttcttggagtgcgcatttttaattgcttaaaaagaatatttgtggataattattgttttaaatgattatttatgattgttcgcggtagtggctattggtgttaaagtgatT

DNA:  CTTCAATCAACAtttTCTTCAATCAACAttTCTTCAATCAACATCTTCAATCAACAaatctgcaatgaaaacaaaaaaaaatctaaaaatcacaagacatatcttccataatttttgactcaattgtaccaatgtgtttttttggcgggaaatgtcaaaaggacccacttaaatcagactctctagagagaagaaaatcaatcaaccatttgaatttagccacaatcttaacagtacatcca

                                                                                                                                      547
                                                                                                                                      |
                                                                                                                           0.99 gcgaatgGTaagaag
DNA:  ttcacgatgtgagaaatgggcagcgtttcatcaaattgacaacaataaagaagtggatgccagaagattgcaacagcgtaaaaaaagtggaacaacagcaaaaaagtgcaaggagaaaacacattaatgttcaaaggcgaatggtaagaagttgatgtagtcaaagtagaaaatgctttcacggggtgtaattaattctataagaattcatatgcacattaaacgtaatcataaaagttga

DNA:  aaataaatgaacttcttgtcttactccaacctctaaggactataagtgagaaattttatattaaaacagcaaagtgctataagtaatttcatactctaatctctttttcatcataccatctcttttcacttccacctattttctacaatattttttcccaagctctttctgttatataaaagttattagtttgaagagatttcttttgtaaagttcaaaagctactcccttgtttttctc
DNA:  tccattcaatttattcgtcttcatcgtcatcaaattatgttccatttattccctatatatgtactgttttagttcctttgccccatgttggttgaggaagaaaaaacctcttgaggagtgttgaaaagattgatggaattgcccagacaatttcaatttattgcaagttaattaatagtgaggaataaaagttattgttattttggtac
```

**Figure S33. Manually-curated *P. duboscqi doublesex* partial gene model**. The genomic region regulated by sex-specific alternative splicing is included. Exonic sequences are indicated by black cases. Black upper cases indicate coding sequences. Intronic sequences are indicated by gray lower cases. Start and stop codons are highlighted in green and red, respectively. In yellow boxes, putative TRA/TRA-2 binding sites. Exon start/end positions in the scaffolds are indicated. Intron boundaries were predicted using transcripts vs genome alignment and confirmed by *de novo* prediction with Berkeley BDGP Splice Site Prediction Tool with default parameters (http://www.fruitfly.org/seq_tools/splice.html); prediction scores are indicated in red.
