## Additional_file_15_Supplementary_Methods for "Genomics and transcriptomics to unravel sex determination pathway and its evolution in sand flies"

**Table of contents**

1) Cloning of *P. perniciosus fru* transcripts
2) Genome assembly of *P. bergeroti* and *P. duboscqi*
3) Manual curation of *tra*, *tra-2*, *dsx* and *fru* gene models in Phlebotominae
4) *De novo* transcriptome assembly of sand fly species
5) List of primers utilized in this study
6) Nucleotide sequences of the sex determining genes identified by *in silico* approach

**1) Cloning of *P. perniciosus fru* transcripts**

The TBLASTN search against the perniBASE dataset using the *A. aegypti* FRU proteins as query (Tab. S1) led us to identify two partially *in silico* assembled transcripts encoding for partial male-specific and non-sex-specific Fruitless (FRU) protein isoforms, both containing a well conserved BTB domain. We further searched within the perniBASE dataset by TBLASTN using as queries the *D. melanogaster* FRU Zinc-Finger terminal domain of type A, B and C (ZnF-A, ZnF-B and ZnF-C) (Salvemini et al. 2013) and we identified two transcripts, c23972.g4.i1 and c25556.g5.i1, encoding for partial proteins with a well conserved ZnF-A or ZnF-C domains, respectively. These transcripts could represent uncomplete assembled *Ppefru* transcripts. We validated this hypothesis by gene specific RT-PCR analysis, using reverse primers located in the two ZnF putative transcripts c23972.g4.i1 and c25556.g5.i1 (PpeFruZnfA-, PpeFruZnfAinner-, PpeFruZnfC-, PpeFruZnfCinner-) and a forward primer located in the *P. perniciosus* male-specific *fru* transcript c23888.g1.i4 (PpFruM+). We successfully amplified four full length cDNAs (*PpefruMA*, 2249 bp; *PpefruFA*, 3026 bp; *PpefruMC*, 1915 bp; *PpefruFC*, 2692 bp) with complete FRU ORFs in male and females of *P. perniciosus*. The four cDNAs were cloned and sequenced as described in Methods.

**2) Genome assembly of *P. bergeroti* and *P. duboscqi***

We produced a draft genome assembly of the sand fly species *P. bergeroti* and *P. duboscqi* using the MINIA assembler (Salikhov et al., 2013) and the available Illumina genomic reads at SRA NCBI Archive (https://www.ncbi.nlm.nih.gov/sra) (*P. bergeroti*: SRR1973671 and SRR1973788; *P. duboscqi*: SRR1973598 and SRR1996565) with the following command:

```
minia -in species_name.fasta -kmer-size 31 -abundance-min 3 -out species_name_assembly_k31_m3 -nb-cores 23
```

We obtained the two following genome assemblies:

| Assembly statistics | *P. duboscqi* | *P. bergeroti* |
|---|---|---|
| Total number of assembled sequences | 2240724 | 1741498 |
| Total length of sequence | 351743961 bp | 359027491 bp |
| Total GC count | 123284946 bp | 127899439 bp |
| GC % | 35.05 | 35.62 |
| Average sequence length | 156 bp | 206 bp |
| Minimum length | 63 bp | 63 bp |
| Maximum length | 32976 bp | 29034 bp |
| N50 stats >= | 190 bp | 365 bp |
| Number of Sequences longer than 10000 | 45 | 22 |
| Number of Sequences longer than 5000 | 469 | 680 |
| Number of Sequences longer than 3000 | 2802 | 4894 |
| Number of Sequences longer than 2000 | 11514 | 18082 |
| Number of Sequences longer than 1000 | 51627 | 64808 |

**3) Manual curation of *tra*, *tra-2*, *dsx* and *fru* gene models in Phlebotominae**

The *P. perniciosus* TRA, TRA-2 DSX and FRUM protein sequences and the *P. papatasi* and *L. longipalpis* orthologous predicted proteins PpaDSXF (PPATMP010078-RA and PPATMP010079-RA), LloDSXF (LLOTMP001672-RA and LLOTMP005763-RA), LloDSXM (LLOTMP005763-RA), PpaFRUM (PPATMP009581-RA) and LloFRUM (LLOTMP008996-RA) were utilized as queries to perform a TBLASTN search against the genomic data set available at VectorBase (https://www.vectorbase.org/) or the newly assembled *P. bergeroti* and *P. duboscqi* draft genomes to identify the approximate position of intron/exon boundaries. Exact intron boundaries were localized by using transcripts vs genome nucleotide alignments and were confirmed by *de novo* prediction with the Berkeley BDGP Splice Site Prediction Tool using the default parameters (http://www.fruitfly.org/seq_tools/splice.html). This approach led us to reconstruct complete *tra* gene models in *P. perniciosus*, *P. papatasi*, *P. bergeroti* and *P. duboscqi* (Fig. S10-S13), partial *tra-2* gene models in *P. perniciosus*, *P. papatasi*, *P. bergeroti* and *P. duboscqi* (Fig. S18-S20), the complete *dsx* gene model in both *P. papatasi* and *L. longipalpis* species (Fig. S23-S24), the complete *fru* gene model in *P. papatasi* (Fig. S25A-B) and a partial *fru* gene model in *L. longipalpis* (Fig. S26).

**4) *De novo* transcriptome assembly of sand fly species.**

We produced a *de novo* transcriptome assembly of the Old World sand fly species *L. longipalpis*, *L. umbratilis* and *L. neivai* using the Trinity assembler (Grabherr et al., 2011; Haas et al., 2014) and the following Illumina RNA-seq reads from SRA NCBI Archive (https://www.ncbi.nlm.nih.gov/sra):

| Species | Run | life stage | sex |
|---|---|---|---|
| *L. longipalpis* | SRR535756 | adult | female |
| *L. longipalpis* | SRR535757 | adult | female |
| *L. longipalpis* | SRR535758 | fourth instar larvae | <not provided> |
| *L. longipalpis* | SRR535759 | fourth instar larvae | <not provided> |
| *L. longipalpis* | SRR535760 | adult | female |
| *L. longipalpis* | SRR535761 | adult | female |
| *L. longipalpis* | SRR535762 | adult | female |
| *L. longipalpis* | SRR535763 | adult | female |
| *L. longipalpis* | SRR535764 | adult | female |
| *L. longipalpis* | SRR535765 | adult | female |
| *L. longipalpis* | SRR535766 | adult | female |
| *L. longipalpis* | SRR535767 | adult | female |
| *L. longipalpis* | SRR535768 | adult | female |
| *L. longipalpis* | SRR535769 | adult | female |
| *L. longipalpis* | SRR535770 | adult | female |
| *L. longipalpis* | SRR535771 | adult | female |
| *L. longipalpis* | SRR535772 | adult | female |
| *L. longipalpis* | SRR535773 | adult | female |
| *L. longipalpis* | SRR535774 | adult | female |
| *L. longipalpis* | SRR535775 | adult | female |
| *L. longipalpis* | SRR535776 | adult | female |
| *L. longipalpis* | SRR535777 | adult | female |
| *L. longipalpis* | SRR535778 | adult | female |
| *L. longipalpis* | SRR535779 | fourth instar larvae | <not provided> |
| *L. umbratilis* | SRR952151 | adult | <not provided> |
| *L. neivai* | SRR5134059 | adult | <not provided> |
| *L. neivai* | SRR5134060 | adult | <not provided> |
| *L. neivai* | RR5134061 | adult | <not provided> |

Downloaded reads were combined for each species in single or paired fastq files, were quality filtered and adaptor trimmed with Trimmomatic software (Trimmomatic-0.32) (Bolger et al., 2014) and assembled using the following Trinity commands:

*L. longipalpis*:

```
Trinity --seqType fq --max_memory 180G --left longipalpis_1.fq --right
longipalpis_2.fq --CPU 23 --jaccard_clip --normalize_reads --output
longipalpis_trinity
```

*L. umbratilis*:

```
Trinity --seqType fq --max_memory 180G --left umbratilis_1.fq --right
umbratilis_2.fq --CPU 23 --jaccard_clip --normalize_reads --output
umbratilis_trinity
```

*L. neivai*:

```
Trinity --seqType fq --max_memory 180G --single nyssomyia.fq --CPU 23 --
normalize_reads --output neivai_trinity
```

We obtained the following transcriptome assemblies:

| Assembly statistics | *L. longipalpis* | *L. umbratilis* | *L. neivai* |
|---|---|---|---|
| Total number of assembled transcripts | 236076 | 176358 | 116069 |
| Total number of Trinity genes | 166427 | 148912 | 75760 |
| Total assembled bases | 198156135 | 81656931 | 68762545 |
| GC % | 40.37 | 41.84 | 38.52 |
| Average sequence length | 839 bp | 463 bp | 592 bp |
| Median transcript length | 483 bp | 304 bp | 317 bp |
| N50 stats >= | 1368 bp | 520 bp | 921 bp |

**5) List of primers utilized in this study**

| Primer name | Sequence (5'--3') |
|---|---|
| PpeSOD+ | AGGAGGCGTCGTTCGTGAAT |
| PpeSOD- | CCTTGCCATGAGGATTGTAG |
| Ppdsx5'utrCommon+ | GTGTGTGGATTCTACTAAGG |
| PpdsxCommon+ | TCAGACGTGCTCAGGCTC |
| PpdsxF- | CGGCTCCGTCATACATATTA |
| Ppdsx3'utrF- | CTTCGCTGCTTGGGATAAGT |
| PpdsxM- | TACTCGTTGGTATCAGGCG |
| PpFruM+ | CGACAAAAGACACTCTATTTC |
| PpefruC- | ATGGTCTGATGAGCCTTGAC |
| PpFruF+ | GGATATGAGCAAGAGATACAG |
| PpFruC-nested | GTCGGATGGTTATTCCAGCG |
| PpeFruZnfA- | GTCTGACTTTGATGACTAATTGG |
| PpeFruZnfAinner- | CAATCTCGCCATTCTGGTTC |
| PpeFruZnfC- | CAACATTTTCTCCACAGAGG |
| PpeFruZnfCinner- | ACAATCTTCGTTGGTACTGTC |
| ppetra_F2inner | AATGGCGACAAAGAGGGAGG |
| Ppetra_Rinner | TTCTCCCTCAAGCGATCCCT |
| PpeTraRace3+ | CAGAGGATTCAGATAAGGAG |
| PpeTraRace3Nested+ | GAGTCGTAGTCGGAGTCTC |
| Tra-5’Race_outer | CGTCCGTTTGTGTTGAGAG |
| Tra-5’Race_inner | TATGACTTTCCTCCCTGTGG |
| PpeTra5'utr | GATCCAATTGTGCAATTGTC |
| PpeTraStop3utr- | CTATTCAGTCAATTTTCAATGG |
| PpeTra-2_F1-6-5 | CACCAGATTATGCACTTTTGC |
| PpeTra-2_R | CCTATGATACATTTACAAGTAAGG |
| PapTraF | CATGTTCCATCTACATCAC |
| PapTraR | AAGGATCGTACCAGTCATAC |
| LutzTraF | ACACAAAAGGACGGGTAATG |
| LutzTraR | TTTGTTCGTCTCCGGCGAG |

### 6) Nucleotide sequences of the sex determining genes identified by *in silico* approach

```
>Pbetra_CDS [Phlebotomus bergeroti] P. bergeroti transformer CDS
ATGATTAAAAAACCTAGCTCCAGGACGCACAAGGAAAAATATGAGGGAAAATACAAAACAAATGGCGGAAGATCAGCATCGAA
GAGGGAGGCAGAGAGACATCCATCAACACATGAAGACACCATCAAAAAATCCCATCATCATCGTGATGAATTAGGAAAGAAAG
ATGAAAGGAAGTCTTATGAGGATGATTTTCCCAAACTTGGGCATGTTCCATCTACATCACGGGATACCCGGGACGGCTTTCAA
CACAAAAGAACGGGTGATGGCAGGGAATCCTCTAAGCATCGAAGAGGTGCTGCCAAGAGAGATTCTTCAAATTCTTCAGAGAC
TGACAGTGAATCCCTCAAATCTTCGAGGAAAAGGCCAGAATCTCGCCGGAGACGATCAAAAAGTCGTAGCAGAAGTCTCGATG
ACTCGTACAGGAGACGTCAGACCTCCACGAGTCGTCGATCTTCACCACGACGCCGGCCAAAGTCTGTTTCTCCACCGCGGGAG
AAAGTGCCCTACTATGCAGATGCCCAGCGTGAGAGGGATCGCCTGAGAGAGAAGTATGGAAGTGATGGGAGGATGAGGAGGAG
GAGCAGCAGCCGCGAGAGACGCAGAAGACGCTCTAGAGATAGAAGATCCAAGACTCCAGTCCGAGAAACCACAAAAAATTGTCA
CAGTTCCTGTTCCTGTTCCCGTCTACCCATCAATTTATCCAGATGGTTCCATGTATGACTGGTACGATCCCTCTTGGCAAACA
GGGCTCTTTCCACAGGGTGTAATGCCTCCAAGACACATTCCACGCCCAATGCGACCTCATGCCCCATTCCCTACTTCACCCTT
CATGGTGGATCCTTTTCATCGGCCAATGAGGCCAGGAATGCCACCTCCAAGACCCTTTCGACCTCCCCAAGGATTCCCTCCTC
CCCGATTCCACAGACCCCATTGA
>Pdutra_CDS [Phlebotomus duboscqi] P. duboscqi transformer CDS
ATGATTAAAAAACCTAATTCCAAGACGCATAAGGAAAAATATGAGGGAAAATACAAAACAAATGGCGGAAGATTGGCATCGAA
GCGAGAAGCAGAGAGACATCCATCAACACATGAAGACACCATTAAAAAATCCCATCATCATCGTGATGAATTAGGGAAGAAAG
ATGAAAGGAAGGCTTATGAAGAAGAATTTCCTAAATTTGGGCATGTTCCATCTACATCACGGGATTCCCGGGACGGCTTTCAA
CACAAAAGGACGGGTGATGGTAGGGAATCCTCTAAGCACCGAAGAGGTGCTGCCAAGAGAGATTCTTCAAATTCTTCAGAGAC
TGACGATGAATCCCTCAAATCTTCGAGGAAAAGGTCAGAATCTCGCCGGAGACGATCTAAAAGTCGTAGCCGAAGTCTGGATG
ACTCGTACAGGAGACGTCAAACCTCCACGAATCGTCGATATTCGCCACGGCGCCGGCCAAAGTCTGTTTCTCCTCCTCGTGAG
AAAGTTCCCTACTATGCAGATGCCCAACGCGAGAGGGATCGCCTGAGGGAGAAATATGGAAGTGATGGGAGGGCTAGGAGAAG
GAGCAGCAGTCGCGAGAGACGCAGAAGGCGTTCTAGAGAAAGGAGATCCAAGACTCCAGTCCGAGAAACCACAAAAAATTGTTA
CAGTTCCTGTTCCTGTTCCTGTCTACCCGTCAATATATCCGGATGGTTCCATGTATGACTGGTACGATCCTTCTTGGCAAGCA
GGACTCTTTTCTCAGGGTGTCATGCCTCCAAGACACATTCCACGCCCAATGCGTCCTCATGCCCCATTCCCTGCCTCACCCTT
CATGGTGGATCCTTTCCATCGGCCAATGAGACCAGGAATGCCACCTCCAAGACCCTTCCGACCTCCCCAAGGATTCCCTCCTC
CCCGATTCCACAGACCCCATTGA
>Ppatra-2_CDS [Phlebotomus papatasi] P. papatasi transformer-2 CDS
ATGAGTACACGTACCTACTATAAGAACTCAATGTCAAGAAGTCCCAGCGAGGAACGACACTATATCTCTCGAGCTGGTTCATC
GCGAGAGAATGGCGACTACAGGAAGTCCCGCAAATCGTCTTCCCGCAGTCGAGCCCGCTACAGATCTCGCTCACGTTCACGGT
CTCGTTCTGAGAAGCGCTACAGAAGCAGGTCACCGTCCCGCCACGATAGGAAGCGCTCCTATTCCAGGAGCCCCAATTCCTCT
CGTCGCAGACACGTAGCAAGTCGAGTCAATTCACAGGATCATCCCCCAAAGTCCAGATGTCTAGGAGTCTTTGGACTCAGTGA
TGCAACAACTGAAGACCAGATTTATCAAATTTTCTCCAAATTTGGAACAATACACCGTACTCAAATAATTATGGATGCAATGA
CGGGATGTTCACGGGGCTTCTGTTTTGTCTACTTTGCCAATGCTGATGATGCCAAAGTGGCAAAGGATAATTGCTCAGGAATG
GAACTCGATGGGAGACGTATTCGAGTGGATTATTCCATTACTCAGAGACCTCATACTCCAACTCCAGGAGTCTACATGGGACA
ACCAACGGGCAGAGGACGAGAGAATCGAGATCGTAGTCGCAGAAATGAGTAA
>Lumtra-2_par_CDS [Lutzomyia umbratilis] L. umbratilis transformer-2 partial CDS
AGAAATCTCTTCCCACAGTCATCAGATAGCTCCGGGTACGACAGGAGGCACTCCAGGACAAAGAAGAAGCGTGAAAATGGGCA
CCACAAGAAATCATCAAAATCCCGAAGGCGCCATCGATCAAGGTCTCACTCAAAGCGACGACACTACTCACGATCACCCACTA
GGACCACCCGGAAGCGCCACATCTCACGCAGCCCAGTCCCAACATACAAGAAATATGAACAACCGGAAGAGGGAAGATGCCTG
GGTGTCTTTGGATTGAGCATGCACACAACAGAGAAGCACGTGCATGAGATCTTTTCCAAATTCGGACCCATTGAACGAACGCA
AATCATTGTGGATTCAAAGACTGGTCGATCTCGGGGATACTGCTTCGTTTATTTTGAAAATACTGAAGATGCAAAGGTAGCCA
AGGATCAATGTACTGGGATTGAGATTGATGATCGTCGCATTCGTGTGGATTATTCCCTCACAGCAAGACCCCATACACCTACA
CCTGGAATTTATATGGGTAAGGCTAGTCAACGGTGTGGGGACTCATCAACATCACGAAGCAACAGATACGAGGATACTTCTTA
CGAGAAAACTCGTAACAGATCACCATCCCCATACAAACGTAGGCGCCGAAGTCGATCTCGTTCACACTCACCAAGAGTTCGCC
GATATCATTAA
>Lnetra-2_par_CDS [Nyssomia neivai] N. neivai transformer-2 partial CDS
AGAAATCTCTTCCCACAGTCATCAGATAGTTCCGGGGACGACAGGAGGTACTCCAGGACAAAGAAGAAGCGTGAGAATGGGCA
CCACAAGAAATCATCAAAATCCCGAAGGCGCCATCGATCAAGGTCTCACTCAAAGCGACGACACTACTCACGATCTCCCACTA
GGACCATCCGGAAGCGGCACATCTCACGCAGCCCAGTCCCATCATACAAGAAATATGAACAACCGGAAGAGGGTAGATGTCTG
GGTGTCTTTGGGTTGAGCATGCACACAACAGAGAAGCATGTTCATGAGATCTTCGCCAAATTCGGACCCATTGAACGAACGCA
AATTATCGTGGATTCAAAGACTGGTCGTTCTCGGGGATACTGCTTTGTTTATTTTAAAAATACTGAAGATGCAAAGGTAGCCA
AGGATCAATGTACTGGGATTGAGATTGATGATCGTCGTATTCGTGTGGATTATTCCCTAACAGCAAGACCCCATACACCGACA
CCTGGAGTTTATATGGGCAAGGCTAGTCAACGGTGTGGGGATCGTGGGGACTCATCAACATCACGAAGCAACAGATACGAGGA
TTCTTCTTACGAGAAGACTCGCAACAGATCACCATCCCCTTATAAACGCAGGCGCCGAAGTCGATCCCGCTCATACTCACCAA
GAGTTCGCCGATATCACTAA
>PpadsxF_CDS [Phlebotomus papatasi] P. papatasi doublesexF CDS
ATGGTGTCGGAGCACAATACGTGGAGCGACGTGATGTCCAATTCGGACCTCACAGACTCGAAAAACGAGTTGTGCGGAGGAGC
CTCGAGCTCCAGCGGCAACTCATTGAGCCCGAGGACGCCACCAAATTGCGCCCGCTGCCGCAATCACGGCCTCAAGATAGCCC
```

TTAAGGGTCACAAACGCTACTGCAAGTACCGTTACTGCAACTGCGAAAAGTGTCGTCTAACGGCCGAACGTCAGCGGGTGATG
GCCTTGCAGACGGCCCTAAGACGTGCTCAGGCTCAGGACGAGGCACGTCAGATCTCCGCGGATGAGGTGCCACCGACGCCACC
GCCAGCCCCGTCCATTTGCACCCTTGCCACTCCTAAGAGTCAGTGCGATAGCGAACCACCGGCACCCATTACGGTGCCCACAC
CGGCACGTTCCCTAGAGGGTTCCTGCCACTCATCATCGGCCACACCCTGTTCTGCTTCCGGAGAACCCATCTCCGTGCCATCC
AGCCGCAAATTGGCCCCAGTCAATCCAACCATCAGAAATCCAACTGCAAGTCAAAGCTCCGAAGTCTTCTTTGACAATTGCCA
GAAATTGCTGGACAAATTCAATTATCCTTGGGAACTGATGCCCCTAATGTATGTGATCTTGGAAAATGCCCCGGACGTGAGGG
AAGCGGCACGGAGAATTGAAGAAGGTCAATGCGTTGTGAATGAGTATACTCGAATGCATAATCTCAATATGTATGATGGAGCC
GAATTGAGGGGCTCCACCCGGCAATGCGGATGATAA
>PdudsxF_CDS [Phlebotomus duboscqi] P. duboscqi doublesexF CDS
ATGGTGTCGGAGCACAATACGTGGAGCGACGTGATGTCCAATTCGGACCTCACAGACTCGAAAAACGAGTTGTGCGGAGGAGC
CTCGAGCTCCAGCGGCAACTCATTGAGCCCGAGGACGCCACCAAATTGCGCCCGCTGTCGCAATCACGGCCTCAAGATAGCCC
TTAAGGGTCACAAACGCTACTGCAAGTACCGCTACTGCAACTGCGAAAAGTGTCGTCTAACGGCCGAACGTCAGCGGGTGATG
GCCTTGCAGACGGCCCTAAGACGTGCTCAGGCTCAGGACGAGGCACGTCAGATCTCCGCGGATGAGGTGCCACCGACGCCACC
GCCAGCCCCGTCCATTTGCACCCTTGCCACTCCTAAGAGTCAGTGCGATAGCGAACCACCGGCACCTATTACTGTGCCCACAC
CGGCACGTTCCCTAGAGGGTTCCTGCCACTCATCATCGGCCACACCCTGTTCTGCTTCCGGAGAACCCATCTCCGTGCCATCC
AGCCGCAAATTGGCCCCAGTCAATCCAACCATCAGAAATCCAACTGCAAGTCAAAGCTCCGAAGTCTTCTTTGACAATTGCCA
GAAATTGCTGGACAAATTCAATTATCCTTGGGAACTGATGCCCCTAATGTATGTGATCTTGGAAAATGCCAATGTGGATGTGA
GGGAAGCGGCACGGAGAATTGAAGAAGGTCAATGCGTTGTGAATGAGTATACTCGAATGCATAATCTCAATATGTATGATGGA
GCCGAATTGAGGGGCTCCACCCGGCAATGCGGATGATAA
>PbedsxF_CDS [Phlebotomus bergeroti] P. bergeroti doublesexF CDS
ATGGTGTCGGAGCACAATACGTGGAGCGACGTGATGTCCAATTCGGACCTCACAGACTCGAAAAACGAGTTGTGCGGAGGAGC
CTCGAGCTCCAGCGGCAACTCATTGAGCCCGAGAACGCCACCAAATTGCGCCCGCTGCCGCAATCACGGCCTCAAGATAGCCC
TTAAAGGTCACAAACGCTACTGCAAGTACCGTTACTGCAACTGCGAAAAGTGTCGTTTAACGGCCGAACGTCAGCGGGTGATG
GCCTTGCAGACGGCCCTAAGACGTGCTCAGGCTCAGGACGAGGCACGTCAGATCTCCGCGGATGAGGTGCCACCGACGCCACC
GCCAGCCCCGTCCATTTGCACCCTTGCCACTCCTAAGAGTCAGTGCGATAGCGAACCACCGGCACCAATTACTGTGCCCACAC
CGGCACGTTCCCTAGAGGGTTCCTGCCACTCATCATCGGCCACACCCTGTTCTGCTTCCGGAGAACCCATCTCCGTGCCATCC
AGCCGCAAATTGGCCCCAGTCAATCCAACCATCAGAAATCCAGCTGCAAGTCAAAGCTCCGAAGTCTTCTTTGACAATTGCCA
GAAATTGCTGGATAAATTCAATTACCCTTGGGAACTGATGCCCCTAATGTATGTGATCTTGGAAAATGCCAATGTTGACGTGA
GGGAAGCGGCACGGAGAATTGAAGAAGGTCAATGCGTTGTGAATGAGTATACTCGAATGCATAATCTCAATATGTATGATGGA
GCCGAATTAAGGGGCTCCACCCGGCAATGCGGATGATAA
>LlodsxF_CDS [Lutzomyia longipalpis] L. longipalpis doublesexF CDS
ATGTCCAATTCGGACCTCACAGACTCGAAGAATGAGTTGTGCGGGGGAGCCTCGAGCTCCAGCGGCAACTCATTGAGCCCGAG
GACGCCGCCAAATTGCGCCCGCTGCCGCAATCACGGCCTCAAGATTGCCCTAAAGGGTCACAAGCGTTACTGCAAATTTCGCT
ACTGCAACTGCGAAAAGTGCCGCCTCACGGCCGAACGACAGCGGGTGATGGCCCTACAGACGGCCCTAAGACGTGCTCAGGCT
CAGGACGAGGCGCGTCAAATCTCCGCGGATGAGGTGCCACCGACACCACCGCCACTTACTGGTCAAATTGCAACAACGCCCAA
AAGTCAGTGCGACGGTGAACTACCGCGATCGAATACAGTGCCAACTCCGGCACGTTCCCTCGAGGGCTCCTGCCACTCATCTT
CGGCAACACCATGCTCCGCTTCTGGTGAACCCATCACCGTGCCATTAAGCCGTAAGCCTCCAGCAGTTAATCCAACCGTTAGA
AGCCCAGCAGCAAGTCAAAGCTCCGAGGTGTTCTTTGACAATTGTCAGAAGTTACTGGATAAATTCAATTACCCATGGGAGTT
GATGCCCCTAATGTATGTGATCTTAGAAAATGCCAACGTAGATATGCAGGAAGCAGCACGAAGGATTGAGGAAGGTCAATGTG
TTGTGAATGAATACACCCGAATGCACAATCTCAATATGTACGATGGAGCCGAACTGCGGGGTTCAACGAGACAATGTGGATGA
TAA
>PpadsxM_CDS [Phlebotomus papatasi] P. papatasi doublesexM CDS
ATGGTGTCGGAGCACAATACGTGGAGCGACGTGATGTCCAATTCGGACCTCACAGACTCGAAAAACGAGTTGTGCGGAGGAGC
CTCGAGCTCCAGCGGCAACTCATTGAGCCCGAGGACGCCACCAAATTGCGCCCGCTGCCGCAATCACGGCCTCAAGATAGCCC
TTAAGGGTCACAAACGCTACTGCAAGTACCGTTACTGCAACTGCGAAAAGTGTCGTCTAACGGCCGAACGTCAGCGGGTGATG
GCCTTGCAGACGGCCCTAAGACGTGCTCAGGCTCAGGACGAGGCACGTCAGATCTCCGCGGATGAGGTGCCACCGACGCCACC
GCCAGCCCCGTCCATTTGCACCCTTGCCACTCCTAAGAGTCAGTGCGATAGCGAACCACCGGCACCCATTACGGTGCCCACAC
CGGCACGTTCCCTAGAGGGTTCCTGCCACTCATCATCGGCCACACCCTGTTCTGCTTCCGGAGAACCCATCTCCGTGCCATCC
AGCCGCAAATTGGCCCCAGTCAATCCAACCATCAGAAATCCAACTGCAAGTCAAAGCTCCGAAGTCTTCTTTGACAATTGCCA
GAAATTGCTGGACAAATTCAATTATCCTTGGGAACTGATGCCCCTAATGTATGTGATCTTGGAAAATGCCCCGGACGTGAGGG
AAGCGGCACGGAGAATTGAAGAAGCTGCGAGTATAATAAAATGTCGATATGGACATGGACTCCCAATCCCAGCACACCAGTAC
TACACATACCTATCTACGGCGGCAGCGAGAGTGCAGACGTCGATATCCATATCCTTCGTACTACTACACATACTGGGCAGCCG
GTATACAAGTCCCTTCCTATATAACTTCTATAACGAGACAGTACTGTCGCCGCATTTGCACAGCAAGTACCTCAATTCGGAAA
GTACTCCTGCGGTACGGAATATATCGGAATCTCCGTCAGCCTATTCATCTCTGCCACTTCCTGCCGCTCTACAAATTACTCGT
CTGATACCAACGAATAGCTTATCTGGTGCATCGACGTAG
>LlodsxM_CDS [Lutzomyia longipalpis] L. longipalpis doublesexM CDS
ATGTCCAATTCGGACCTCACAGACTCGAAGAATGAGTTGTGCGGGGGAGCCTCGAGCTCCAGCGGCAACTCATTGAGCCCGAG
GACGCCGCCAAATTGCGCCCGCTGCCGCAATCACGGCCTCAAGATTGCCCTAAAGGGTCACAAGCGTTACTGCAAATTTCGCT
ACTGCAACTGCGAAAAGTGCCGCCTCACGGCCGAACGACAGCGGGTGATGGCCCTACAGACGGCCCTAAGACGTGCTCAGGCT
CAGGACGAGGCGCGTCAAATCTCCGCGGATGAGGTGCCACCGACACCACCGCCACTTACTGGTCAAATTGCAACAACGCCCAA
AAGTCAGTGCGACGGTGAACTACCGCGATCGAATACAGTGCCAACTCCGGCACGTTCCCTCGAGGGCTCCTGCCACTCATCTT

CGGCAACACCATGCTCCGCTTCTGGTGAACCCATCACCGTGCCATTAAGCCGTAAGCCTCCAGCAGTTAATCCAACCGTTAGA
AGCCCAGCAGCAAGTCAAAGCTCCGAGGTGTTCTTTGACAATTGTCAGAAGTTACTGGATAAATTCAATTACCCATGGGAGTT
GATGCCCCTAATGTATGTGATCTTAGAAAATGCCAACGTAGATATGCAGGAAGCAGCACGAAGGATTGAGGAAGCAATGGGAG
AGTGCAGACGTCGATATCCATATCCTTCGTACTACTACACATACTGGGCAGCCGGTAGTAGTCCCTACCTGTTCAATTACAAC
ATCCTCTCGTCGCAATTGCAAAATAAGAGCTTTAGCTGTGACAGTATTTCATCAACACGTGGCATCTCAGAGGCACCATCGTC
CTACCCCTCAATATCACTACCTGCCGCCGCCCTACAAATCAGACGTCTGATACCAACTGGTAACCTAGCCGGTGCATCGACGT
AG
>LlofruM_par_CDS [Lutzomyia longipalpis] L. longipalpis fruitlessM  partial CDS
ATGATGTCACCACCGAATATGTACGGTGGTCCCTTTCAACCGTATCGGGGACAACCACACTCGTTGGTGTCCCCGCGCGAAGA
CAGCCCACCGACTTCGGTTTTAAACCTCAAGCGCTACACCACCAATGACCCTCCATCATCGCCGACCATACCTGTATCTGCGC
ATCATCACCTGCATCACACACGTGAGCAGGAGACATCGATGGCACATCAGCGACCGCTGTCAGCTCAGTCTCCACAGAGATAC
ACCACTGACCAAGGCAATATGGACCAGCAGTTCTGTCTTCGCTGGAATAATCATCCGACCAACTTGACTGGCGTGCTGACTTC
CTTGCTCCAGNGGGAGGCCCTCTGTGATGTCACACTGGCGTGC
>PdufruM_CDS [Phlebotomus duboscqi] P. duboscqi fruitlessM CDS
ATGATGACTACTCCGGACATCTTCGGCAGCCCGTTTCCGCCGTATAGGTGTCAGCCGTTGCCTGTACTTCCACCACGTGATGA
TAGTCCGCCAATGCCGGCGCTTGACCTTAAACGCTATCCCAGCACTGAGGTCCCACCGGCATCAACAACACCTGTCCGTTTAT
CTGAACGTGAAATGCCGGTGCCGCACCTGCGTCCATCTTCAGCACAAACACCACCTAGATACACCACTGACCAAGGCAATATG
GACCAGCAGTTCTGTCTTCGCTGGAATAATCATCCGACCAACTTGACTGGCGTGCTGACTTCTTTGCTCCAGAGGGAGGCCCT
CTGTGATGTTACATTGGCATGCGATGGTGGTGAAATTGTCAAGGCACATCAGACCATCCTGTCAGCATGCAGTCCATATTTTG
AAAGCATTTTCCTGCAGAACTCTCATCCGCATCCAATAATATATTTGAAAGATGTTAGATACTCAGAGATGCGCTCACTCCTT
GATTTCATGTACAAGGGTGAGGTGAACGTTGGCCAAAGATCGTTACCGACATTTCTAAAGACAGCCGAAAGTTTACAG
>PbefruM_CDS [Phlebotomus bergeroti] P. bergeroti fruitlessM CDS
ATGATGACTACTCCGGACATCTTCGGCAGCCCCTTTCCACCGTATAGGGGTCAGCCGTTGGCTGTACTTCCACCACGTGATGA
CAGTCCGCCAACGCCGGCACTTGACCTCAAACGCTATGCCAACACTGAGGTCCCTCCGGCATCAACAACACCTGTCGCACCAC
ACCTCCGTTTACCGCTGCATCATGCACATGAACGTGAAATGCCGATACCGCACCAGCGTCCATCTTCGGCACAAACACCACCT
AGATACACCACTGACCAAGGCAATATGGACCAGCAGTTCTGTCTTCGCTGGAATAATCATCCGACCAACTTGACTGGCGTGCT
GACTTCTTTGCTCCAGAGGGAGGCCCTCTGTGATGTTACATTGGCATGCGATGGTGGTGAAATTGTCAAGGCACATCAGACCA
TCCTGTCAGCATGCAGTCCATATTTTGAAAGCATTTTCCTGCAGAACTCTCATCCGCATCCAATAATATATTTGAAAGATGTT
AGATACTCAGAGATGCGCTCACTCCTTGATTTCATGTACAAGGGTGAGGTGAACGTTGGCCAAAGATCGTTACCGACATTTCT
AAAGACAGCCGAAAGTTTACAG
>PpafruM_CDS [Phlebotomus papatasi] P. papatasi fruitlessM CDS
ATGATGACTACTCCGGACATCTTCGGCAGCCCCTTTCCGCCGTATAGGTGTCAGCCGTTGGCTGTACTTCCACCACGTGATGA
TAGTCCGCCAATGCCGGCACTTGACCTCAAACGCTATCCCAGCACTGAGGTTCCTCCGGCATCAACAACACCTGTCCGTTTAC
CTGAACGTGAAATGCCGATGCCGCACCTGCGTCCATCTTCGGCACAAACACCACCTAGATACACCACTGACCAAGGCAATATG
GACCAGCAGTTCTGTCTTCGCTGGAATAATCATCCGACCAACTTGACTGGCGTGCTGACTTCTTTGCTCCAGAGGGAGGCCCT
CTGTGATGTTACATTGGCATGCGATGGTGGTGAAATTGTCAAGGCACATCAGACCATCCTGTCAGCATGCAGTCCATATTTTG
AAAGCATTTTCCTGCAGAACTCTCATCCGCATCCAATAATATATTTGAAAGATGTTAGATACTCAGAGATGCGCTCACTCCTT
GATTTCATGTACAAGGGTGAGGTGAACGTTGGCCAAAGATCGTTACCGACATTTCTAAAGACAGCCGAAAGTTTACAGGTACG
CGGTTTGACCGACAACAACAATATCAACTACCGACCAGAGAGCGACAGGGATCGCGATTCAGAAACGAATGCGAGTGGTGCTA
TGAAACATTATGATAAAACAGAACGGGACAGAGATCGCGACCGGGAGCGGCTGGACCGCGATCGTGAGGAGAATTCGGAGAGT
AAAGACCGCGATAGGGAGACACCTGTGGATCACCTGAGCAGCAGCAGCAGGAGCAGCAAGCGGAAACGTGAAAATTCATTAAA
CTGTGATAATTCAATGCGTGGCGCCAGTGTCCAAGAAAGGCATTATTCTCAGGATTCTCAGGCATCGTCGCATAGTAGTTATA
AATCCAGTCCATTGCCAAAATTAAATCCCCTGGAAGGAGAAGACACACGCCGAAATTCACCAGCGTTAAATGCCAGCGGCGCC
AATCAATCAGTTAGCATTAAACAAGAATTACCTGATATGGGCCATCATCCTGGTTTACCACCAGAATTACTTCCGCCCACTTC
AATGTCCCTGCATCCTGAGGATATGACAAGTCTACTTCCGGCTCATGGACTACAAGTGGAGACCTCGGAAAATGACTCGCAGC
ATCCTCAAATGGACCACAGTGATAATATCGATGGGCCGGGGGGGCATCGTCCACCTCCACCACTTCCACCGCCGCCACACCTC
CATCATCATGGACAGCACAGCAGTGGAGACGGAGAGTCAAAGCATCCACTGGTGCCAAGTCGATCGCGTTGTGGCGATGGCGG
CAGCAGTCGCGCCAGTCCCATCATGGGCGTCGTCGCCGCTGCCCTCCAGCACCATCAGCACCACCAGCATCAGATGTCCTACC
ACAATATGTTCTCGCCCACGAGAGAGCTGGCCGGCACCATGTGGCGATGTCGAACGTGCGGCAAGGAGGTCACCAACAGGTGG
CATCACTTCCATTCGCACACAGCCCAACGGAGCATGTGCCCCTACTGCCCGGCCACTTACAGTCGCATCGACACACTACGCTC
CCACCTCAGAGTAAAGCATCCCGATCGTCTGATCAAGAACTAG
